# The GENCODE CLS project: massively expanding the lncRNA catalog through capture long-read RNA sequencing

**DOI:** 10.1101/2024.10.29.620654

**Authors:** Tamara Perteghella, Gazaldeep Kaur, Sílvia Carbonell-Sala, Jose Gonzalez-Martinez, Toby Hunt, Tomasz Mądry, Irwin Jungreis, Fabien Degalez, Carme Arnan, Ramil Nurtdinov, Julien Lagarde, Beatrice Borsari, Cristina Sisu, Yunzhe Jiang, Ruth Bennett, Andrew Berry, Marta Blangiewicz, Daniel Cerdán-Vélez, Kelly Cochran, Covadonga Vara, Claire Davidson, Sarah Donaldson, Cagatay Dursun, Silvia González-López, Sasti Gopal Das, Kathryn Lawrence, Daniel Nachun, Matthew Hardy, Zoe Hollis, Mike Kay, José Carlos Montañés, Pengyu Ni, Emilio Palumbo, Carlos Pulido-Quetglas, Marie-Marthe Suner, Xuezhu Yu, Dingyao Zhang, François Aguet, Kristin Ardlie, Stephen B. Montgomery, Jane E. Loveland, M. Mar Albà, Mark Diekhans, Andrea Tanzer, Jonathan M. Mudge, Paul Flicek, Fergal J Martin, Mark Gerstein, Manolis Kellis, Anshul Kundaje, Benedict Paten, Michael L. Tress, Rory Johnson, Barbara Uszczynska-Ratajczak, Adam Frankish, Roderic Guigó

## Abstract

Accurate and complete gene annotations are indispensable for understanding how genome sequences encode biological functions. For more than twenty years, the GENCODE consortium has developed reference annotations for the human and mouse genomes, becoming a foundation for biomedical and genomics communities worldwide. Nevertheless, collections of important yet poorly-understood gene classes like long non-coding RNAs (lncRNAs) remain incomplete and scattered across multiple, uncoordinated catalogs. To address this, GENCODE has undertaken the most comprehensive lncRNA annotation effort to date. This is founded on the manually supervised computational annotation of full-length targeted long-read sequencing, on matched embryonic and adult tissues, of orthologous regions in human and mouse. Altogether 17,931 human genes (140,268 transcripts) and 22,784 mouse genes (136,169 transcripts) have been added to the GENCODE catalog representing a 2-fold and 6-fold growth in transcripts, respectively - the greatest increase in the number of annotated human genes since the sequencing of the human genome. Our targeted design assigned human-mouse orthologs at a rate beyond previous studies, tripling the number of human disease-associated lncRNAs that have mouse orthologs. Novel lncRNA genes consistently exhibit biological signals of functionality, and they greatly enhance the functional interpretability of the human genome. While poorly expressed in bulk RNA-Seq samples, many of them are highly expressed in specific cell populations, maybe even contributing to cell-type determination. The expanded GENCODE lncRNA annotations mark a critical step toward deciphering the human and mouse genomes.

## Introduction

Genome sequencing has enormously impacted our understanding of human biology and disease. However, genome sequences are of little value in absence of a reliable map of genes and transcripts, without which it is impossible to understand genome function and to interpret the effect of genetic variation on phenotypes. Shortly after the publication of the first human genome drafts twenty-five years ago^1,2^, the ENCyclopedia Of DNA Elements (ENCODE) project^3^ was initiated to identify all elements in the human genome that confer biological function. As part of this broader effort, GENCODE was launched to produce a definitive catalog of all human genes and transcripts^4^. Over the years, GENCODE^4–9^, an international partnership of manual annotation, computational biology, and experimental groups has established itself, together with RefSeq^10–13^, as the primary reference gene annotation catalog for human and mouse.

While GENCODE and RefSeq are now in close agreement on the ‘canonical’ set of protein-coding genes^14,15^, lncRNA annotations are substantially less mature. A number of alternative lncRNA catalogs have been produced; these are typically based on short-read RNA-Seq data, which are not used by GENCODE due to concerns on the fidelity of the transcript models obtained^16–19^. As a result, and despite efforts such as RNA Central^20^, downstream users are faced with a fragmented landscape of non-interoperable, inconsistent and sometimes redundant lncRNA annotations, many of which are also structurally incomplete. This slows progress in the field, especially when it comes to understanding the biological relevance of lncRNAs.

Here, we report the GENCODE efforts to produce a unified reference lncRNA catalog in the human and mouse genomes. Toward that aim, we specifically employ the Capture Long-read Sequencing (CLS) strategy^21,22^ to enrich transcripts originating from targeted genomic regions. We designed a very large capture array, with orthologous probes in the human and mouse genomes, targeting different non-GENCODE lncRNA annotations and additional regions suspected of unannotated transcriptional activity. We employ a cDNA library preparation protocol, CapTrap-Seq^23^, that enriches for 5’ to 3’ complete RNA molecules. We used this approach in a matched collection of adult and embryonic tissues, in human and mouse, selected to maximize transcriptome complexity. The resulting libraries were sequenced pre- and post-capture using PacBio and Oxford Nanopore (ONT) long-read sequencing technologies, as well as Illumina for short-reads.

The manually supervised computational annotation of this data has led so far to the inclusion of 17,931 additional human genes (140,268 transcripts) and 22,784 additional mouse genes (136,169 transcripts) in GENCODE, most of which were identified only post-capture. This is, by far, the largest addition to the GENCODE catalogs since the first drafts of the human and mouse genomes, and it constitutes a substantial advance toward a complete gene catalog for these species. The mapping of these novel lncRNA transcripts into other genomes has enabled the prediction of hundreds of thousands of transcripts across primates.

Novel lncRNA genes consistently exhibit biological signals similar or even stronger than those of previously annotated lncRNAs and, in some cases, even comparable to those of protein-coding genes. They greatly enhance the functional interpretability of the human genome, as we can assign millions of previously “orphan” omics measurements to “bona fide” well-constructed transcriptional units, as well as tens of thousands of “orphan” genetic variants associated with phenotypes. They are associated with disease phenotypes, and, while poorly expressed and difficult to detect in bulk RNA-Seq samples, they are often highly expressed in specific cell populations, and some may even contribute to the definition of cell subtypes.

The work described here marks a transformative leap in our ability to chart and interpret the non-coding genome. By dramatically expanding and refining the catalog of lncRNAs in human and mouse, GENCODE’s efforts offer a critical foundation for future studies into gene regulation, disease mechanisms, and cellular diversity, illuminating vast regions of the so-called dark genome.

## Results

### Targeting and sequencing the long non-coding transcriptome with CapTrap-CLS

We designed a capture array targeting a large fraction of the putative non-coding transcriptome, including the major non-GENCODE lncRNA annotations^16,18,19,21,24–27^, as well as small non-coding RNAs, enhancers^28^, evolutionarily conserved RNA structures^29,30^, regions that host non-coding GWAS^31,32^, that show putative protein-coding sequence conservation^33^, or that are ultraconserved^34^. Probes were designed in the human genome version GRCh38 using GENCODE v27 as reference annotation (**Figure 1A, Table S1**). Orthologous regions of the mouse genome were targeted in a corresponding mouse capture library (GENCODE vM16 on GRCm38.p6, **Table S1**). In total, we targeted 176,435 human regions, covering 2.9% of the genome, and 148,965 regions in mouse, covering 2.8% of the genome.

**Figure 1.**
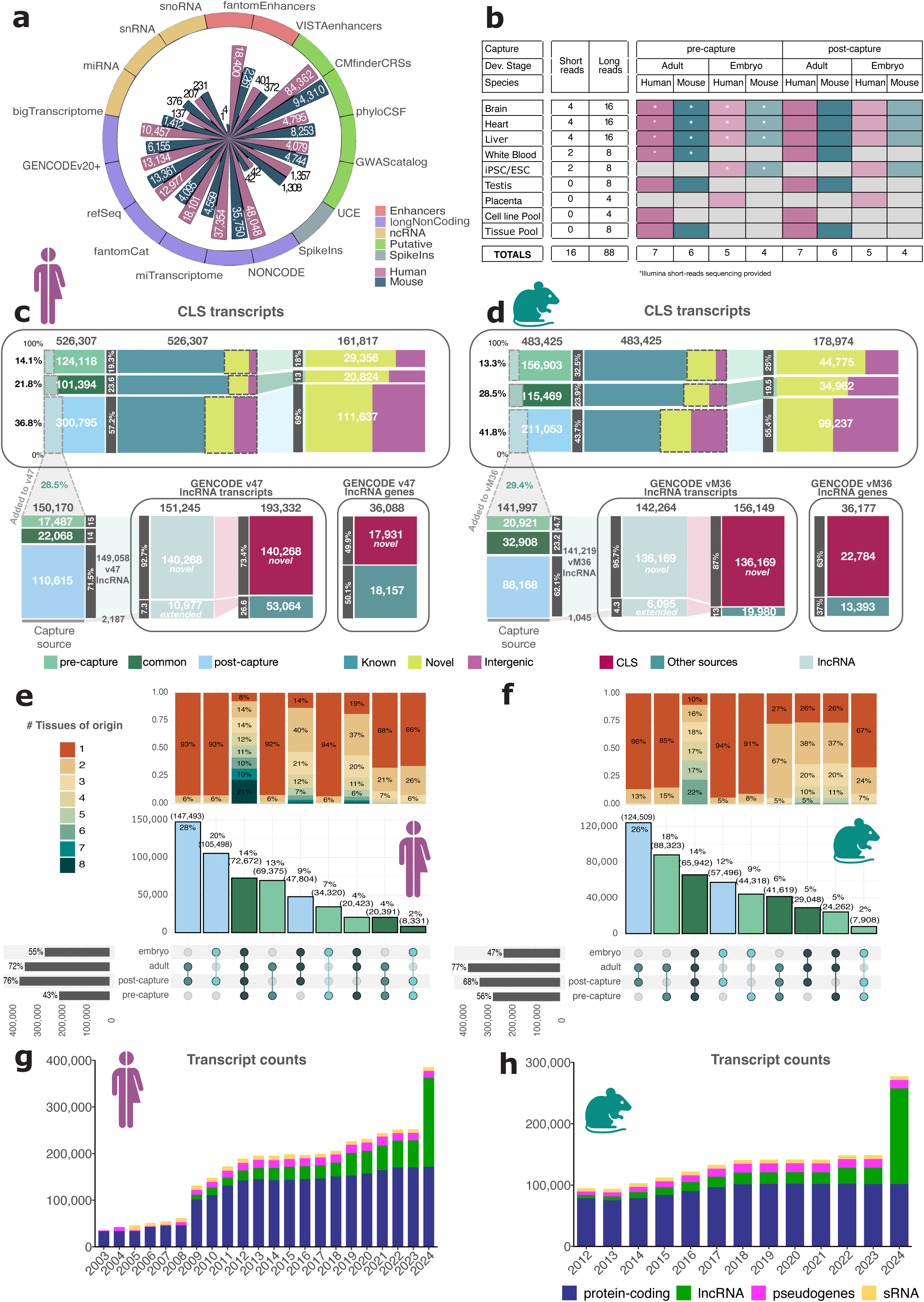
Targeting and sequencing the long non-coding transcriptome with CapTrap-CLS. **A)** The capture panel; each bar reports the number of targeted regions per annotation catalog, for human and mouse, grouped by annotation class. **B)** Samples used for CapTrap-CLS. Samples from matched adult and embryonic tissues from human and mouse were sequenced using PacBio and Oxford Nanopore Technologies (ONT) long-read platforms. Short-reads were sequenced with Illumina and highlighted by an asterisk when available. **C) and D)** Summary of the initial set of CLS transcripts and of the set finally included in GENCODE for human (C) and mouse (D). Top panel, CLS transcripts are categorized based on the novelty status with respect to GENCODE v27 (human) and vM16 (mouse). Bottom panel: CLS transcripts added to GENCODE v47 (human) and vM36 (mouse). The number of CLS models labelled as “Added to v47” is slightly smaller than the number of novel CLS transcripts in v47 because ∼1% of the v47/vM36 lncRNA were created via older versions of the CLS data, and hence have an untrackable CLS capture source. See **Figure S3** for a more detailed description. **E) and F)** The panels show the yield of CLS transcripts in human (E) and mouse (F) across developmental stages and capture protocol. The barplot on the left shows the number of CLS transcripts detected (from bottom to top) pre-capture, post-capture, as well as from adult and embryonic samples (percentage computed over the totality of the transcripts generated). The upset plot shows the intersections across these categories; the dots are colored according to the developmental stage of origin (whether adult, embryo or both), while the bars display the overlap between pre-capture and post-capture experiments. The barplot above shows the proportion of shared transcripts across tissues. **G) and H)** GENCODE annotation history. Numbers of transcripts on primary assembly chromosomes in every year’s last GENCODE release in human (G) and mouse (H) broken down by broad biotype. IG/TR genes excluded.

We prepared CapTrap-Seq libraries from a matched collection of adult and embryonic tissues (brain, liver, and heart), in human and mouse, plus samples designed to maximize transcriptome complexity (white blood, testis and placenta, iPSC/ESC cell lines, as well as pools of adult tissues and cell lines, **Table S2**). Most samples were sequenced pre- and post-capture using three different technologies: Illumina, PacBio, and ONT. In total, we produced 104 data sets (16 short-read, 438 million reads, and 88 long-read, 736 million reads **Figure 1B**, https://guigolab.github.io/CLS3_GENCODE/summary_GENCODE.html). The effectiveness of the capture protocol is indicated by the enrichment of reads originating from the targeted regions in the post-capture samples (from 3.7x to 36x depending on the species, tissue, and platform), as well as by the enrichment of ERCC spike-ins^35^ used as controls (**Supplementary Materials, Figure S1**).

We generated transcript models from long RNA-Seq reads using LyRic^36^, an in-house framework specifically designed for CLS data (**Supplementary Materials**), and that compared favourably to other methods for the task of confidently identifying novel transcripts^37^. These were built for each sample separately, then merged across tissues and stages to produce a comprehensive set of CLS models **(Figure S2)**.

Across all samples, ignoring variations in the termini, we generated 526,307 CLS transcript models in human and 483,425 in mouse **(Figure 1C,D, S3AB, S4AB, Supplementary Materials)**. Of these, 161,817 (31%) were novel in human and 178,974 (37%) in mouse (with respect to GENCODE versions v27 and vM16, **Figure S3CD**, **Table S3**). These were predominantly detected in post-capture samples (82% in human and 75% in mouse, **Figure 1C-F**, **Figure S3B**), with yield varying across tissues **(Figure 1E-F, Figure S3B**), and across target regions **(Figure S5**, **S6)**.

Most sequence and structural features of the novel CLS models are similar to those of lncRNAs previously annotated in GENCODE **(Figure S7, Table S4).** They are, however, more tissue-specific than annotated loci: 83% of human and 72% of mouse CLS transcripts are detected in just one sample (compared with 66% and 62% of known loci, respectively). CLS transcripts detected both in adult and embryo tissues are the most broadly expressed **(Supplementary Materials, Figure S8).**

### Incorporating CLS models into the GENCODE annotation

The set of CLS models was used as input by the HAVANA team of expert manual annotators to produce GENCODE releases v47 and vM36. Manual curation is a founding principle of GENCODE, vital for ensuring the stringency required for ‘reference quality’ gene annotation. Since the original ‘gene by gene’ approach is not scalable to the one million CLS models produced here, we deployed a new computational workflow named TAGENE that is manually supervised at key steps, and was developed iteratively via extensive testing by the annotators **(Supplementary Materials)**. We specifically chose parameters to achieve a minimal false positive rate (i.e., the creation of false annotations) at the expense of an elevated false negative rate (i.e., the rejection of true annotations). Thus, as deployed here, TAGENE eliminates all CLS models antisense to protein-coding genes or located entirely within the genomic bounds of pseudogenes, and employs a strict intronic filtering step based on short-read data for the remainder (**Supplementary Materials, Figure S9**).

Using these stringent filters, and additional methodological adjustments, we defined a set of CLS transcripts to be included in GENCODE. Thus, we annotated 140,268 and 17,931 novel lncRNA transcripts and genes in human, and 136,169 and 22,784 in mouse (with respect to v27 and vM16 respectively), the vast majority incorporated as of releases v47 and vM36 **(Table S5)**. This represents a substantial increase over the number of lncRNA annotations present in GENCODE v46 (>2x) and vM35 (6x, **Figure 1C,D,G,H, Figure S10**), and brings, for the first time, the number of mouse lncRNA loci (36,177) on a par with human (36,088). Collectively, these new annotations increased the transcriptional footprint of GENCODE annotations on the genome by 27.7Mb and 28.9Mb respectively (compared to v46 and vM35, **Table S6)**. CLS models that were not incorporated into GENCODE in this initial round of annotation may yet be incorporated into future releases as this workflow matures.

### Unified lncRNA catalog

A central goal of this work was to advance toward a complete catalog of human and mouse lncRNAs. By means of our capture design, we doubled the proportion of lncRNAs from non-GENCODE catalogs from 15% in version v27 to 29% in v47 **(Figure 2A, Figure S11AB**), improving the quality of GENCODE lncRNAs across various metrics^19^ **(Figure S11CD)**. Despite the inclusion of many novel CLS transcripts, a significant number of lncRNA genes from non-GENCODE catalogs remain absent (77,835 do not overlap annotated loci in v47, **Figure S12A)**. While expanding the range of tissues that we monitored here is likely to bring in additional candidates, lncRNAs not incorporated into GENCODE have generally weaker splice site support **(Figure S11AB)** and are more catalog-specific than those identified through the CLS approach **(Figure S12B)**. Despite their large aggregate size, a large fraction of novel lncRNA genes in v47 (46%) do not originate from existing lncRNA catalogs **(Figure S12C-F)**, underlying the discovery potential of the CLS strategy.

**Figure 2.**
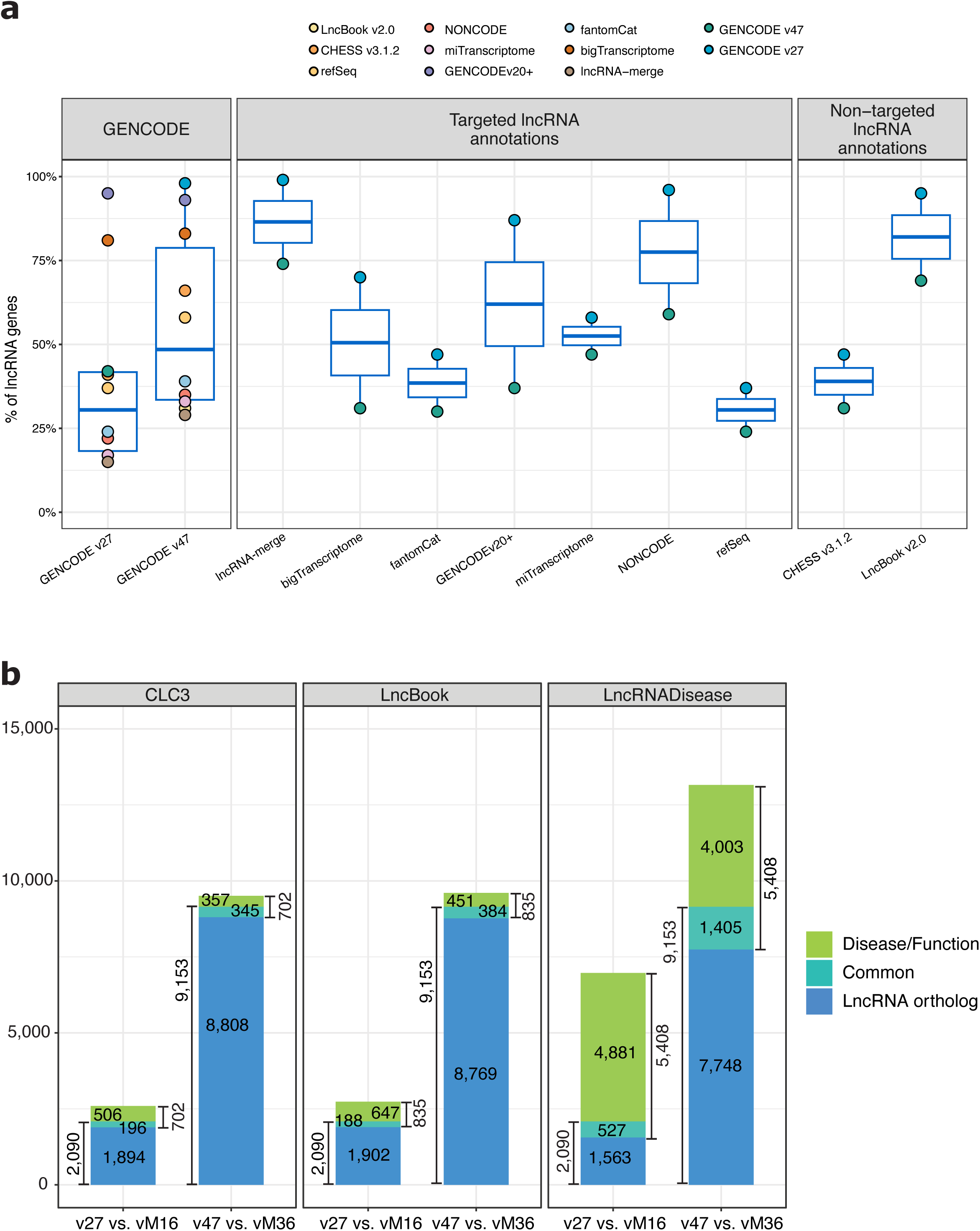
Expansion of the GENCODE lncRNA annotation compared to other lncRNA catalogs. **A)** Gene-level overlap across annotations. The values correspond to the percentage of gene loci from the catalogs (x-axis) that overlap the different annotations (represented in the box-plot). For instance, 29% of the lncRNAs in the merge of all catalogs (lncRNA-merge) are included in GENCODE v47. Conversely, 74% of the lncRNAs in v47 are included in lncRNA-merge. Overlap is defined as a complete overlap of the gene body; both spliced and monoexonic genes are included in this analysis. See also **Figure S12A**. **B)** Venn diagrams illustrating the human–mouse lncRNA orthologs corresponding to clinically relevant human lncRNAs, across three independent databases. For instance, of the 9,153 human lncRNAs with mouse orthologs in v47, 1,405 are included in the set of 5,408 disease genes in LncRNADisease.

### An orthology map of lncRNAs between human and mouse

Assigning gene orthology between species is crucial for understanding their evolutionary history and, in the case of human, for developing animal models of diseases. LncRNAs are particularly challenging in this regard, due to their lack of sequence conservation^38,39^. The CLS capture libraries addressed this limitation by targeting orthologous genomic regions between human and mouse. Coupled with sampling equivalent tissues from both species, this approach greatly improved the detection of transcripts from orthologous loci. Using a stringent reciprocal orthology framework implemented in ConnectOR^40^, we identified 9,153 human lncRNA loci orthologous to 9,142 mouse loci, representing approximately 25% of the annotated lncRNAs in each species. This marks a significant increase compared to earlier GENCODE versions (v27 and vM16), which reported only 13% and 15% orthologous lncRNAs **(Figure S13AB)**. As a result, we substantially increased the identification of clinically relevant counterparts of human lncRNAs in the mouse genome **(Figure 2B)**.

### Protein-coding genes and pseudogenes

Although this study focuses on lncRNAs, 84,149 novel human CLS models map to 11,944 known protein-coding genes. The equivalent numbers for mouse are 97,744 CLS models and 13,886 coding genes. So far, 9,454 high confidence protein-coding CLS-derived transcripts have been added to GENCODE v49, with many more transcripts likely to be added in future. In addition, analyses of large-scale tissue proteomics data and evolutionary conservation of putative ORFs led to the conservative discovery of eight novel human and 24 novel mouse protein-coding genes **(Supplementary Materials, Figure S14, S15)**. Similarly, although we did not focus on pseudogenes, a number of probed regions overlapped them; we found that our capture strategy enhanced the sensitivity in recording expression changes in pseudogenes compared to parent genes **(Supplementary Materials, Figure S16, Table S7**).

### Novel lncRNAs advance the functional interpretation of the human genome

Here we show how the GENCODE annotation, extended by incorporation of the CLS data, greatly enhances the functional interpretability of the human genome. We compare novel CLS models (that is, models in v47 not in v27) against previously annotated lncRNAs and protein-coding genes in v27. For most analyses we employ a set of decoy models that attempt to mimic the background (non-genic) behavior of the genome (see Methods). When the biological signal monitored can be originating from overlapping annotated genes, we restrict our analyses to novel intergenic lncRNAs (novel lincRNAs). We show that novel CLS models account for millions of previously unassigned omics signals, far exceeding what would be expected from the mere increase in annotated transcribed regions. Our analyses also serve to validate the biological relevance of the expanded annotation, showing that the novel lncRNA transcripts have biological signals usually much stronger than the background genome, similar or stronger than those in previously annotated lncRNAs and, in some cases, even comparable to those in protein-coding genes.

### Transcription initiation

In total, we predicted 80,284 novel TSSs in the human genome, 36.2% of which either overlap with CAGE clusters from FANTOM5^41,42^ or are associated with ProCapNet machine-learning predictions^43^, a percentage much larger than for decoys (2.4%). In total we could attach novel TSSs to 10,715 orphan CAGE tags (i.e., tags that could not be associated with previously known TSSs, 7.2% of all orphan tags). The CAGE support for annotated lncRNAs and protein-coding genes is much larger than for novel CLS models, as expected given that these latter are mostly seen only post-capture and more difficult to detect in bulk RNA-Seq samples. Remarkably, the ProCapNet support, which does not directly depend on available transcriptional data, is comparable for CLS, lncRNAs and protein-coding genes **(Figure 3A).**

**Figure 3.**
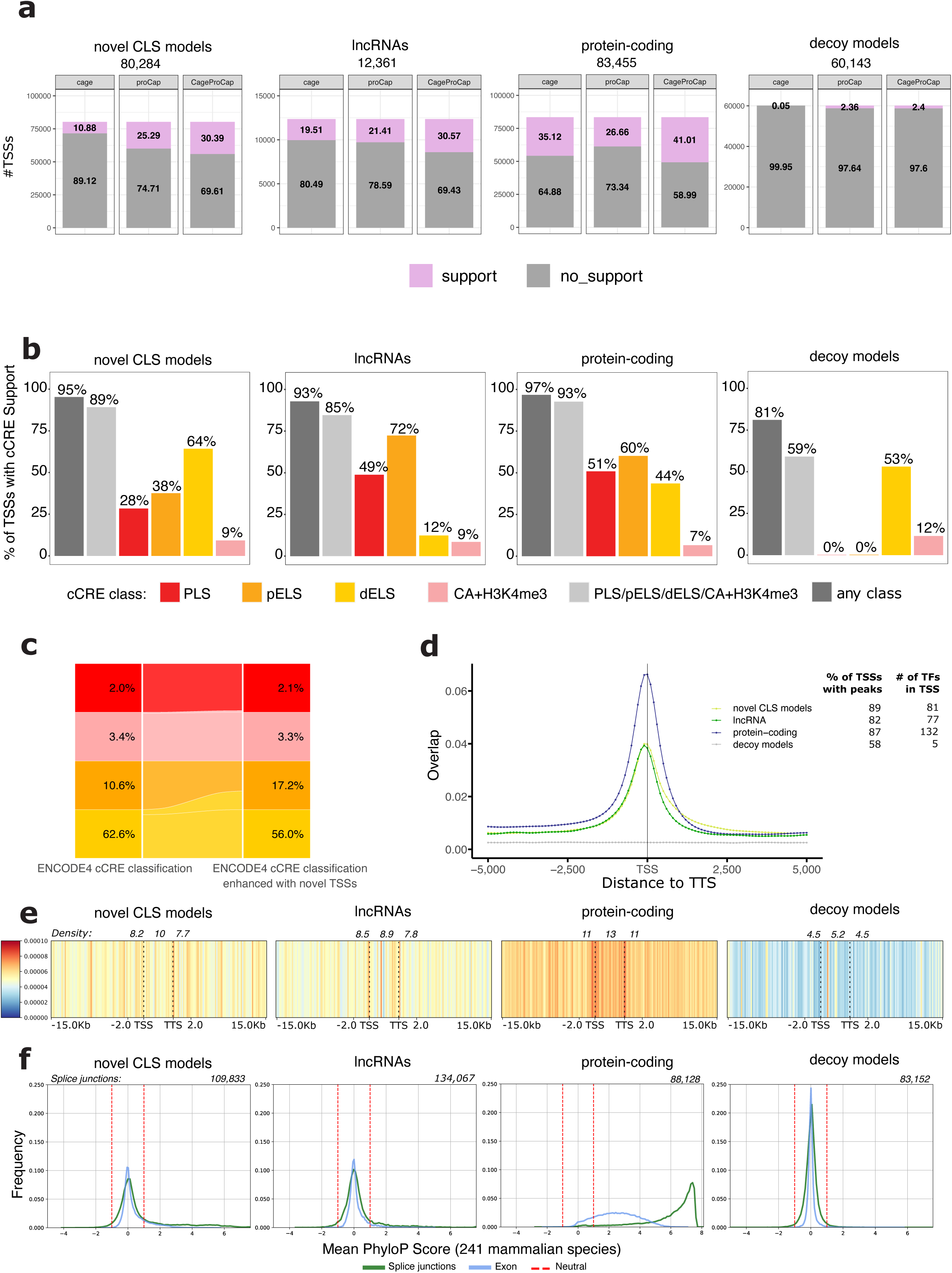
Enhancing the functional interpretability of the human genome. The figure shows how the incorporation of CLS data greatly enhances the functional interpretability of omics measurements on the human genome, assessed on, from left to right, *i)* novel CLS models, *ii)* annotated lncRNA as of GENCODE v27, *iii)* annotated protein-coding genes as of GENCODE v27, and *iv)* decoy models (to simulate background signal). **A)** Proportion of supported Transcription Start Sites (TSSs) within each set using CAGE clusters, proCapNet predictions and both. **B)** Proportion of TSSs supported by different types of cCREs. TSSs with cCRE support are considered those for which the distance between the TSS and the center of the cCRE is < 2kb. The tag “any class” includes additional types of cCREs not shown (CA-CTCF, CA-TF, CA, TF). **C)** Alluvial diagram showing the re-classification of TSS-proximity-dependent cCRE categories in the ENCODE registry upon inclusion of novel TSSs in the expanded annotation. Two pairs of categories are shown *i)* PLS versus H3K4me3 marking in accessible regions (CA-H3K4me3), and *ii)* pELS versus dELS, which share the same histone marking signature, but rely on different proximities to closest TSS (200bp and 2kb, respectively). The percentages indicate the proportion of cCREs from the entire registry that belong to each category in the original classification (on the left) and upon enhancement with novel TSSs (right). **D)** Peaks of transcription factor binding centered on TSS. The plot shows the average (across 1,800 TFs) coverage by ChIP-Atlas peaks of each consecutive 500bp window around the TSS. The coverage increases while we approach the TSS in all cases but for decoys. **E)** GWAS density profile along the gene body and the surrounding ±15kb areas. The exonic density in novel lincRNAs is larger than for decoys (p-value 6.5e-28), comparable to the one of annotated lncRNAs (p-value 0.094), and lower than for protein-coding exons (p-value 0). The numbers above each plot indicate the density of GWAS in the exon, and in the ±2kb proximal regions. **F)** Frequency of per-transcript exon and splice-junctions mean PhyloP scores. The dashed red lines indicate the range considered under neutral selection. On top of each plot the total number of splice-junctions evaluated.

### Chromatin structure

Histone modifications are assumed to play an important role in the regulation of gene expression. The ENCODE consortium recently generated an updated list of 2,348,854 candidate *cis*-regulatory elements (cCREs)^44^ in the human genome after integrating data from 5,712 experiments (which, in addition to ChIP-Seq of histone modifications, include DNase-Seq, ATAC-Seq, and ChIP-Seq of transcription factors)^45,46^. We found that about 89% of all novel TSSs were supported by at least one cCRE, a proportion similar to protein-coding genes and annotated lncRNAs, but larger than in decoy models (59%) **(Figure 3B, S17AB, Supplementary Materials)**. As the ENCODE cCRE classification depends on the proximity to TSSs, we re-classified cCREs in the ENCODE registry taking into account our enhanced collection of TSSs. As a result, more than 153,000 cCREs previously classified as distal dELS (about 10% of them) were re-classified as proximal pELS, bringing down the proportion of dELS in the human genome from 63% to 56% and increasing, conversely, the proportion of proximal pELS from 11% to 17% **(Figure 3C, S17C).**

### Transcription factor binding

Binding of transcription factors (TFs) triggers initiation of transcription. The ChIP-Atlas database^47^ collects ChIP-Seq data for 1,806 human TFs across 30,119 experiments. We merged ChIP-Seq peaks for the same TF, producing a comprehensive set of 65,155,727 peaks along the human genome. We found 89% of the CLS TSSs covered by at least one such peak, a number much larger than for decoys (58%). Similarly, we found overlapping peaks for an average of 81 different TFs for CLS TSSs, but only for five in decoys. We additionally computed the fraction covered by ChIP-Atlas peaks of a 500bp sliding window running -5kb to +5kbp from each TSS. The density profile aggregated over all TFs, increasing as we approach the TSS, is almost identical for CLS and lncRNAs, and somehow weaker than for protein-coding genes. The profile is flat for decoy models **(Figure 3D)**. Overall, the novel CLS models help to assign to promoter regions 3,836,462 ChIP-Atlas peaks previously mapping to intergenic regions (7.1%, **Table S8**).

### GWAS hits

GWAS allows us to connect variants in the genome with phenotypic traits. By identifying the genes affected by these variants we can hypothesize the molecular mechanisms underlying the traits. We computed the density of GWAS hits from the GWAS catalog^32^ within the boundaries of genes and their regulatory regions **(Figure S18**). Overall, we found a density of 10 GWAS hits/100kb within the novel lincRNA exons, twice that for decoys (5.2) and comparable to that of annotated lncRNAs and protein-coding genes **(Figure 3E)**. A similar trend can be observed in their proximal regulatory regions and distal enhancers^48^. The GWAS density decays with the distance to the gene body for novel lincRNAs, annotated lncRNAs and protein-coding genes, in contrast to the flat profile observed for decoy models **(Figure 3E)**. Overall, the percentage of orphan GWAS in the human genome went down from 31% (v27) to 18% (v47), with 19,484 hits mapping now within the boundaries of novel lincRNAs and their proximal regulatory regions.

### Sequence conservation

The conservation of lncRNA sequences is generally low, while splice-junctions often show significant conservation^49^. We evaluated it using the Zoonomia 241-way mammalian genomic alignment^50,51^, measuring conservation via mean per-transcript PhyloP scores^52^ for exon sequences and splice-junctions. Decoy models provide a neutral evolution baseline and define the PhyloP scores between -1.0 and 1.0 as neutral for sequence conservation (**Figure 3F, Table S9**). This results in 97% of the decoy exons being classified as neutral, with 2% showing sequence conservation. GENCODE protein-coding transcripts show the expected high levels of mammalian conservation, with 84% of exons and 97% of the splice-junctions conserved. Novel CLS transcripts show weaker conservation (16% and 26% respectively), although much higher than decoys, and higher than previously annotated lncRNAs (6% and 14%, respectively, **Figure 3F)**.

### Small RNA precursors

Many classes of small RNAs are processed from long RNAs. For instance, 1,244 (67%) of the 1,869 human miRNAs annotated in GENCODE v27 are contained within annotated long RNAs in the same genomic orientation, which could potentially be their precursors. Here, we identified an additional 163 orphan miRNAs contained within novel lncRNA transcripts, bringing the proportion of human miRNAs with candidate precursors to 75% **(Table S10)**. Notably, the host genes of 35 miRNAs are novel lincRNAs **(Figure 4A)**. As an example, for the orphan microRNA cluster MIR513 comprising MIR513a1, MIR513A2, MIR513B and MIR513C, we identified the putative host gene (ENSG00000296782, antisense to a previously known lncRNA). Members of the MIR513 family play a role in various diseases, including different neoplasms and carcinomas, and have been found in healthy tissue mostly in testis. This specific spatiotemporal expression pattern is consistent in the potential host gene (**Table S11**). Small nucleolar RNAs (snoRNAs) are another class of small RNAs processed from long precursors; we increased the number of human snoRNAs with host genes from 572 (61%) to 639 (68%) (**Table S12**). Similar observations can be made in mouse (**Table S12**). We found that splice-junctions of host lncRNAs, in particular novel lncRNA transcripts, exhibit much higher conservation than lncRNAs in general (39% of exons and 74% of splice-junctions conserved, **Figure S19**).

**Figure 4.**
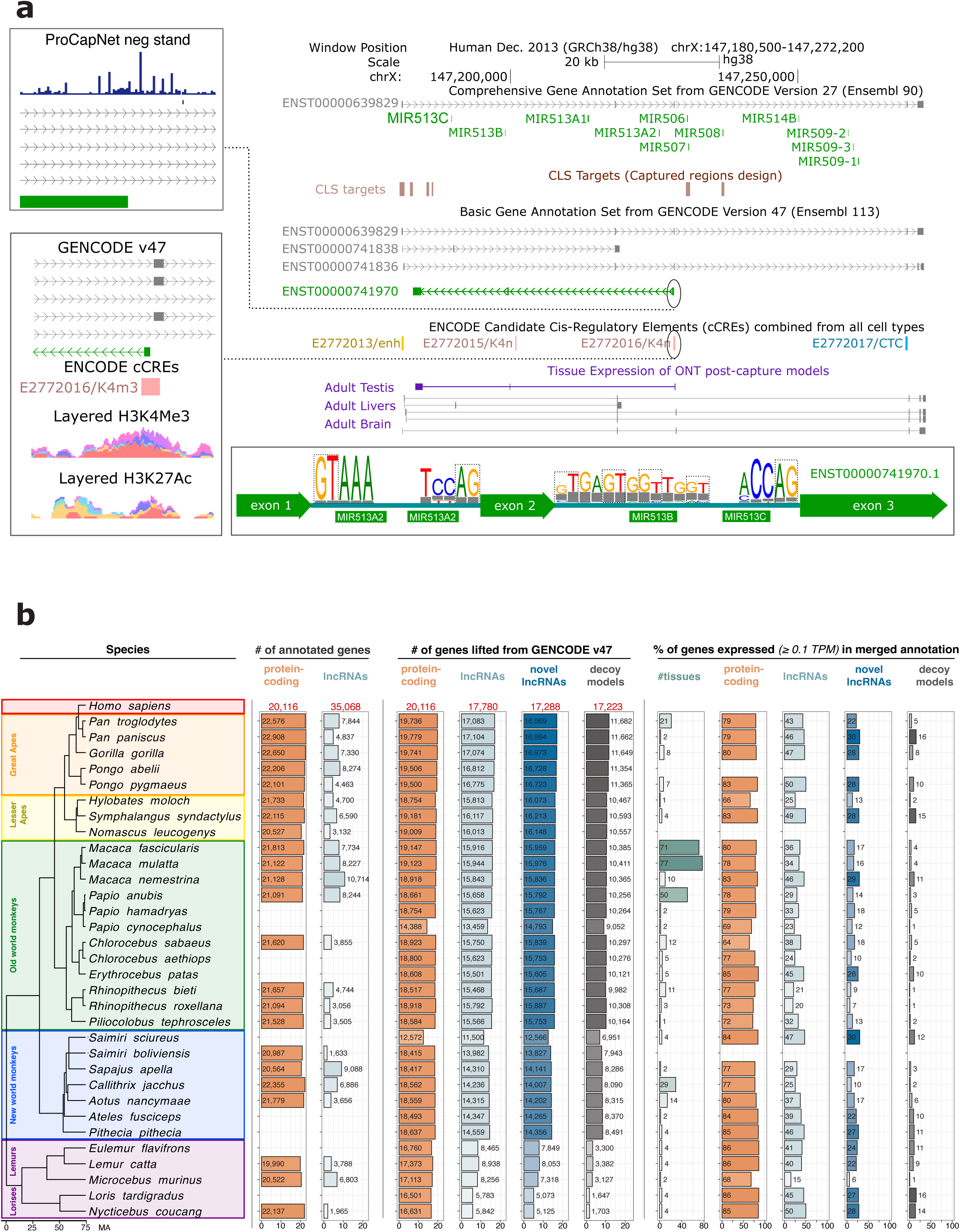
**A)** Example of a putative novel miRNA host gene. The orphan microRNA cluster MIR513 (green transcripts in the track GENCODE v27) resides within a novel lncRNA (ENST00000741970.1, green transcript in the track GENCODE v47 comprehensive), identified targeting a conserved RNA structural element (track CLS long-read RNAs, subtrack CLS targets). This lncRNA is located on the reverse strand, in the same sense as the MIR513 cluster and antisense to another lncRNA (ENST00000639829.1, grey transcript in the tracks GENCODE v27 and GENCODE v47). A cCRE (track ENCODE cCREs) at the 5′ end suggests transcriptional regulation via histone modifications H3K4me3 and H3K27Ac (panel at the bottom left). ProCapNet predictions also supports a TSS at that site (panel at the top left). The panel at the bottom displays splice-junctions conservation for the two introns of ENST00000741970.1 across primate assemblies in the 470-way genomic alignment (track Hiller Lab 470 Mammals). Sequence logos^83^ show the base distribution in the alignment, with gray bars indicating assemblies where no alignment is present. Conservation of the canonical splice site motif GU-AG deep into the superfamily Euarchontoglires. The MIR513 members themselves are derived from MER19C transposable elements, which date back to the root of Euarchontoglires and suggest that this small RNA - host gene relationship most likely emerged around 90 million years ago. All tracks shown are standard UCSC Genome Browser tracks. Remapping the RNA Tissue Atlas^48^ to GENCODE v47 reveals significant expression of this novel host gene (ENSG00000296782.1 / ENST00000741970.1) in only two tissues, YTS cell lines, and melanoma (**Table S11**). In contrast, the expression of the antisense lncRNA gene (ENSG00000284377.3) shows a different expression pattern. **B)** Summary statistics across 32 primate species, representing five phylogenetic groups. From left to right, the column display the *i)* number of protein-coding genes and lncRNAs annotated in RefSeq; *ii)* number of protein-coding genes, lncRNAs, novel lncRNA as a separate category, and decoy models successfully lifted from GENCODE v47 (minimum coverage ≥ 0.5); *iii)* number of tissues with available short-read data and percentage of genes expressed (≥ 0.1 TPM) in at least one tissue, for each biotype.

### Non-canonical translation

Similarly, lncRNAs are known to host small, translated non-canonical ORFs (ncORFs) as assessed by the use of Ribo-Seq data^53^. We used datasets from three human tissues (brain, liver and testis^54^) and identified 44,425 ncORFs with translation signatures in human and 53,629 ncORFs in mouse. Overall, 18% of the novel lncRNAs transcripts contained one or more translated ncORFs, compared to 0.02% of the decoys (**Figure S20A**). We found similar results in mouse (**Figure S20B**).

### Using the CLS transcripts in the lncRNA annotation of primate genomes

While GENCODE focuses on annotating human and mouse genomes, projects such as Ensembl often utilise models from one species to improve the annotation of another via transcript to genome mapping. This process can work well for closely related species, and is especially useful when there is a relative paucity of data to build annotations *de novo* in that species. Thus, we projected annotations from human GENCODE v47 onto 32 primate genomes, spanning between ∼6.4 (chimpanzees) and ∼73.8 (lemurs and lorises) million years of divergence from human (**Figure 4B, Table S13**). As expected, both the number and the sequence coverage of lifted lncRNAs decreased with phylogenetic distance, with a breaking point at ∼40 million years, corresponding to the split from the prosimians (**Figure S21AB**). With a coverage (proportion of the gene span successfully mapped) of more than 50%, we successfully projected 98% of GENCODE novel lncRNAs in chimpanzees and 42% in lemurs and lorises, yielding an average recovery rate of 82% across all primates. For comparison, 91% of protein-coding genes, 80% of annotated lncRNAs, and 51% of decoys were retained. Altogether, this process added over ∼750,000 predicted genes across the primate genomes (around 23,000 on average per species, **Table S14**). To assess whether these expanded annotations correspond to transcribed regions, we mapped short-read RNA-Seq data from 2,470 samples (161 projects) across 29 primate species (**Table S15**). We detected expression of a median 18% of novel lncRNA transcripts per tissue per species compared to 5% of decoys. The link between gene projection and orthology across human, mouse and primates has been also investigated (**Figure S22**).

### Expression patterns and regulation of novel lncRNAs across tissues, cell-types and diseases

In this section, we show, in specific cases, that novel lincRNAs are associated with phenotypes at different levels. First, with molecular phenotypes under genetic regulation. Second, with organismic phenotypes of disease. Third, with cellular phenotypes, where they contribute to uncover defined cell populations which would not be evident without them.

### Genetic regulation of novel lncRNAs

To investigate the extent to which novel lincRNAs’ expression and splicing are affected by common genetic variation, we requantified 298 GTEx caudate nucleus brain samples^55^ with GENCODE v47 as a reference annotation and searched for expression quantitative trait loci (eQTLs). Overall, 1,427 novel lincRNAs have sufficient expression levels for eQTL mapping (**Figure S23**). We found 574 novel lincRNAs as eGenes (i.e., genes with at least one genome-wide significant eQTL **Figure 5A**); further, 17% of lead eSNPs for these eGenes were not significantly associated (nominal p-value > 0.05) with any other previously annotated gene in a 1Mb region (**Figure 5B**). We also performed similar analyses on splicing quantitative trait loci (sQTLs, **Figure 5B**). These findings suggest that many novel lincRNAs are under distinct genetic regulation, implicating them as previously unrecognized contributors to gene expression and splicing variation in the human brain.

**Figure 5.**
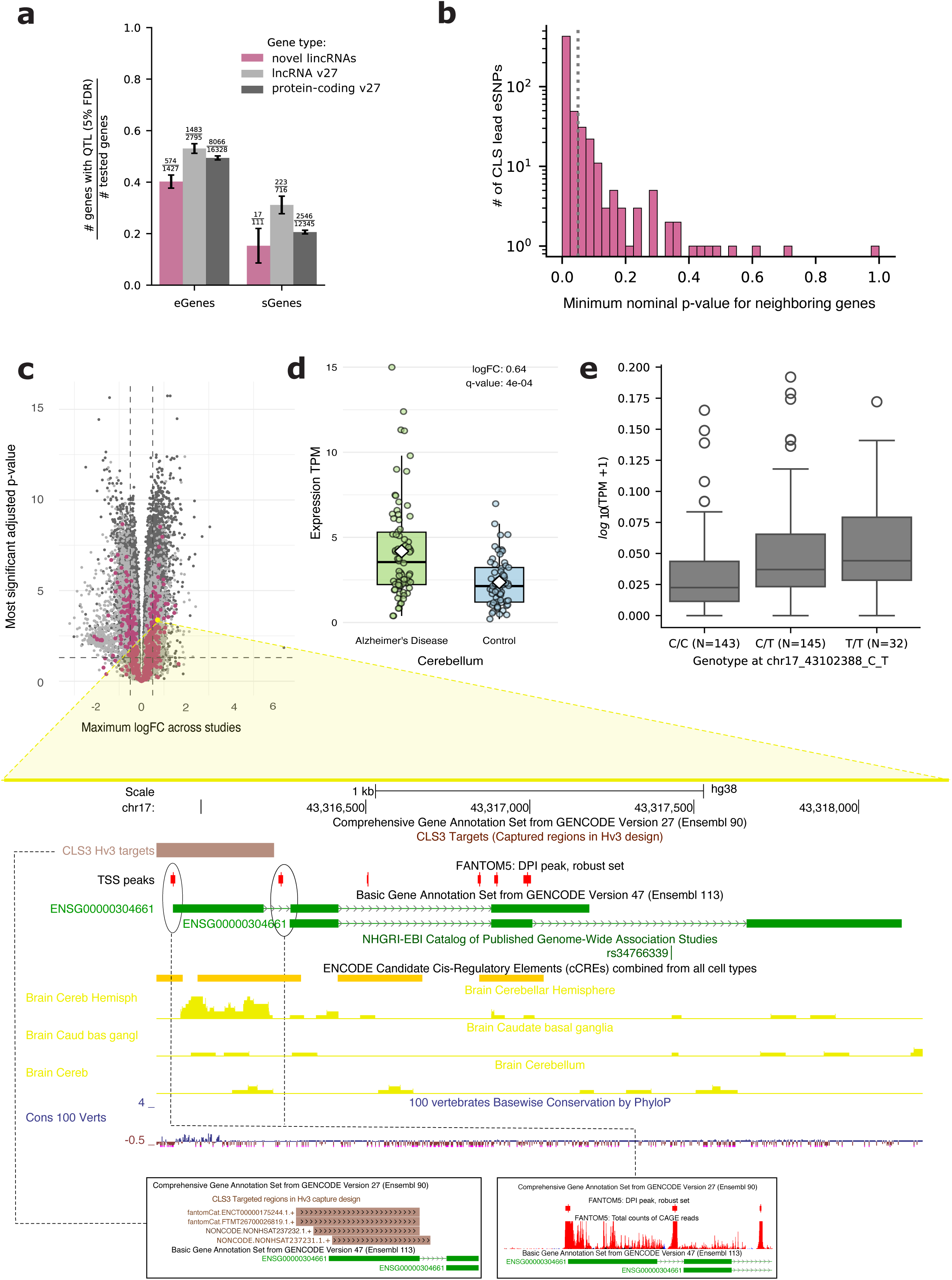
**A)** Proportion of novel lincRNAs, previously annotated lncRNAs (v27), and protein-coding (v27) genes found as eQTLs and sQTLs at 5% FDR. Tested genes for eQTLs are those considered sufficiently expressed, while tested genes for sQTLs are those with > 1 splice isoform sufficiently expressed (see Methods). Error bars represent the 95% confidence interval. **B)** For CLS lead eSNPs, the minimum p-value for association between that eSNP and any other previously annotated gene in a 1MB region. The dotted line is p-value = 0.05. **C)** On the top, volcano plot displaying, for each gene, the largest absolute value of logFC and the most significant q-value, across brain areas and diseases. On the bottom, the locus ENSG00000304661 (green transcripts in the track GENCODE v47 basic). Located on the forward strand, it was captured by targeting NONCODEv5 and fantomCat (track CLS long-read RNAs, subtrack CLS targets). This lncRNA improves the annotation of the originally targeted elements (not shown), and presents strong CAGE peaks at both TSSs. The longest isoform hosts the variant rs34766339, associated with Alzheimer Disease (AD), and annotated as intergenic in the GWAS catalog. **D)** Expression of ENSG00000304661 in cerebellum of patients and controls in syn5550404. The gene is upregulated in AD with a logFC of 0.64 (q-value 0.0004). **E)** Expression of ENSG00000304661 in brain caudate nucleus across the three genotypes at SNP rs10445303 (eQTL for this gene).

### Novel lincRNAs expression across neurological disorders

To investigate the potential pathophysiological role of novel lincRNAs in diseases, we focused on neurological disorders. Indeed, lncRNAs have rapidly emerged as critical mediators in neurological health and dysfunction, with a recent surge in the number of such elements associated with disease development and progression^56^. We used the extended GENCODE v47 annotation to map 1,652 short-read RNA-Seq samples^57–62^, spanning different neurodevelopmental and neurodegenerative conditions, and 1,052 non-overlapping controls. Overall, we found 322 novel lincRNAs differentially expressed between cases and controls in at least one study (**Table S16**). For instance, ENSG00000304661 is upregulated in the cerebellum of Alzheimer’s Disease (AD) patients (**Figure 5C,D**), and we have associated its expression with the upstream SNP rs10445303 in caudate nucleus (q-value 7.9e-07, **Figure 5E**). This lincRNA also hosts in its longest transcript the variant rs34766339, annotated as intergenic, and significantly associated with levels of ADP-ribosylation factor-like protein (ARL4D) in AD^63^.

### Novel lincRNAs expression at the single-cell level

As novel lincRNAs are identified mostly post-capture, they are poorly expressed in bulk RNA samples: for example, they are, on average, 2.3-fold less expressed than annotated lncRNAs in the caudate nucleus samples from GTEx (average TPM 0.33 vs 0.76). However, this is more the result of restricted expression rather than inherently low expression levels.

Indeed, we reprocessed 656 single-cell experiments corresponding to 29 different tissues from the Human Common Cell Atlas^64,65^, leveraging a total of 2,489,458 cells upon stringent filtering (**Figure S24A**). We used v47 as the reference annotation. We found novel lincRNAs to be expressed in a lower proportion of cells than annotated lncRNAs (about 2.4-fold, 2,79% vs 6.75% on average across experiments). However, their cellular expression is comparable (0.56 vs 0.55 scaled counts, on average across experiments and cells, **Figure S24B**). More generally, we found that limited breadth of expression is a more important contribution than intrinsic low expression levels to the relatively low expression of lncRNAs reported in bulk RNA-Seq studies. Indeed, protein-coding genes are expressed on average in 18,4% of the cells, a 6.6-fold more than novel lincRNAs, while their expression levels are only about 1.45-fold more that of intergenic transcripts (0.8 scaled counts on average across experiments and cells, **Figure S24B**). This estimate, however, could be partially biased, as the Human Common Cell Atlas includes single-nuclei experiments.

When clustering cells within each experiment, we identified 8,229 clusters, 2,213 of which featured at least one novel intergenic lncRNAs marker. In total, 1,913 novel intergenic lncRNAs were classified as markers (**Table S17**) and could potentially participate in cell-type specification. For example, ENSG00000307510 (**Figure 6A**) is consistently a marker across a set of single-cell experiments from embryonic hippocampus. This gene is mainly detected in inhibitory neurons, mostly restricted to a subset of somatostatin (*SST*+) GABAergic interneurons. Its expression seems to peak around gestational week 20 (**Figure 6B**), a period marked by high neurogenesis^66^. This novel lincRNA is expressed in adult hippocampus as well as across other brain regions in GTEx (v8), where it is an eGene for SNP rs2613060 in caudate nucleus. The splice-junctions display strong signals of conservation across vertebrates (**Figure 6A**), and we also observe consistent expression in hippocampus as well as across other brain areas across primates (**Figure 6C, Figure S25**). This novel lincRNA hosts in its first intron the variant rs67114353, significantly associated with dementia^67^ and annotated as intergenic in the GWAS catalog (**Figure 6A**).

**Figure 6.**
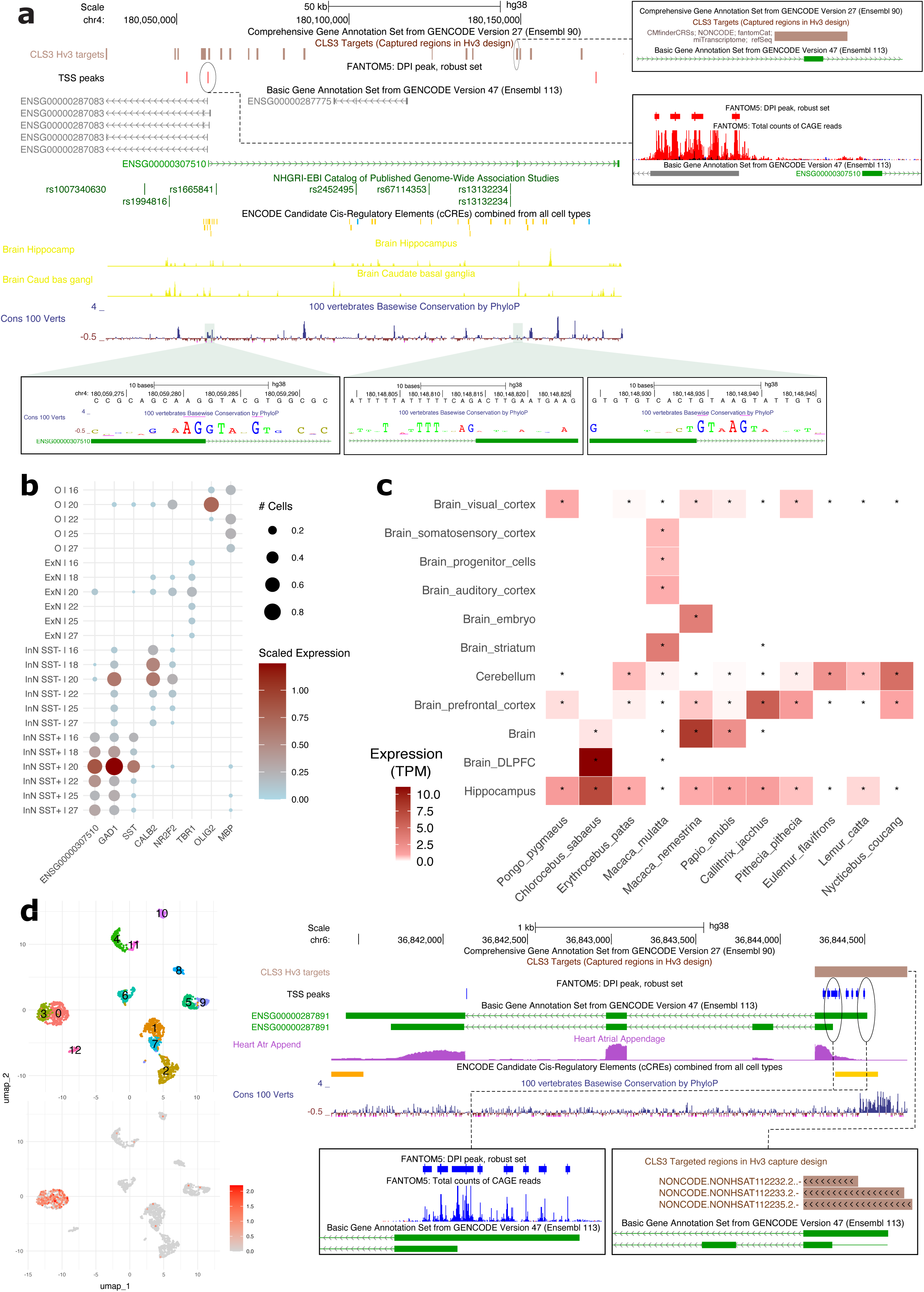
**A)** The novel lincRNA ENSG00000307510 (green transcript in the track GENCODE v47 basic), located on the forward strand, has been captured targeting several catalogs (NONCODEv5, refSeq, miTranscriptome, fantomCat and CMfinderCRSs, for further details see track CLS long-read RNAs, subtrack CLS targets). It presents some CAGE support at the TSS. Despite being mostly detected as a pre-mRNA in the single-cell experiments requantified in this study, exonic read coverage has been detected in GTEx (track GTEx V8 RNA-Seq Read Coverage by Tissue), among the others in brain hippocampus and caudate nucleus, this latter tissue in which it is an eGene. The locus hosts in its first intron the variant rs67114353, significantly associated with Dementia, and annotated as intergenic in the GWAS catalog. This novel lincRNA also shows strong conservation across primates and vertebrates (track UCSC 100 Vertebrates), especially in the first donor and acceptor sites. **B)** Average scaled expression and proportion of cells expressing ENSG00000307510 and other relevant marker genes across the different human embryonic hippocampus experiments. Expressions collected at the same gestational weeks (GW) were averaged. On the y axis, the following cell-types are displayed; Somatostatin-positive inhibitory neurons (InN SST+), somatostatin-negative inhibitory neurons (InN SST-), Excitatory Neurons (ExN), and Oligodendrocytes. Cell-type is assigned to Seurat clusters according to expression of GAD1 (InN), SST (InN SST+), CALB2 and NR2F2 (InN SST-), TBR1 (ExN), and OLIG2 and MBP (Oligodendrocytes); clusters with mixed identities were excluded. Data is grouped according to the cell-type and ordered by gestational week of sampling. For visualisation purposes, dots are shown only when ≥ 5% of the cells in the cluster express the gene. **C)** Expression (in TPM) across different primates species (phylogenetically ordered), for the tissues requantified in this study. An asterisk marks the cases where the data was collected. For visualisation purposes, only species with TPM ≥ 0.5 in at least one tissue are reported, as well as only brain tissues where TPM ≥ 0.5 in > 1 species are displayed. **D)** On top left, UMAP projections of the clustering obtained for heart single-nuclei experiment ERX4319182. Clusters 0, and 3 are novel compared to the quantification with GENCODE v47 upon exclusion of novel lincRNAs, as they were previously a joint group of cells. The scaled expression of ENSG00000287891 (green transcript in the track GENCODE v47 basic), is displayed in the bottom left panel. The locus, located on the reverse strand, presents strong CAGE peaks at the TSSs and has been captured targeting transcripts from the NONCODEv5 catalog (track CLS long-read RNAs, subtrack CLS targets), for some of which the splice-junctions are perfectly recapitulated (not shown). Short-read coverage in Heart Atrial Appendage is displayed (track GTEx V8 RNA-Seq Read Coverage by Tissue).

Similarly, for the experiment reported in **Figure 6D**, the novel lincRNA ENSG00000287891 shows restricted expression to two clusters mainly composed of cardiomyocytes. This locus, supported by a clear exonic read coverage in Heart Atrial Appendage (GTEx v8, **Figure 6D**), shows tissue-specific expression for heart samples, where it is a marker in 63 (43%) experiments requantified in this study.

Finally, by comparing with the clustering obtained excluding the intergenic lncRNAs from the annotation, we found that 850 clusters emerged upon their inclusion. For 151 of these novel clusters we identified at least one novel intergenic lncRNAs marker. These could potentially play a role in the determination of yet undetected cell subtypes (**Table S17**). An example of such is ENSG00000287838, which, in the same experiment reported in **Figure 6D**, is a significant marker for a restricted group of cells associated with a novel cluster (cluster 0, logFC 3.9, q-value 2.7e-57). All these findings suggest that many novel lincRNAs can be used as markers of cellular functions, and may even participate in cell-type determination.

## Discussion

The generation of a catalog of lncRNAs as complete as that of protein-coding genes is essential to fully mine the biological information encoded in the human and mouse genomes, as growing numbers of carefully documented cases attest to the biological relevance of lncRNAs^68–71^. However, in contrast to protein-coding genes, lncRNAs do not show sequence compositional biases, nor strong phylogenetic conservation. Moreover, they tend to have restricted expression patterns, which makes them difficult to be properly captured in bulk RNA approaches. Thus, existing catalogs^16,18,19,24–27^ contain incomplete and fragmentary models and poorly overlap one another. Here, we have used the models in these and other catalogs to design our targeted sequencing approach, which has led to incorporation in GENCODE of hundreds of thousands of human and mouse full-length lncRNA transcripts, consolidating a unified lncRNA catalog for these species.

That such a large number of human and mouse genes remained unannotated twenty-five years after the publication of the first drafts of the human and mouse genomes illustrates both the complexity of the information encoded in the DNA sequence, and the crucial impact of technological advances to explain it. While many of the transcripts included in GENCODE are partially present in some lncRNA catalogs, and are supported by scattered short-read data, it is only thanks to the technological capacity brought over by targeted long-read sequencing that we have been able to reliably identify their complete sequence.

While our work constitutes a major advance toward completing the catalog of human and mouse genes, many lncRNA genes may still remain to be discovered. In fact, there are still a large number of novel CLS models that have not yet been incorporated into GENCODE, simply because they do not reach the minimum short-read support. Many, if not most, are likely to correspond to real genes. In addition, while our experimental design has identified a large fraction of the transcriptional activity in the human genome (91% of protein-coding genes and 67% of the lncRNAs annotated in GENCODE previously to v47), the survey of additional cell-types, conditions, developmental stages, subcellular compartments, and RNA populations (i.e., poly(A)-RNAs) in samples from individuals of diverse genetic background^72,73^, is likely to uncover additional genes. We are reluctant to provide a precise estimate of the number of genes remaining to be annotated, but based on the genomic space available, the density of GWAS hits and other factors, we expect this number to be lower (probably much lower) than the number of novel genes reported here.

GENCODE’s task is limited to map reliable transcriptional loci into the human and mouse genomes, but not to assign specific biological function to them. We believe, nevertheless, that our analyses provide substantial evidence that the novel loci reported here are clearly distinct from background transcriptional noise and that they are associated with biological function. Moreover, while they appear to be poorly expressed in bulk RNA-Seq samples, we often found them expressed in specialized cell-types. Hundreds act as markers, and may even help to uncover previously undetected cell subtypes that had remained “invisible” because of annotation incompleteness. This, together with the greatly enhanced interpretability of omics measurements and of genetic variants contributed by the novel lncRNAs reported here, underlines the critical relevance of accurate gene annotations.

In this regard, the protocols and the data produced by GENCODE as part of their effort to annotate human and mouse genomes could be of great relevance to annotate all eukaryotic genomes, as illustrated in the case of primates, where we confidently predicted hundreds of thousands of previously unannotated lncRNAs. Furthermore, the vast amount of data that we produced here directly or indirectly constitutes a unique resource to train deep learning and large language models. These seem particularly appropriate to deal with the intrinsic semantic nature of biomolecular sequences, as illustrated by the success in predicting protein structures from amino acid sequences^74^, but their accuracy crucially depends on the size and quality of the training data. The combination of these with more traditional AI rule-based systems, building on our extensive experience in manual curation, could be at the basis of automatic methods producing annotations of high quasi-manual quality across the entire phylogenetic spectrum^75^. These annotations are essential to maximize the benefit of projects underway to sequence the genomes of all eukaryotic species on Earth^76–78^.

## Acknowledgements

Research reported in this publication was supported by the National Human Genome Research Institute of the National Institutes of Health under grant numbers 2 U24 HG007234, U24 HG011451, U24 HG012090 and T32 HG000044; Wellcome Trust [WT222155/Z/20/Z]; European Molecular Biology Laboratory. This publication is part of the grant PRE2022-102742, funded by MCIN/AEI /10.13039/501100011033 and by the FSE+. This work and its publication were also supported by the National Science Center PL grant 2021/42/E/NZ2/00434 to B.U.R. and by FWF Austrian Science Fund V-762 (Elise-Richter Program) to A.T. We acknowledge support of the Spanish Ministry of Science and Innovation through the Centro de Excelencia Severo Ochoa (CEX2020-001049-S, MCIN/AEI /10.13039/501100011033), the Generalitat de Catalunya through the CERCA programme and to the EMBL partnership. R.J. is supported by Science Foundation Ireland through Future Research Leaders award 18/FRL/6194 and by the Irish Research Council through Consolidator Laureate award (IRCLA/2022/2500). S.G-L. is supported by the predoctoral program AGAUR-FI grants Joan Oró (2024 FI-1 00570) from the Generalitat de Catalunya and the European Fund Social Plus. M.A. acknowledge the funding from grant PID2021-122726NB-I00 funded by MCIN/AEI/10.13039/501100011033 and by “ERDF: A way of making Europe”, by the European Union. The work was also funded by the European Union (ERC, NovoGenePop, project number 101052538). We are grateful to the CRG Core Technologies Programme for their support and assistance in this work. We would like to acknowledge the CRG Genomics Unit for assistance with short-read Illumina sequencing and the NGS Sequencing Core facility at CSHL headed by S. Goodwin for their assistance in generating PacBio Sequel II data. We thank the Guigó laboratory for their valuable input and help with sample handling. We acknowledge T. Montserrat-Ayuso and S. Booeshaghi for the discussion on single-cell data processing, and G. Santpere for the inputs on brain cell-type annotation. We thank R. Garrido Enamorado and R. Carbonell Garcia (CRG) for administrative support. Disclaimer: The content is solely the responsibility of the authors and does not necessarily represent the official views of the National Human Genome Research Institute or the National Institutes of Health. Views and opinions expressed are however those of the authors only and do not necessarily reflect those of the European Union or the European Research Council. Neither the European Union nor the granting authority can be held responsible for them.

## Methods

### Samples, library preparation and sequencing

#### Ethical statement

This study involved the use of commercially obtained human total RNA samples and mouse tissues for RNA extraction. Ethical approval from an institutional review board (IRB) was not required for the use of human RNA, as these were pre-isolated, anonymized commercial products purchased from certified vendors. All procedures involving these materials adhered to established ethical standards for research using commercial human biological products. For mouse-derived RNA, tissues were obtained from animals purchased through the PRBB (Parc de Recerca Biomèdica de Barcelona) animal facility. All animal procedures were approved by the PRBB Institutional Animal Care and Use Committee and conducted in accordance with institutional and national guidelines for the care and use of laboratory animals.

#### Samples

Total RNA was obtained from a diverse set of human and mouse tissues and cell lines. No biological or technical replicates were conducted in this study.

Commercial human total RNA samples were purchased, including RNA from adult and embryonic heart, brain, liver, white blood cells, testis, placenta, and induced pluripotent stem cells (iPSCs). One pooled library was generated from four human cell lines (HCT-116, IMR-90, MCF-7, A549). Another pooled library included RNA from 13 tissues: colon, bladder, lung, thyroid, trachea, thymus, esophagus, cervix, adipose tissue, skeletal muscle, spleen, prostate, and small intestine.

Mouse tissues were obtained from adult and embryonic C57BL/6 mice, and total RNA was extracted in-house. These included heart, brain, liver, white blood cells, testis, and embryonic stem cells (ESCs). A pooled library was prepared from 12 mouse tissues: colon, bladder, lung, thymus, esophagus, kidney, ovary, adipose tissue, skeletal muscle, spleen, prostate, and small intestine.

#### RNA extraction

Total RNA extraction from mouse tissues was performed in-house using TRIzol reagent followed by purification with the PureLink RNA Mini Kit (Thermo Fisher Scientific). Human tissue total RNA was obtained from commercial sources (**Table S2**). The purity of RNA samples was evaluated using a NanoDrop One spectrophotometer (Thermo Fisher Scientific), while RNA concentration was measured using the Qubit High Sensitivity RNA Assay Kit (Thermo Fisher Scientific, Cat. No. Q32852). RNA integrity and quality were assessed using RNA Nano chips on an Agilent Bioanalyzer system, using RNA Integrity Number (RIN) to confirm sample quality.

#### Capture design

e target selected regions of the human (hg38) and mouse (mm10) genomes to design a custom capture array. Probe design was performed by NimbleGen, using 120bp oligonucleotides to tile across the selected regions. The number of probes per region was adjusted based on the region’s length to ensure full coverage. The regions targeted for probe design included *i)* non-GENCODE lncRNA annotations^18,19,24–27^, *ii)* small non-coding RNAs, *iii)* enhancers^28^, *iv)* RNAs predicted to contain evolutionary conserved structures^29,30^, *v)* regions either hosting non-coding GWAS hits^31,32^, *vi)* or showing characteristics of protein-coding genes as predicted by PhyloCSF^33^, *vii)* or being evolutionarily conserved^34^. When available, probes were designed against the corresponding datasets for human and mouse (NONCODE, refSeq, phyloCSF, and small RNAs), while eight annotations (i.e., miTranscriptome, fantomCat, bigTranscriptome, CMfinderCRSs, GWAScatalog, UCE, fantomEnhancers, and VISTAenhancers) were lifted-over from the human to the mouse genome. In addition, we designed probes against GENCODE+^19^ catalogs, which is the union of either GENCODE v20 (for human) and GENCODE vM3 (for mouse), with CLS^21^ models from the pilot phase. These probes were then used in the CLS protocol for targeted full-length transcript sequencing^21,22^. In total, we targeted 176,435 regions (84,103,329bp, 2.9% of the genome length) in the human genome, of which 116,383 (66%) were intergenic with respect to GENCODE v27 and therefore targeting 54,545,313bp of the unannotated space. Similarly, we targeted 148,965 regions (66,937,555bp, 2.8% of the genome length) in the mouse genome, of which 114,926 (77%) were intergenic with respect to GENCODE vM16, covering 50,735,573bp of the unannotated space. In total, 107,701 regions were orthologous between human and mouse (**Table S1**).

#### Spike-in controls

A 1:1 mixture of capped ERCC^35^ and Lexogen SIRV spike-in controls was prepared as previously described^23^ and added to all samples prior to cDNA library preparation. This mixture served as an internal standard to trace and monitor the sample preparation process. To measure the efficiency of our protocol, we included in our capture array probes designed to target 42 among the least abundant ERCC spike-ins, excluding the 8 most abundant ones (**Figure S1A**).

#### Library preparation and sequencing

Total RNA from human and mouse samples was used to prepare double-stranded cDNA libraries following the CapTrap-Seq protocol^23^, designated as pre-capture libraries. The pre-capture libraries were divided into two aliquots; the first processed to accommodate the requirements of the Oxford Nanopore (ONT) or PacBio sequencing platforms. The second aliquot was subjected to capture experiment using a capture panel (post-capture libraries), as described in our previously published CLS protocol^21,22^ and sequenced with ONT and PacBio following the same strategy as for pre-capture libraries. The platform-specific sequencing libraries prepared from each aliquot were loaded onto ONT or PacBio long-read sequencing platforms. Sequencing was performed using the respective manufacturing protocols for amplicon sequencing (Amplicon by Ligation SQK-LSK109 for ONT and SMRTbellTM Express Template Prep Kit 2.0 for PacBio). For ONT sequencing, a MinION device with ONT R9.4 flow cells was used, following the standard MinKNOWN protocol script. PacBio data were generated using the Sequel II platform. One flow cell per sample was used on each platform without multiplexing.

#### PacBio data preprocessing

The preprocessing and quality assessment of PacBio sequencing data were conducted externally at Cold Spring Harbor Laboratory (CSHL) in accordance with the manufacturer’s protocols. To make the PacBio data compatible with the downstream LyRic processing, PacBio FASTQ files containing CCS (Circular Consensus Sequencing) reads were generated using the pb_gen workflow (https://github.com/guigolab/pb_gen).

#### ONT data preprocessing

Basecalling for ONT sequencing was performed using Guppy v6 SUP (developed by ONT and only available for ONT community members, https://nanoporetech.com/document/Guppy-protocol). NanoPlot^79^ was utilized to generate metrics, including read length distributions, quality scores (Q-scores), and the total number of reads, providing insights into the overall performance of the sequencing run. Additionally, the *split_on_adapter* utility from the Duplex Tools suite^80^ was employed to evaluate the presence of concatamers in the ONT sequencing data. This utility was adjusted in accordance with the CapTrap-Seq adapter and primer sequences. The utility was run twice to achieve a higher stringency in detection of concatemeric reads. In the first step, we split the reads based on the presence of the ONT adapter linked to the CapTrap-Seq primer. In a second round, read splitting was based solely on the presence of CapTrap-Seq adapters within the reads. As a final step of this quality control, the multi-split reads were discarded.

#### Data produced

In total, we sequenced 88 samples producing 736 million long-reads (631 million reads ONT and 105 million reads PacBio). Detailed statistics can be found at (https://guigolab.github.io/CLS3_GENCODE/summary_GENCODE.html).

### Generation of CLS transcripts models and inclusion in GENCODE

#### Long-read data processing and generation of per-sample CLS models

The CLS long-read data were initially processed with LyRic^36^ for each individual sample separately (**Supplementary Materials**) to obtain high-confidence transcript models. Long reads obtained mapped to hg38 and mm10 genome assemblies for human and mouse, respectively, using minimap2^81^ (default settings, in addition to `-x` preset as “`splice:hq`” for PacBio and “`splice`” for ONT, `--MD --secondary=no -L`).

The following steps were performed as part of the LyRic pipeline. Read orientation was inferred through detection of poly(A) site and splice-junctions (SJ) orientation. The *polyAmapping.pl* LyRic module infers the reads strand based on reads-to-genome alignment (arguments used: `--minClipped=10`; sets the minimum length of A or T stretch required to call a poly(A) site, `--minAcontent=0.8`; decides the required fraction of nucleotides A (or T, if minus strand), `--discardInternallyPrimed`; to avoid false poly(A) sites as a result of internal mis-priming during the cDNA library construction, `--minUpMisPrimeAlength=10`; minimum length of genomic A stretch immediately upstream a putative site required to call a false positive). The read strand was also inferred from SJ orientation through majority vote. The final read strand is decided based on the poly(A)-SJ agreement. In cases where a conflict between poly(A) tail and canonical SJs strand assignment was detected, this latter was prioritised. Such cases were later tagged as possible artifacts.

To obtain high confidence reads, we filter to retain, for spliced reads, only those with *i)* canonical SJ (GT|GC/AG), *ii)* SJ not surrounded by direct repeats and *iii)* high Phred score (PacBio, 30; ONT; 10) around the SJs (+/- 3bp window). Reads that have a poly(A) tail aligned to stretches of A nucleotides in the genome were also filtered out, since they may represent chimeric transcripts originating from internal priming.

As a final step of the LyRic pipeline, reads were merged using tmerge^82^ to get non-redundant per-sample models, with the following arguments: `--exonOverhangTolerance 8 --minReadSupport 2 --endFuzz 8`. The models were tagged as 5’-supported if they overlap CAGE tags^41,42^ while the aforementioned poly(A) sites were used to tag support at the 3’ ends.

The sort read data produced for the pre-capture samples was not used to build the CLS models.

#### Merging of CLS models across samples

In a second step, LyRic merged the CLS transcript models across all samples to produce a unique set of models for human and mouse. The per-sample CLS models are pooled together and merged using tmerge^82^ (with default parameters). Since the models produced by LyRic are tagged with 5’ and 3’ support, we produced an “anchored” set, where CLS models with tagged termini are not extended beyond by merge with other models. In this way, we protect the supported termini to preserve potential alternative, tissue-specific, transcriptional start and end sites, while reducing the redundancy present across samples (**Figure S2**).

During the merging process, meta-data associated with each sample (i.e., sequencing platform, capture status, developmental stage, tissue of origin, etc.) is propagated to the anchor-merged CLS models. It is therefore possible to recover the original models from the anchor-merged ones. The final set of (anchor-merged) CLS models (and the associated meta-data) produced by LyRic can be found at: https://github.com/guigolab/CLS3_GENCODE/tree/main/data_release#cls-anchored-models

To reduce redundancy, LyRic collapses the (anchor-merged) CLS models into intron-chains, merging transcripts sharing the same internal exon-intron structure while ignoring the termini variation, using gffcompare^83^. Monoexonic models are also merged into a single transcript when overlapping more than 50% of their length, irrespective of the support at the termini. The joint set of intron chains and extended monoexonic transcripts will be referred to as CLS transcripts and can be found at: https://github.com/guigolab/CLS3_GENCODE/tree/main/data_release#cls-transcripts. LyRic can also cluster the merged CLS models in uniquely identifiable loci using bedtools^84,85^.

The set of CLS transcripts, together with the raw long-read data, was used by TAGENE to produce the final set of lncRNA transcripts to be included in GENCODE. As part of the processing by LyRic, transcripts are searched for potential artifacts according to the TAGENE specifications (**Supplementary Materials**), and tagged. CLS transcripts overlapping annotated pseudogenes or located antisense to annotated genes were filtered out. All tags were propagated back to the original set of CLS anchored models, where, in addition, transcripts in which the poly(A) tail and splice-junctions orientation provided conflicting strand information are also tagged as potential artifacts.

LyRic produces comprehensive summary statistics per sample (statistics for this project can be found here (https://guigolab.github.io/CLS3_GENCODE/summary_GENCODE.html), as well as a track hub of the final set of CLS models for visualization in the UCSC Genome Browser.

#### TAGENE pipeline

We developed the TAGENE workflow to manage the conversion of CLS models to GENCODE gene annotation. Firstly, TAGENE removes all transcripts potentially considered artefacts. These include models that are antisense to protein-coding genes based on any overlap between the regions defined by the first and final gene or model coordinate. This was done primarily to remove models that were mis-stranded by the alignment process. We also removed all models located entirely within the genomic bounds of GENCODE processed pseudogenes on either strand, primarily to filter out reads that should in reality have been mapped to their parental locus. In particular, annotators noted the tendency of parental reads to be incorrectly aligned at processed pseudogenes due to the preferential alignment of the poly(A) tail in the read against the poly(A) tail retro-insertion sequence in the genome. Next, in order to devise a filter based on supporting evidence, we examined how annotator confidence in a CLS model correlated with the support for its splice-junctions as of recount3 short-read RNA-Seq data. Conservatively, we opted to filter out all models that contained any intron supported by < 50 reads, although we estimate that around half of the models rejected solely on this basis are likely to be correct. In addition, to minimise the introduction of false splice-junctions, we removed the transcripts that contained introns mapping entirely within the boundaries of a single tandem repeat. We also checked that all novel splice-junctions were well supported by read alignments, rejecting models built from alignments with mismatches or indels close to the splice sites. For further details refer to the **Supplementary Materials.** Finally, a substantial complication of this work from an annotation point of view is the potential for CLS models to join together previously distinct lncRNA loci in GENCODE. We complemented the TAGENE predictions with manually curated transcripts to help resolve the cases where new models had the potential to merge existing distinct lncRNA genes into single loci.

#### Extending GENCODE annotation

For some downstream expression re-quantifications, the human GENCODE v47 was extended with the most reliable intergenic CLS loci. To do so, we selected the intergenic spliced CLS transcripts, discarding any artifacts except the transcripts supported by < 50 recount3 reads (88% of the human spliced CLS transcripts not included in the final annotation). To reduce redundancy, we merged these transcripts using tmerge with `--exonOverhangTolerance 8`. Further, they were grouped into loci and added to v47, to get an “extended” annotation (1,066 genes and 1,309 transcripts).

#### Decoy models

We generated decoy models by randomly relocating the spliced structures of the CLS loci (n = 45,819) in the intergenic space. We first used bedtools^84,85^ to merge the CLS loci to GENCODE v27, extend the boundaries of each entry by 10kb in both directions, and subtract those coordinates from the reference chromosomes (GRCh38/hg38, excluding the mitochondrial genome). From this newly defined intergenic space were removed the ENCODE blacklist regions (ENCFF356LFX) and the centromeres^86^, as well as other genomic gaps (UCSC table browser, group ‘Mapping and Sequencing’, track ‘Gap’, table ‘gap’, format ‘bed’). For each CLS locus, drawn at random, the set of associated spliced transcripts was relocated in one of the aforementioned intergenic chunks chosen at random, ensuring no overlap among them. Loci for which those requirements could not be fulfilled were excluded. Strand information was not propagated. Eventually, we obtained 17,223 random loci (85,283 spliced transcripts), covering around 250 million bases (70.2% of the viable unannotated space).

### The unified lncRNA catalog for human and mouse genomes

To account for variability across annotations, gene loci were built using *buildLoci*^87^ for each lncRNA targeted annotation (except for GENCODE v27 and v47). A non-redundant lncRNA catalog (lncRNA-merge) was created by merging transcripts from all targeted annotations using tmerge (`--exonOverhangTolerance = 8`). Monoexonic transcripts were excluded from merging. Gene loci boundaries for lncRNA-merge were generated using *buildLoci*. Gene-level overlaps were computed with bedtools intersect (default settings plus `-f 1 -F 1 -e -s`). Quality comparisons between GENCODE v47 and other annotations followed the approach detailed in previous work^19^, with two key modifications. First, the “completeness” metric was replaced with “support” defined as 5’ ends overlap with robust phase 1/2 FANTOM CAGE clusters^41,42^ (n = 201,802) within ±50bp, and presence of a canonical polyadenylation motif^88^ within 10–50bp upstream the 3’ ends. Second, transcript structure accuracy was evaluated based on recount3^89^ support of splice-junctions (≥ 50 reads per junction). The resulting proportions are shown as pie charts.

#### LncRNA orthology

LncRNA orthology was predicted using an updated version^40^ of the ConnectOR pipeline^90^, which performs reciprocal LiftOver of syntenic regions between human (hg38) and mouse (mm10/mm39) genomes. ConnectOR was run in exon mode with default parameters to generate strand-specific orthology predictions. Instances labeled as “not lifted” or “one-to-none” were considered non-orthologous. A negative control set was generated using shuffleGTF (https://github.com/cobRNA/utilities), which randomizes gene positions while preserving exon structure. We ran it 100 times per comparison and used median value to estimate false discovery rate (FDR). The FDR was ∼3% for GENCODE v27/vM16 and ∼6% for v47/vM36.

### Protein-coding genes and pseudogenes

#### PhyloCSF search

To find CLS transcripts likely to contain additional, not previously annotated, conserved novel protein-coding regions, we searched for complete open reading frames (ORFs) *i)* at least 25 codons long, *ii)* some portion of which not annotated as protein-coding or pseudogene or antisense to a either two in GENCODE v43 or vM32, *iii)* at least 15 nucleotides of which were not annotated as transcribed in v27 or vM16, *iv)* having a positive PhyloCSF score and positive PhyloCSF-Psi score, *v)* whose alignment had relative branch length at least 0.4, and *vi)* for which there was no species in which some portion of the alignment overlapped the alignment of an annotated coding region (which usually indicates a pseudogene). PhyloCSF^33^ was run using the `58mammals` parameters on alignments extracted from the 58 placental mammal subset of the hg38 100 vertebrates whole genome alignment (for human) or the 40 placental mammal subset of the mm10 60 vertebrates whole genome alignment (for mouse), downloaded from the UCSC Genome Browser. The resulting candidates were then manually examined using CodAlignView (https://data.broadinstitute.org/compbio1/cav.php) to find novel, likely coding, regions.

#### Proteomic search

Proteomic evidence for the translation of the CLS transcripts came from large-scale mass spectrometry analyses. In the case of the human CLS data, the translated CLS regions and the human reference protein sequences annotated in GENCODE v43 were mapped to spectra from four large-scale, tissue-based, proteomics experiments^91–94^, while for mouse, the CLS translations and the GENCODE vM32 reference protein sequences were mapped to spectra from a single compendium of normal tissue experiments^95^. Spectra from the five experiments were downloaded from ProteomeXchange^96^.

Peptide-spectrum matches (PSMs) were generated with COMET^97^ using default parameters, including a maximum precursor charge of 4, a maximum fragment charge of 3 and a mass tolerance of 10. Peptides were limited to a minimum length of 7 and a maximum length of 40 amino acids. Oxidation of methionines was allowed. The PSMs detected by COMET were post-processed with Percolator^98^ using default parameters, including setting the test and training FDR to 0.01. Only fully tryptic peptides with up to two missed cleavages and PSM that had Percolator posterior error probabilities (PEP) values below 0.0002 were considered. These filters ensure a FDR of the novel peptides of 1.41% for human (0.25% at the PSM level) and 2.04% for mouse (0.28% for PSMs).

Peptides just a single amino acid different from annotated protein sequences were further disregarded, as explainable by single amino acid variants or post-translational modifications of the annotated protein sequences. To validate the translation of a CLS ORF, at least two non-overlapping peptides were required.

#### Pseudogenes

We used featureCount^99^ to quantify expression at the CLS loci level, leveraging the raw long-read data generated by this study. The loci were tagged with same-strand annotation overlaps to GENCODE v43 for the human and GENCODE vM16 for the mouse. The following parameters were applied: `-L -s 0 -T 8 -t exon -g gene_id`. For downstream analysis, gene IDs from the CLS transcripts were mapped to the gene IDs in the GENCODE annotation, using bedtools intersect with a requirement of at least a 1bp overlap. Next, protein-coding genes were categorized into two groups: parent and non-parent, based on the parent protein-coding gene identified by PseudoPipe^100^ (**Table S7.4** for human and **Table S7.5** for mouse). Using the mapping to GENCODE, pseudogenes and parent genes relationships in the CLS transcripts were established.

We then analyzed gene expression patterns across various experimental conditions using DESeq2^101^. The experiments involved three variables: *i)* whether the sample was captured, *ii)* the sequencing technology used, and *iii)* the tissue origin of the sample. To assess the individual effects of capturing and sequencing technology, we controlled for the remaining two variables as covariates. We defined significantly differentially expressed genes as those with an absolute log_2_FC > 1 and FDR-adjusted p-values < 0.001.

### Functional interpretability of the human genome

#### TSS support

Sets of non-redundant representative TSSs were built for *i)* lncRNAs (v27), *ii)* novel CLS models, *iii)* protein-coding genes (v27) and *iv)* decoy models. Herein, consecutive TSSs within 100bp from each other were merged and associated to a single representative, chosen to be the median TSS among the collapsed ones. Specifically to generate the set for novel CLS, their TSSs (extended 100bp) were previously intersected with GENCODE v27 (184,093 TSSs) to define the set of novel TSSs that do not overlap already annotated ones. Upon merging, we obtain a non-redundant set of 80,284 novel TSSs.

#### Support by histone modifications

New classification of ENCODE4 cCREs, specifically PLSs, pELSs, dELSs and CA-H3K4me3, was obtained based on their distance between their midpoint and the closest novel TSS; this distance was computed using bedtools intersect, run with default parameters along with the additional options `-d -t first`. By design, all PLSs and pELSs’ classification were kept as they were; CA-H3K4me3 cCREs that were within 201bp from the closest TSS were re-classified as PLSs, and dELSs within 2,001bp from the closest TSS were re-classified as pELSs. This same analysis was performed using decoy models TSSs.

#### Transcription factors binding

The ChIP-Atlas 3.0 database^47^ integrates 428,000 ChIP-Seq, ATAC-Seq and Bisulfite-Seq experiments from six species. We downloaded the summary (**fileList.tab**) file from the ChIP-Atlas portal. We filtered this file for the experiments that contain “hg38” in the genome assembly (second) column and “TFs and others” in the class (third) column. We next filter the type (fourth) column by the valid name of human transcription factors and the cell-type (sixth column) for “All cell-types”. Thus, we obtained a list of 1,806 human transcription factors (TF) for which there is at least one ChIP-Seq experiment in ChIP-Atlas. For each TF we downloaded the full collection of peaks across all tissues at FDR < 1e-5 threshold. We merged the peaks for the same TF (1bp overlap) and obtained 65,155,727 peaks in total (36,077 on average per TF). Peaks for different TFs may overlap each other.

#### GWAS analysis

We pruned the NHGRI-EBI GWAS catalog^32^ (downloaded on 24 October 2023) down to 134,059 unique variants, using a greedy approach similar to what previously described^102^. We ranked SNPs by significance and iteratively collapsed them within a 5kb window, following a top-down approach, assigning each entry to its representative signal to ensure an unbiased calculation of GWAS density. Additionally, all associations falling within the HLA locus (for hg38, chr6: 29,723,340 - 33,087,199) were discarded. We used bedtools to intersect the whole catalog against *i)* novel lincRNAs, complemented with intergenic spliced CLS transcripts not included solely because of a low recount3 score (n = 9,772), *ii)* the set of randomly generated decoy models. The GENCODE v27 geneset, further split into *iii)* protein-coding, and *iv)* lncRNAs which overlap with protein-coding genes was no longer than 10% of their length (n = 8,922). Finally, the intergenic space with respect to either vi) GENCODE v27 or vii) GENCODE v47. For the purpose of the current investigation, the genomic space was restricted to reference chromosomes only (mitochondrial excluded), refined as previously described for decoy models. Proximal regulatory regions have been considered the 2kb extensions at both ends.

We used EPIraction^48^ to predict distal enhancers. Briefly, we added the TSS of decoy models to the list of EPIraction genes and re-quantify expression in all EPIraction tissues. We considered the decoy genes enough expressed if they attracted at least 0.05 TPMs or 20 reads. We predicted the enhancers for expressed decoy genes similarly to other genes in EPIraction. Depending on the threshold one enhancer can be predicted to regulate several genes, moreover the particular enhancer may regulate one gene in one tissue and different gene in another. We chose a threshold of 0.02 and considered the particular enhancer regulating the protein-coding gene if there is at least one tissue where they were predicted interacting with score above threshold and no other CLS or annotated lncRNA or decoy genes regulated by this enhancer at this threshold in any tissue. Similarly we developed a list of candidate regulatory regions for other biotypes.

The GWAS density was computed dividing the total number of unique hits by the total genomic area spanned [1]:

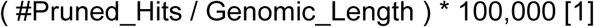

The significance of our results was assessed by means of pairwise Wilcoxon tests on the exonic GWAS density per transcript computed across *i)* the set of novel lincRNAs, *ii)* the set of decoy models, *iii)* the protein-coding annotated genes, and *iv)* the annotated lncRNAs refined as detailed above. Finally, GWAS density in the ±15kb region flanking gene bodies was computed with deeptools v3.5.2^103^ using a bin size of 200bp, and setting the TSS to TTS window size to 5kb.

#### Long transcripts act as hosts of orphan small RNAs

The small RNA set comprises GENCODE genes of gene types miRNA, snRNA, snoRNA, scaRNA, sRNA and Mt_tRNA, as well as tRNAs from a complementary GENCODE annotation file. The long RNA set contains the remainder, with protein-coding genes, lncRNAs and pseudogenes being the most prominent biotypes. To assess the position of small RNAs and their putative host genes in the genome, we partitioned the genome into genic and intergenic regions, and the genic partition further into exonic and intronic. Partitions were created both for stranded annotations, as well as independent of their strand. The former were used to identify host transcript regions to exclude any influence on genome composition and conservation by genes on the opposite strand. When studying the small RNA-host gene relationships, intergenic is defined as the space between long genes on the same strand, because miRNAs and snoRNA are co-transcribed with their hosts and post-transcriptionally processed mostly from introns. Studying conservation of splice sites in lncRNAs we excluded protein-coding genes, to avoid any sequence conservation signal interfering. We therefore defined intergenic/non-protein-coding space as the intervals between protein-coding genes (definition see above), not considering their strand. By comparing the positions of small RNAs (see above), long RNAs (see above), intergenic space and CLS loci, we created new subsets, for which we propose biological relations and functions. MicroRNAs and snoRNAs, for instance, reside in introns of host genes, i.e., lncRNAs or protein-coding genes.

#### ncORF

We analyzed three Ribo-Seq datasets of testis, three of liver and three of brain from both human and mouse species^54^ (ArrayExpress accession number E-MTAB-7247). Translated sequences were identified with RibORF v2.0^104^ with default threshold, more than 10 reads per sequence detected, and found in at least one of the three samples per tissue. For the subsampling analysis, we selected the sample with the minimum number of reads among all samples (28,202,132 reads), and then randomly took reads from the other samples so that they all had this same number of reads.

#### Sequence conservation across mammals

We used mammalian conservation of human lncRNAs as a metric to compare the GENCODE novel lncRNA annotations with previously existing lncRNAs and protein-coding genes. We obtained PhyloP scores for the Zoonomia 241-way mammalian alignment from the UCSC Genome Browser^105^. We evaluate conservation per-transcript and per-unique intron. Since we aimed to compare the nature of GENCODE annotations rather than examine precise conservation levels, we chose these straightforward methods, focusing primarily on the per-transcript analysis, for which we computed mean exon and splice-junction PhyloP scores.

Transcripts with more than 10% of the exon bases not having PhyloP scores or having any of its splice-junction bases lacking scores were excluded. Similarly, all unique intron splice-junction bases must have scores to be included. We set the neutrally evolving range based on the PhyloP score distribution from -1.0 to 1.0. To avoid confounding conservation signals from protein-coding genes, we separately analyzed lncRNAs overlapping protein-coding loci. The breakdown of the transcript categories examined and the frequencies of accelerated, neutral, and conserved PhyloP scores are given in **Table S9**.

### Using the CLS transcripts in the lncRNA annotation of primate genomes

The extended GENCODE annotation and decoy models were lifted to 32 primate genome assemblies (7 clades - **Table S13**) using the LiftOff^106^ pipeline, which utilizes the minimap2 mapper with the following parameters: `-a --end-bonus 5 --eqx -N 50 -p 0.5`. A database was initially constructed using the `-g` option and subsequently applied with the `-db` option. The `-infer_genes` and `-infer_transcripts` options were used to infer absent gene or transcript models. The `-polish` option was employed to realign exons and restore proper coding sequences in cases where lift-over resulted in start/stop codon loss or introduced in-frame stop codons. In addition to the lifted coordinates, two measures were computed: coverage (the proportion of the reference gene length matching the target genome) and sequence identity (the percentage of aligned nucleotide positions remaining identical between the reference and lifted annotations). Gene models with coverage ≥ 0.5 were considered valid. The conservation of lncRNAs projected across primates was carried out at both species and clade level. A lncRNA was considered well-projected if the coverage was ≥ 0.5 for at least 50% of the species included in the clade. To build the augmented annotation including RefSeq reference and GENCODE v47 lifted models two functions from the AGAT^107^ tool were used: `agat_sp_merge_annotations.pl` to combine the annotations, and `agat_convert_sp_gff2gtf.pl` to convert the resulting GFF file to GTF format. The resulting number of genes identified in each annotation is available in **Table S14**.

To quantify the expression of the novel genes in non-human primate species we searched European Nucleotide Archive^108^ for the RNA-Seq data for all primate (tax_id=9443) and not human (tax_id=9606) species. We filtered out long-read RNA-Seq experiments (by read length and corresponding annotations) and single-cell experiments that encode the information about the barcodes in the first read. Having multiple samples for one particular tissue we preferentially selected the ones with pair-end reads. If we had more than one study or more than one sample we first took the study with the highest amount of reads from the tissue of interest and then continued adding the samples until we reached 250 million of reads. We did not mix together single-end and pair-end reads. We used STAR^109^ (ENCODE standard options for RNA-Seq) to map these reads to the corresponding primate genomes and transcriptomes augmented from GENCODE v47. To quantify gene expression, we run RSEM^110^ with default parameters, without the information about RNA strandness.

### Novel lncRNAs show distinct patterns of expression across cell-types and diseases

#### Genetic regulation of novel lncRNAs

RNA-Seq from the GTEx caudate nucleus brain samples were quantified with GENCODE v47 following the pipeline described in GTEx Consortium^55^ (https://github.com/broadinstitute/gtex-pipeline), using all software versions for v10. The eQTLs and sQTLs were called using the v10 GTEx pipeline (q-value < 0.05). Expressed genes for eQTL calling were those that passed GTEx expression thresholds: ≥ 0.1 TPM in ≥ 20% samples and ≥ 6 reads (unnormalized) in ≥ 20% samples. Expressed genes for sQTL calling had at least two splice isoforms that fulfilled the aforementioned requirements.

To determine whether eSNPs for novel lncRNA eGenes associated with other nearby genes, we calculated the minimum nominal p-value for association between the lead eSNP and any other gene in a 1MB window from the eSNP.

#### Neurodisorder meta-analysis

The extended GENCODE annotation was used to map 1,652 RNA-Seq samples^57–62^ from patients with Alzheimer’s Disease (AD, n = 739 samples) [syn5550404, syn3159438], Progressive Supranuclear Palsy (PSP, n = 158 samples) [syn5550404], Pathological Aging (PA, n = 89 samples) [syn5550404], Schizophrenia (Scz, n = 255) [syn12299750, syn18097439, syn4590909], Bipolar Disorder (BP, n = 73) [syn18097439, syn4590909], Autism Spectrum Disorder (ASD, n = 338 samples) [syn4587609], and overall 1.052 non-overlapping controls, from different areas across the major brain lobes and cerebellum. The data obtained from individual datasets were processed separately, in a similar way as previously described^111^. Starting from the raw FASTQs, the data were processed using grape-nf v1.3.0^112^, using the human genome build GRCh38/hg38. Statistics about the mappings were obtained using MultiQC v1.14^113^, Picard v2.6.0^114^, and QualiMap v.2.3^115^, to check for quality and sample outliers for each study independently. Principal components were calculated from sequencing statistics and used as covariates in downstream analyses, when deemed necessary. Within each study, the analyses were carried for each brain area, for each disease, separately. To filter out lowly expressed genes while preserving the signal gathered from novel lincRNAs, only those genes with both TPM ≥ 0.5 and count ≥ 10 in at least 50% of the samples were kept for further analysis. We identified 27,022 unique genes passing the expression threshold in at least one brain area, in at least one study, of which 822 are novel lncRNAs. Outlier samples were detected using standardized network connectivity Z-scores and removed when < -2^116^. Differential expression analysis, using limma v3.57.7^117^, was conducted after adjusting for known batch effects with ComBat-Seq from sva v3.49.0^118^. The model below was tweaked within each analysis to account for study-specific relevant metadata. DEGs were considered significant when FDR-corrected p-value < 0.05.

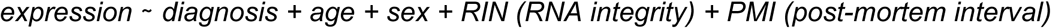

Finally, p-values obtained for each independent analysis, within the same disease, were integrated using the inverse normal function from the metaRNASeq v1.0.7^119^, providing the population sizes as weights. To ensure stringency, coherence of fold-change direction across analyses was required.

#### Single-cell Human Commons Cell Atlas reprocessing

The extended GENCODE v47 annotation was used to reprocess the single-cell Human Commons Cell Atlas^64^, comprising 656 individual datasets encompassing 29 different tissues. We requantified the data leveraging the infrastructure of algorithms for single-cell analysis (https://github.com/cellatlas/human)^120^. We adopted the `nac` workflow^121^, including for quantification the reads that pseudoalign to multiple genes (`--mm`) and producing the total summed count matrix (`--sum total`).

We used the bustools-filtered count matrix to assess the overall effect of novel lincRNAs expression on cell-clustering. To do so, we employed Seurat (v5.1.0)^122^, filtering cells with median number of expressed features ≥ 3 times the median absolute deviation (MAD)^123^. Cells exhibiting extremely low MALAT1 expression were detected enforcing a lower threshold of 5 MAD, set to zero only in case of blood samples. We also employed an upper threshold of 3 MAD for the fraction of mitochondrial reads, setting a maximum ceiling of 80% and restoring it to 5% when lower^124^. Doublets were removed using DoubletFinder (v2.0.4)^125^. We discarded 34 experiments counting less than 50 cells.

After all filtering and QC measurements we formed two read-count matrices; one including *i)* the whole GENCODE v47 geneset and *ii)* upon removal of the novel lincRNAs. Normalization was performed using the SCTrasform method, with default parameters, regressing out inherent variation caused by mitochondrial gene expression. Any other Seurat built-in function employed (i.e., RunPCA, FindNeighbors, FindClusters, and RunUMAP) was called with default parameters. For each experiment, correspondence between the two clustering solutions was tracked using the Jaccard similarity index [2]

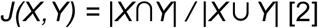

Clusters were ranked in descending order of J, and pairs were matched using a top-down approach, ensuring that the most similar clusters are prioritized for association. Novel clusters were considered those paired with J index ≤ 0.7. Finally, agreement between the two clustering solutions was computed using the Kendall rank correlation coefficient [3]

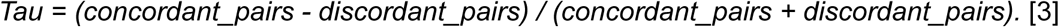

We further excluded 50 experiments with Tau ≤ 0.7, chosen to guarantee at least 85% of the cell-to-cluster assignments to be preserved. We argue indeed that lower consistency may reflect poorer performance of the experiment, and would unlikely be solely attributable to the expression of few novel lincRNAs, complicating downstream interpretation. Within each experiment, we further discarded clusters displaying extreme values of median mitochondrial (> 95th percentile) and MALAT1 expression (< 25th percentile for all tissues but blood, for which this filter was not applied). Additionally, we discarded 26 experiments for which ≤ 2 clusters were generated.

To identify novel lincRNAs potentially driving the generation of novel (sub)-clusters we employed FindAllMarkers to retrieve genes differentially expressed between each cluster, against the rest of the population (adjusted p-value < 0.05, and log_2_FC ≥ 1). To reduce the number of iterations, genes were tested only when expressed in at least 20% of the cells in the cluster under investigation. We kept markers detected in at least two experiments within the same tissue, and further excluded markers associated with clusters populated by ≤ 50 cells or to cluster fusions (defined as being paired more than once with moderate similarity: J ≥ 0.3). To ensure stringency, we report only markers ranking among the top 3,000 variable features in the associated experiment.

## Data availability

Raw PacBio and ONT reads are available through ArrayExpress accession E-MTAB-14562. A UCSC Genome Browser CLS track hub, with the read mappings and transcript models is available at https://hgwdev.gi.ucsc.edu/~markd/cls-stable-hub/.

## Code availability

Docker image with reference R packages used for analysis can be pulled at ghcr.io/guigolab/r4_gencode_phase3:latest.

The codes and links to the original files can be found at; https://github.com/guigolab/CLS3_GENCODE, doi: 10.5281/zenodo.13941033.

---

**Figure S1.**
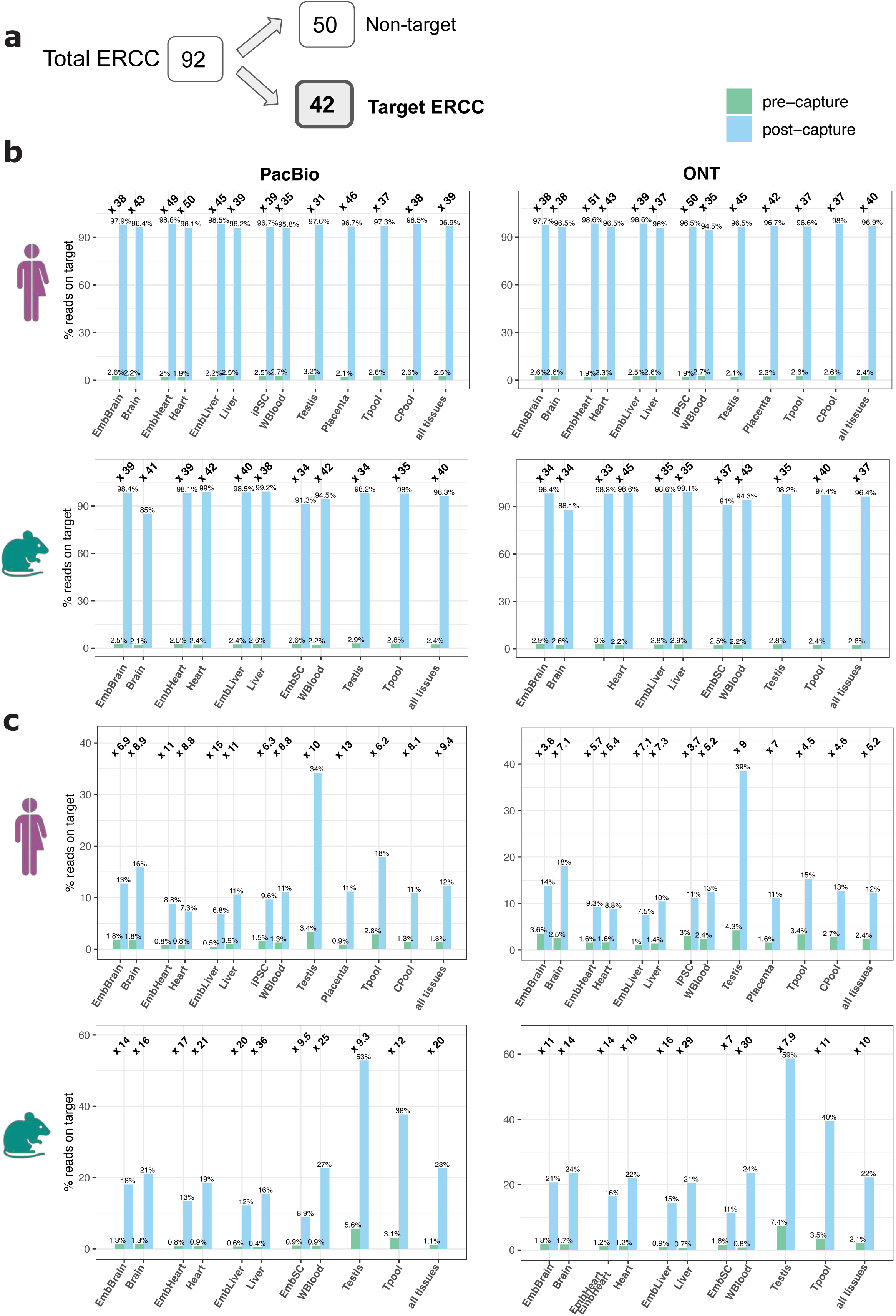
Capture on spike-ins. **A)** Spiked-in synthetic External RNA Control Consortium (ERCC) RNA sequences targeted in the capture design. Read enrichment for the **B)** targeted control ERCC sequences, and **C)** targeted regions across all catalogs post-capture. On the x axis, the samples are preceded by Emb when embryonic, as well as for visualisation purposes some have been contracted; WBlood (White Blood), Tpool (Tissue pool), and Cpool (Cell line pool).

**Figure S2.**
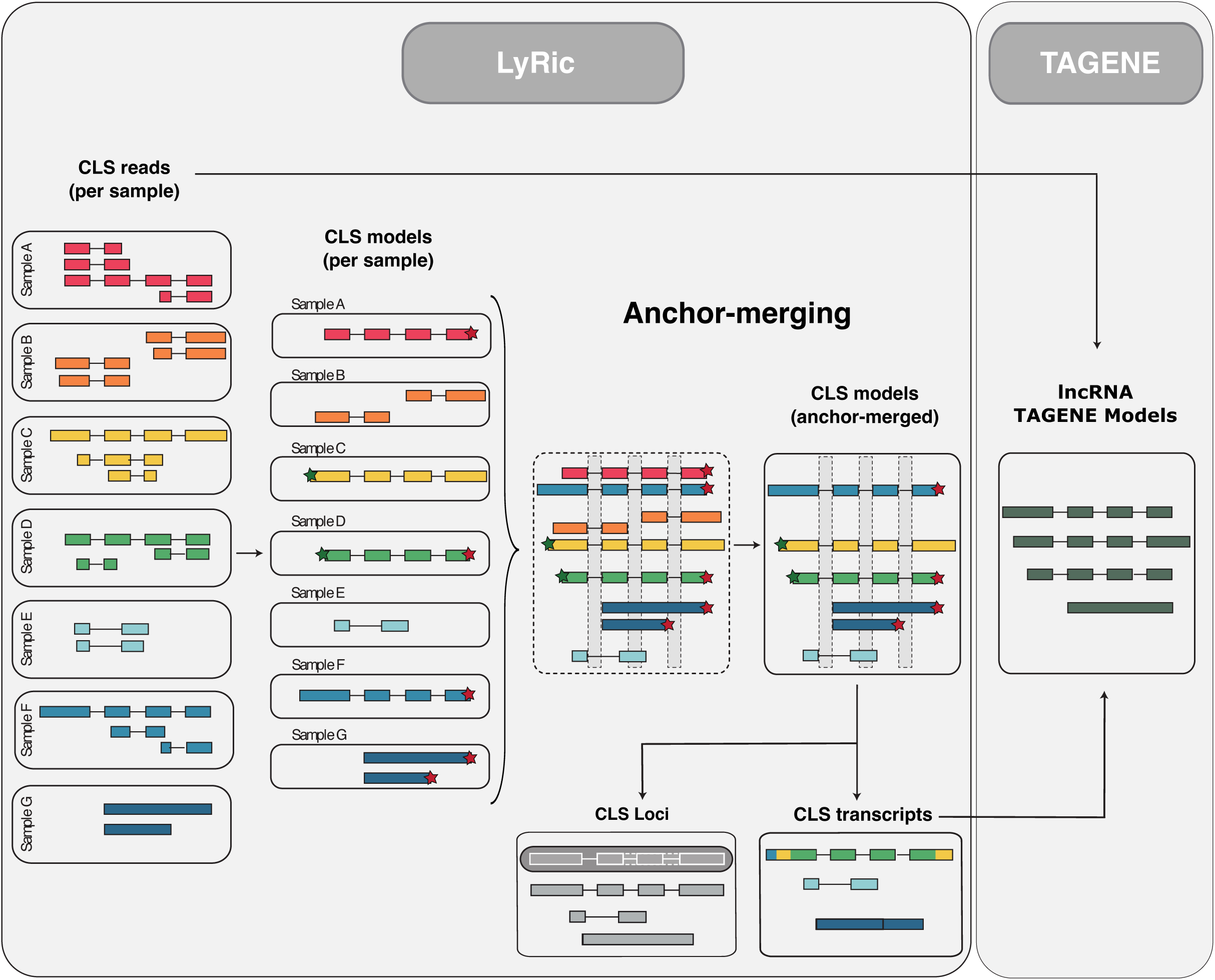
CLS model creation workflow. CLS reads were processed using LyRic to obtain high-confidence “CLS models’. These per-sample models were then “anchor-merged” across all tissues and developmental stages to obtain a comprehensive set of anchored models (**see Methods**), preserving the potential tissue specific ends usage. These anchor-merged CLS models were further collapsed irrespective of the supported ends to obtain “CLS transcripts” with reduced redundancy. These CLS transcripts together with the CLS reads were processed via TAGENE to have the final “TAGENE lncRNA models” incorporated into GENCODE. Based on the artefacts tags of TAGENE, CLS transcripts were filtered. The overlapping anchor-merged models are also clustered together based on same-strand exonic overlap into “CLS loci”.

**Figure S3.**
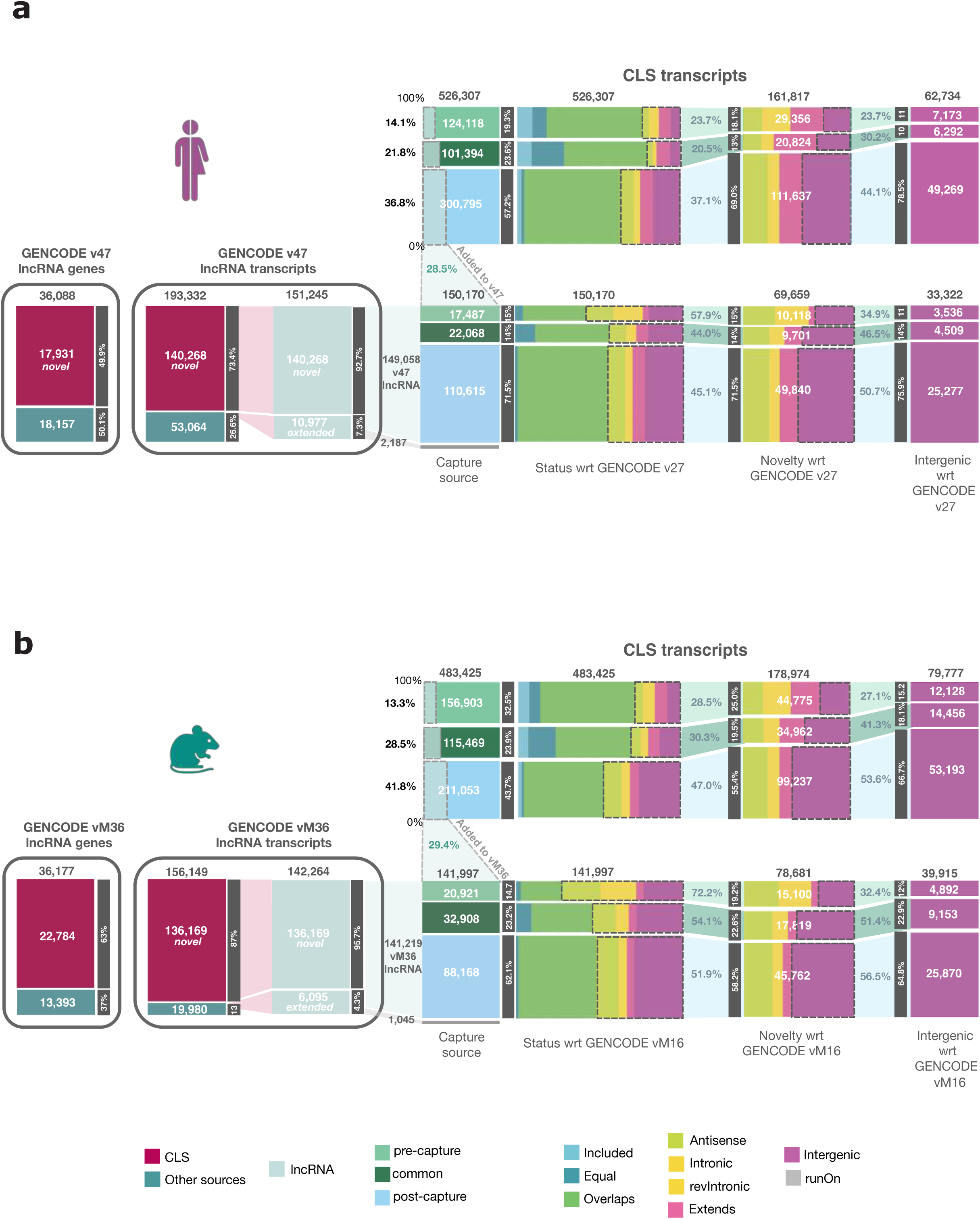
CLS transcripts summary. A detailed graphical classification for the obtained CLS transcripts (top-right panel) as well as those added to GENCODE (bottom-right panel) in **A)** human and **B)** mouse. The right panel shows the CLS transcripts grouped by capture source (the dark gray rectangles on the right depict the proportion obtained from each capture source with the topmost rectangle always depicting “pre-capture” design, the bottom one depicting “post-capture” design while the middle one depicting transcripts commonly obtained by both the capture sources; “common”), followed by further distribution of transcripts from each capture source based on the novelty status with respect to gencode v27. The dotted rectangles here highlight the novel transcripts with respect to v27. The shaded portions originating from these depict the proportion of novel transcripts relative to the total number for each capture source. The dark gray rectangles depict the proportion of novel transcripts from the respective capture source relative to all the novel transcripts. Further highlighted by dotted rectangles are transcripts intergenic with respect to v27. Again, the shaded regions originating from the novel transcripts show the proportion of those, from each capture source, that are intergenic. Finally, the right-most rectangles depict the intergenic transcripts only, with the gray rectangles showing the proportion across capture designs. In the lower panel, we show that 29% of CLS transcripts in both human and mouse are incorporated by GENCODE. These are distributed similarly based on capture source, and further based on novelty, with the most novel transcripts contributed by the post-capture design. The left-bottom panel of the figure shows the number of GENCODE transcripts and genes incorporated into v47 and vM36 thanks to CLS data, making up 93% and 96% of the lncRNAs transcripts, and 50 and 63% of lncRNA genes in v47 and vM36, respectively.

**Figure S4.**
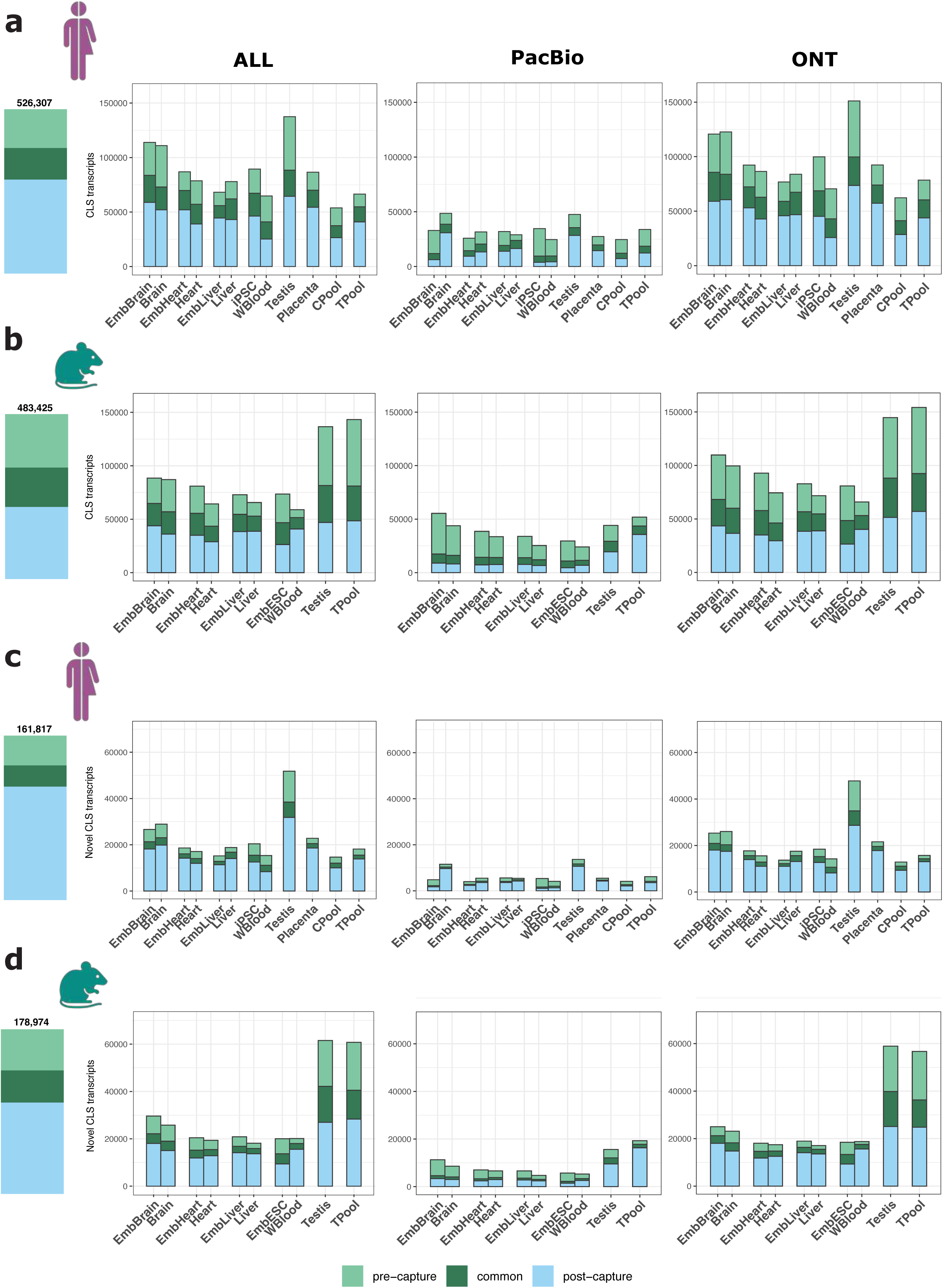
CLS transcripts yield across experiments. **A)** All CLS transcripts generated in human and **B)** mouse, across tissues and developmental stages. **C)** Novel CLS transcripts generated in human and **D)** mouse, across tissues and developmental stages. Bar plots (left) display the total number CLS transcripts obtained across all samples. The three panels on the right show (from left to right) the number of CLS transcripts obtained across “both PacBio and ONT” (ALL), as well as individually through “PacBio” and “ONT” sequencing platforms.

**Figure S5.**
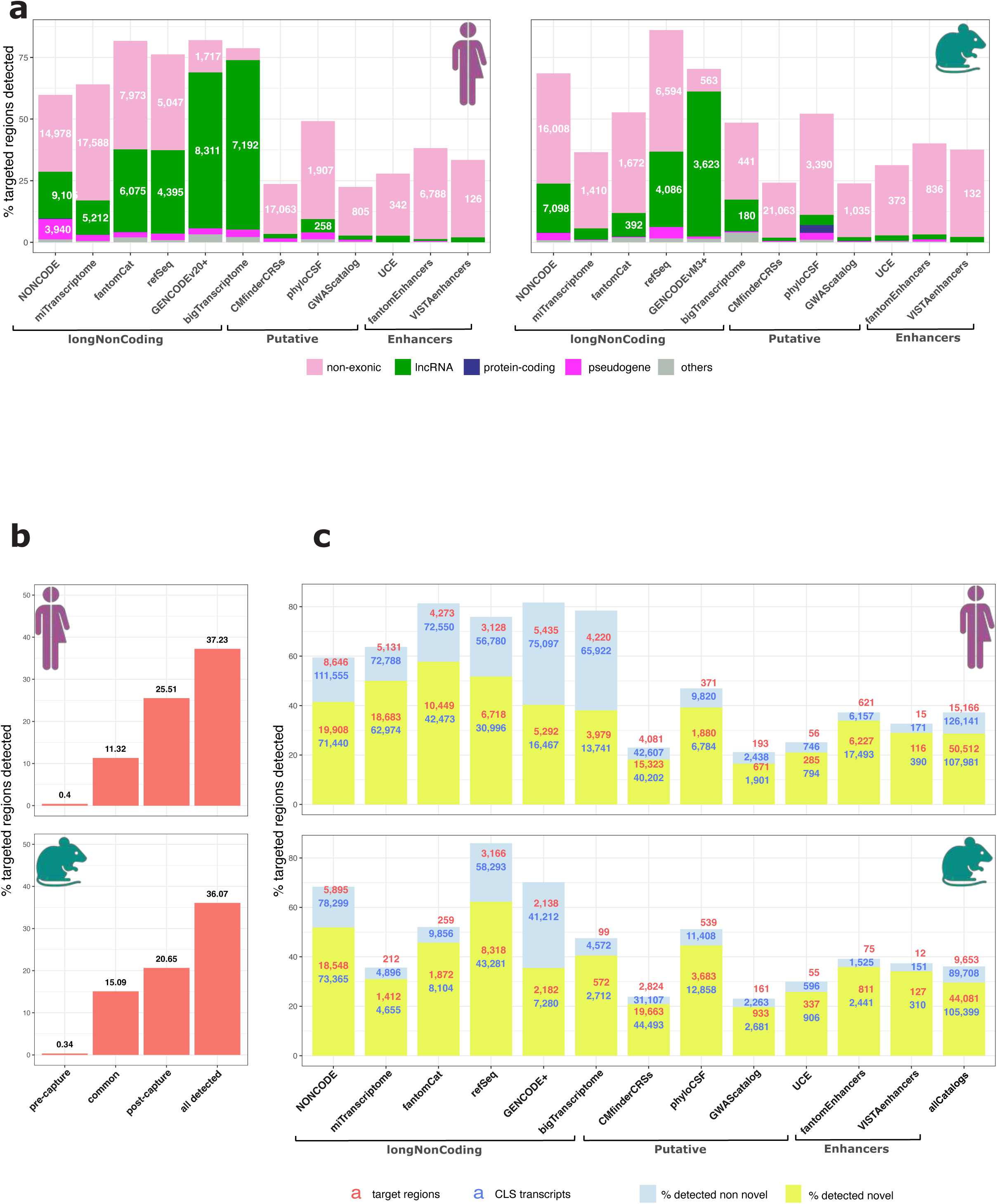
Targeted regions detected. **A)** Proportion of targeted regions detected and their biotype distribution along the human genome with respect to GENCODE v27 and GENCODE vM16 for human and mouse. **B)** Proportion of probed regions detected pre-capture, post-capture, and in both, and across all the experiments in human (top) and mouse (bottom). **C)** Proportion of targeted regions that help detect novel and known CLS transcripts in human (top) and mouse (bottom).

**Figure S6.**
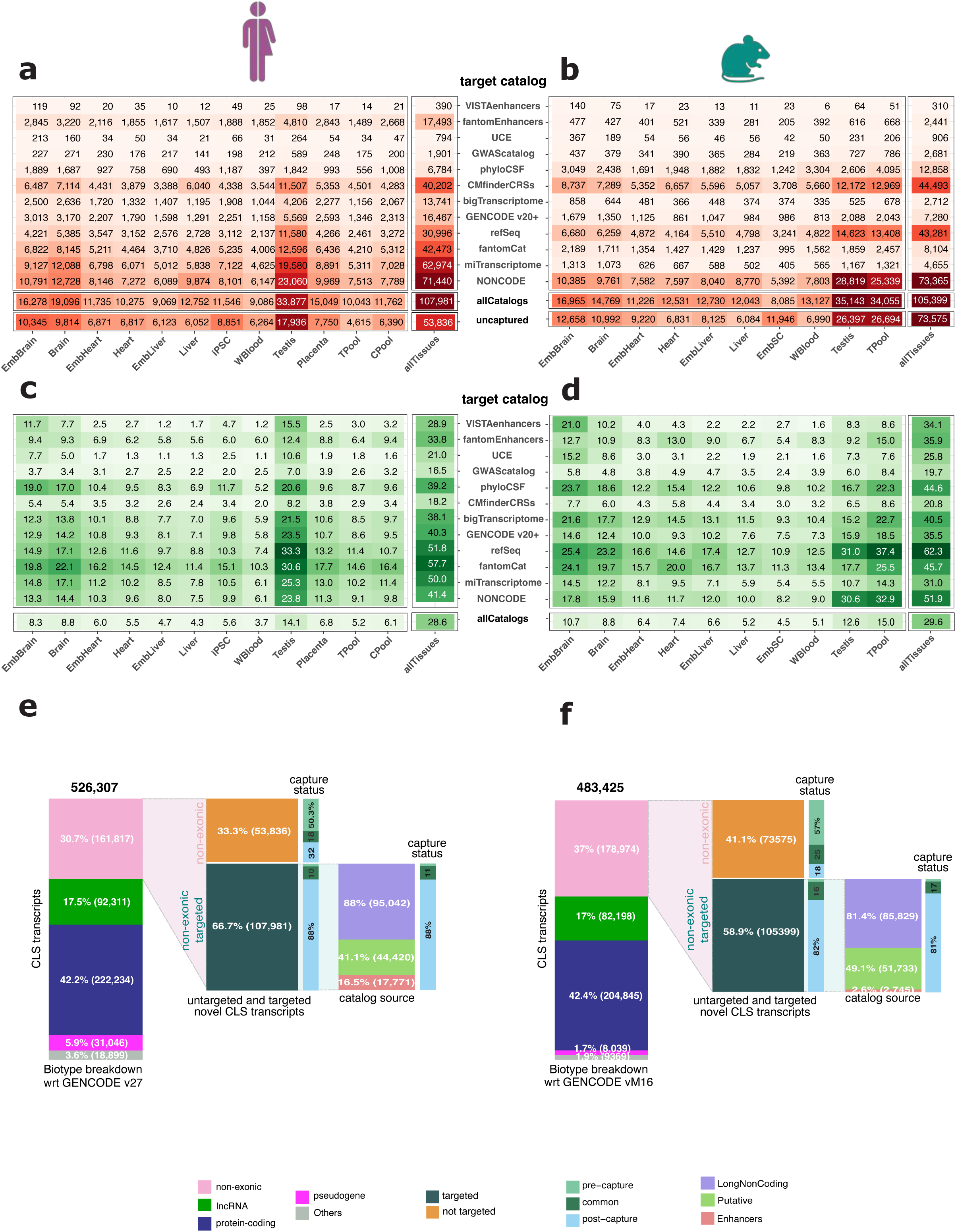
Novel CLS transcripts captured. Matrix depicting the number of novel CLS transcripts overlapping the targeted regions across each feature type and sample, as well as collated across samples and all the catalogs, in addition to the novel transcripts that did not have any overlapping probes in **A)** human and **B)** mouse. The proportion of target regions that help detect these novel transcripts are also reported in **C)** human and **D)** mouse. Barplot depicting the biotype breakdown for all the CLS transcripts with respect to v27 and vM16 for **E)** human and **F)** mouse. The non-exonic/novel transcripts are further divided based on whether these were obtained as a result of the captured regions or not, also depicting the experiment source.

**Figure S7.**
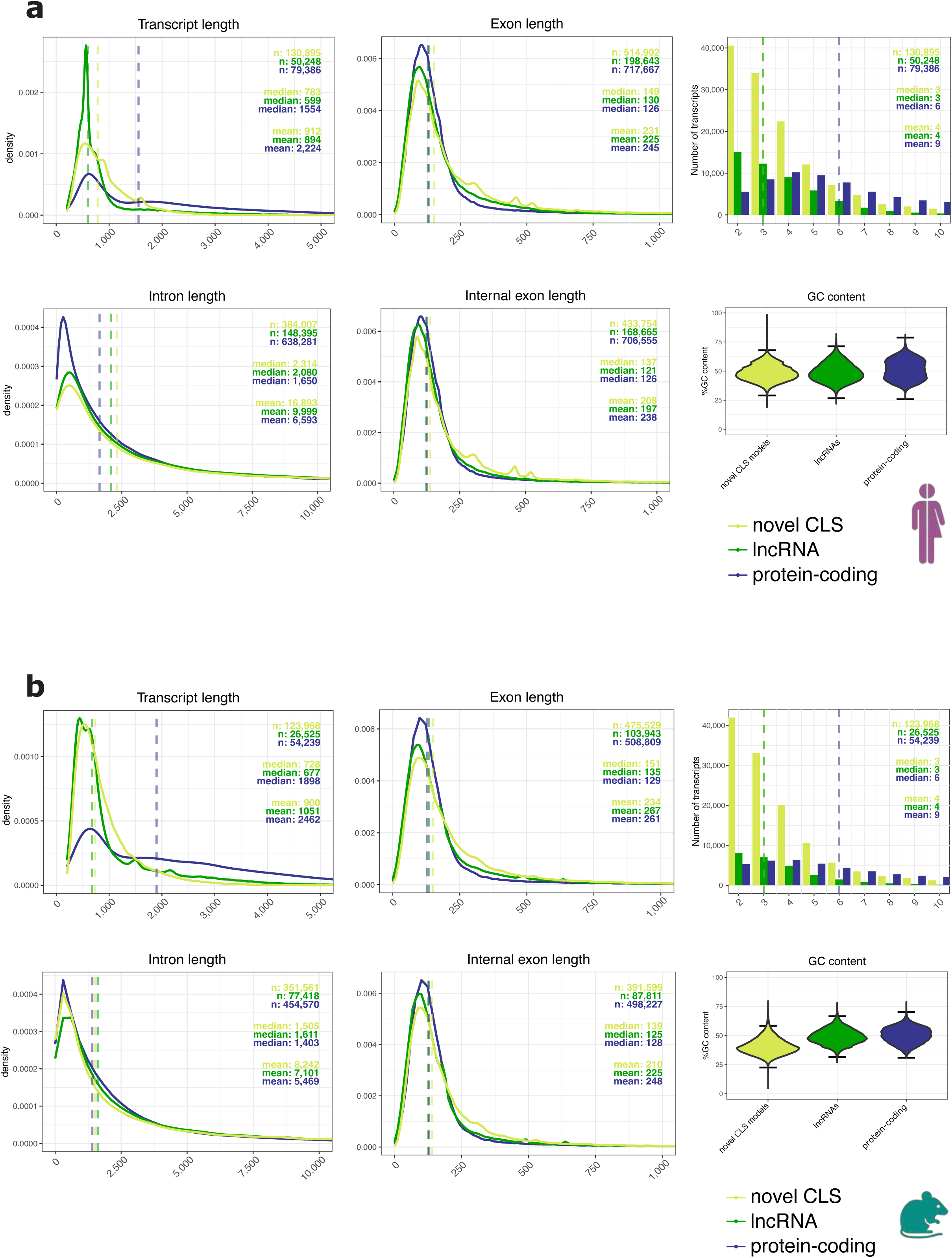
Distribution of lengths of transcripts, exons, introns, and number of exons. For **A)** human and **B)** mouse, the plots display the distribution of transcripts length, exons length and introns length, number of exons and GC content, compared across novel CLS transcripts, annotated lncRNAs, and protein-coding transcripts.

**Figure S8.**
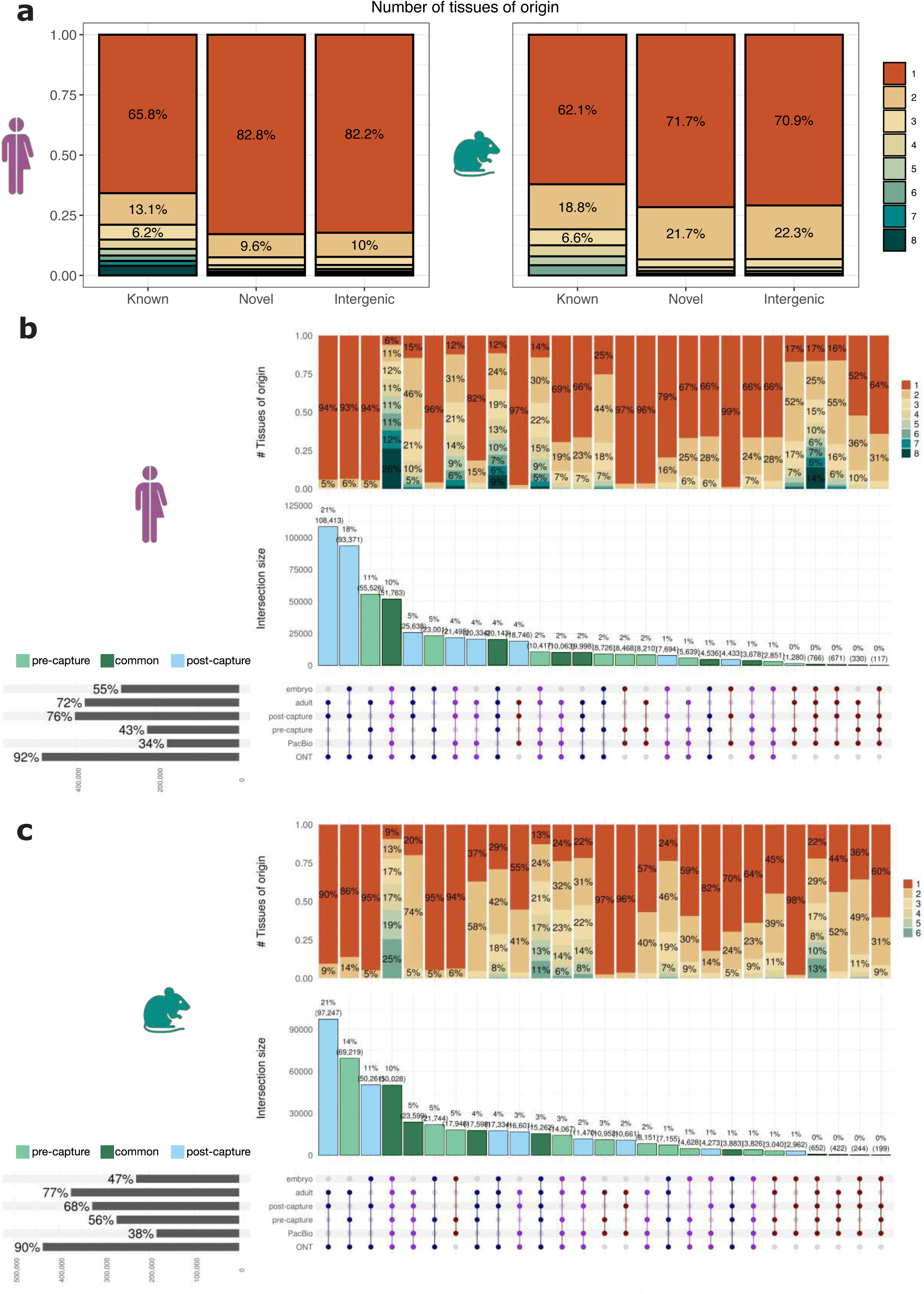
Classification of CLS models. **A)** Number of tissues in which CLS models are detected in human (left panel) and mouse (right panel), grouped by novelty status, as described in **Table S4**. Origin of CLS transcripts in human **B)** and mouse **C)** across development stage, capture protocol and sequencing technology. The upset plot shows the intersections across these categories; the dots are colored according to the technology of origin (whether unique to ONT, unique to PacBio, or detected in both), while the bars display the overlap of models between pre-capture and post-capture experiments. The barplot above shows the proportion of shared transcripts across tissues.

**Figure S9.**
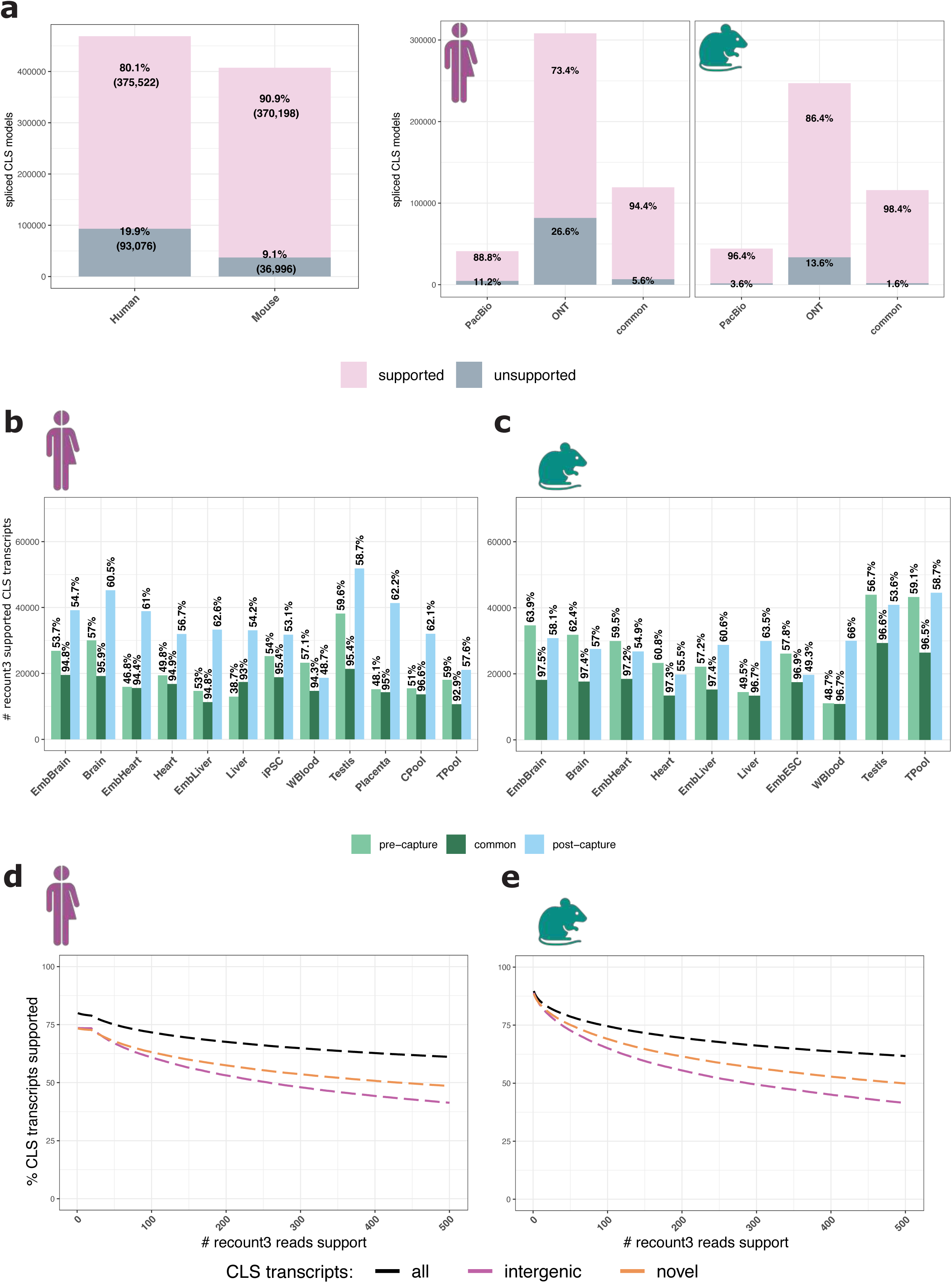
Recount3 support. **A)** Left panel: Proportion of spliced CLS transcripts supported by at least one recount3 exon-junction read in human and mouse. Right panel: recount3 support for spliced transcripts originating from PacBio only, ONT only and those obtained through both, for human and mouse. Proportion of recount3 supported spliced transcripts across all tissues, separated by the capture design, for **B)** human and **C)** mouse. Proportion of spliced CLS transcripts (y-axis) supported by increasing recount3 score support (x-axis) for **D)** human and **E)** mouse.

**Figure S10.**
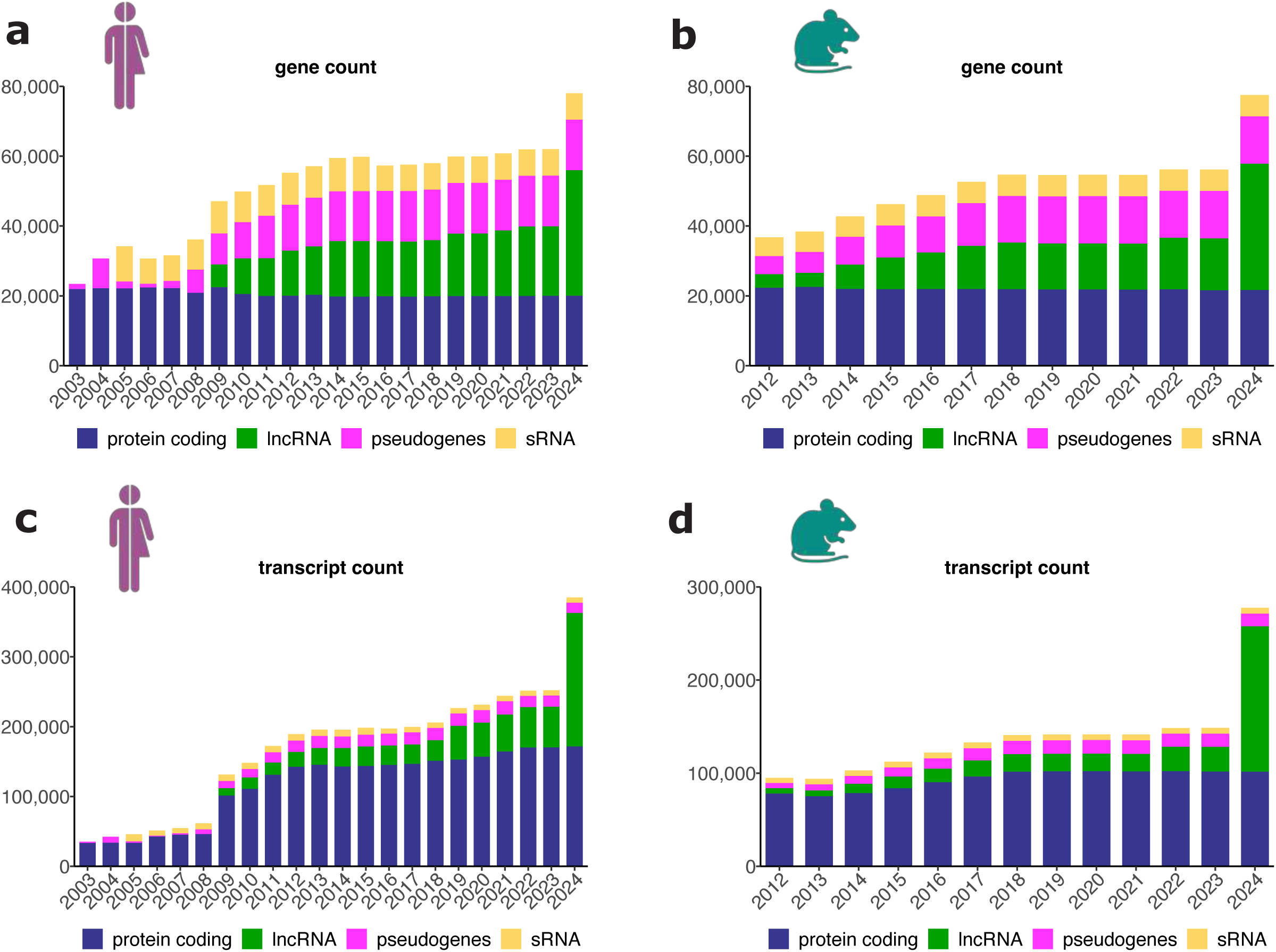
GENCODE annotation history. Numbers of genes and transcripts on primary assembly chromosomes in every year’s last GENCODE release in human **(A, B)** and mouse **(C, D)** broken down by broad biotype. IG/TR genes excluded.

**Figure S11.**
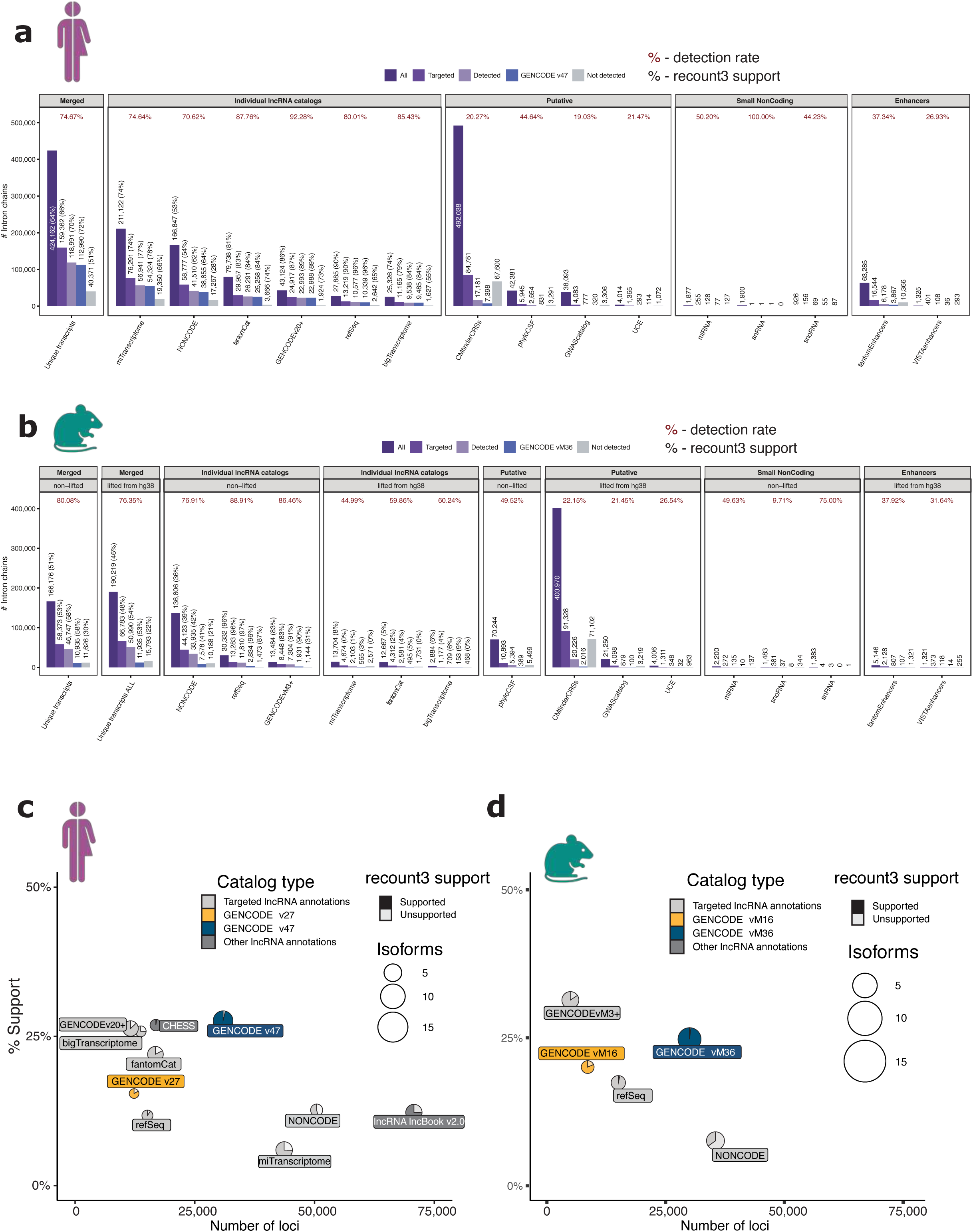
Targeting lncRNA catalogs. Number of intron chains from individual **A)** human and **B)** mouse datasets targeted by the CapTrap-CLS approach, categorized as targeted, detected, not detected, and incorporated into GENCODE, v47 and Mv36 respectively. The unique intron chains represent a non-redundant set compiled from all individual annotations. Percentages in brackets indicate the proportion of transcripts in each catalog with all splice-junctions supported by recount3 reads (i.e., each splice-junction has at least 50 supporting reads). Percentages shown at the top of the bars represent the detection rate; Comparison of **C)** human and **D)** mouse lncRNAs catalogs. On the x-axis, “Comprehensiveness”, represents the total number of gene loci, while on the y-axis, “Support” indicates the percentage of transcript structures whose TSS (±50 bases) is supported by a FANTOM CAGE cluster, and whose TTS includes a canonical polyadenylation motif within 10-50 bp upstream. Note that CAGE support is a poor metric to evaluate support for captured transcripts, as these are poorly represented in bulk RNA-Seq samples. Circle diameters show “exhaustiveness”, or the average number of transcripts per gene. Pie charts show the proportion of transcripts with all splice junctions supported by recount3 reads (with at least 50 reads). Only spliced models were included in this analysis. CLS transcripts here refer to transcripts identified using CapTrap-CLS, which are spliced, located on the reference chromosomes, and derived from individual lncRNA catalogs.

**Figure S12.**
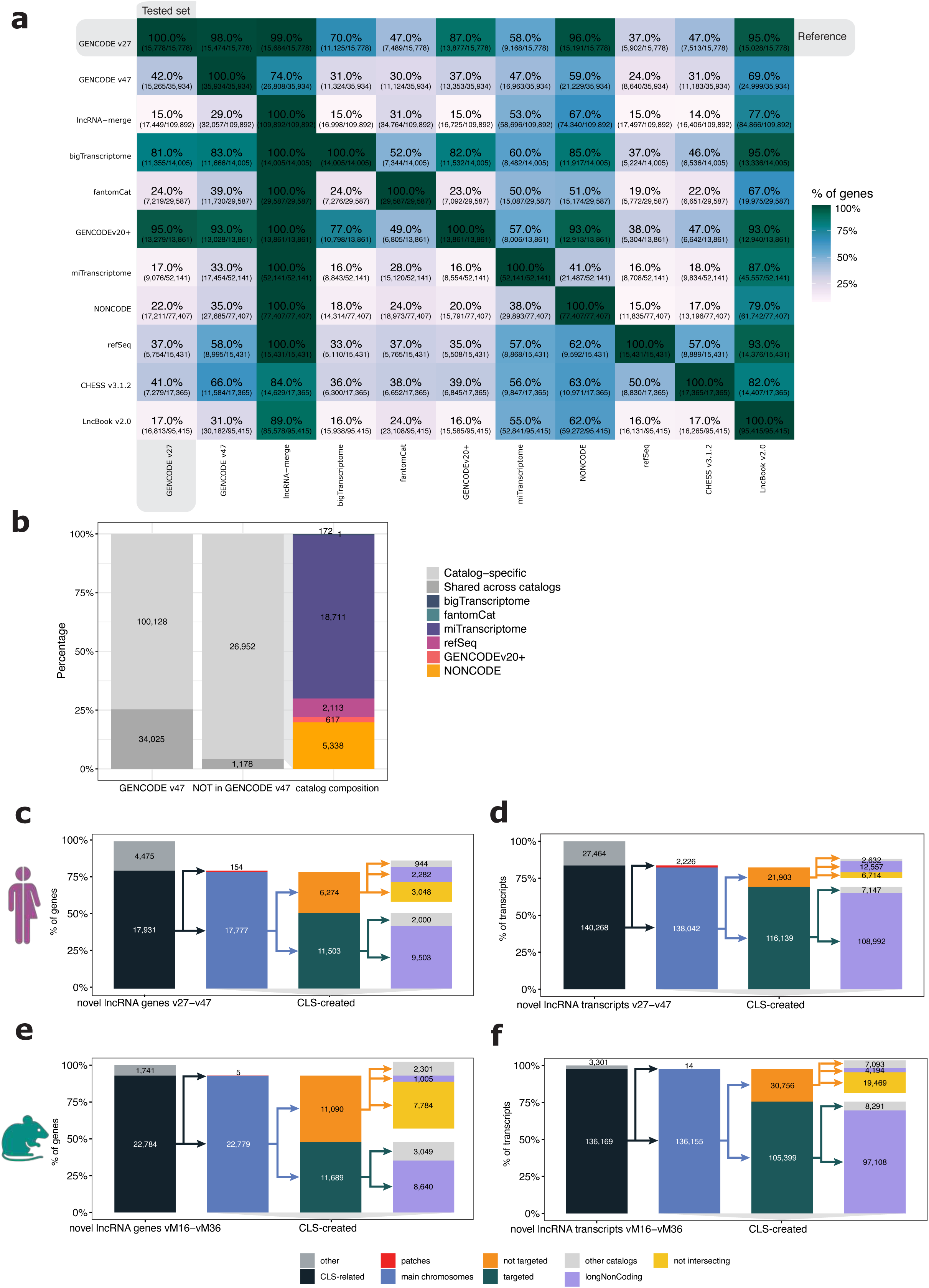
Expansion of the GENCODE lncRNA annotation compared to other lncRNA catalogs. **A)** Gene-level overlap between annotations, based on a strict definition. The values represent the percentage of gene loci in each row’s annotation that overlap with those in each column. Overlap is defined as a complete overlap of at least one gene span on the same strand. Both spliced and monoexonic genes are included in this analysis. **B)** Catalog specificity of lncRNA intron-chains detected (29.8% shared across catalogs; 69.2% catalog-specific) and not detected (12.1% and 87.9% respectively) by the CLS experiment. The origin of human (**C** and **D**) and mouse (**E** and **F**) genes and transcripts incorporated into GENCODE v47 and vM36, respectively. A gene or transcript is classified as originating from the lncRNA catalog if it is linked to at least one lncRNA target.

**Figure S13.**
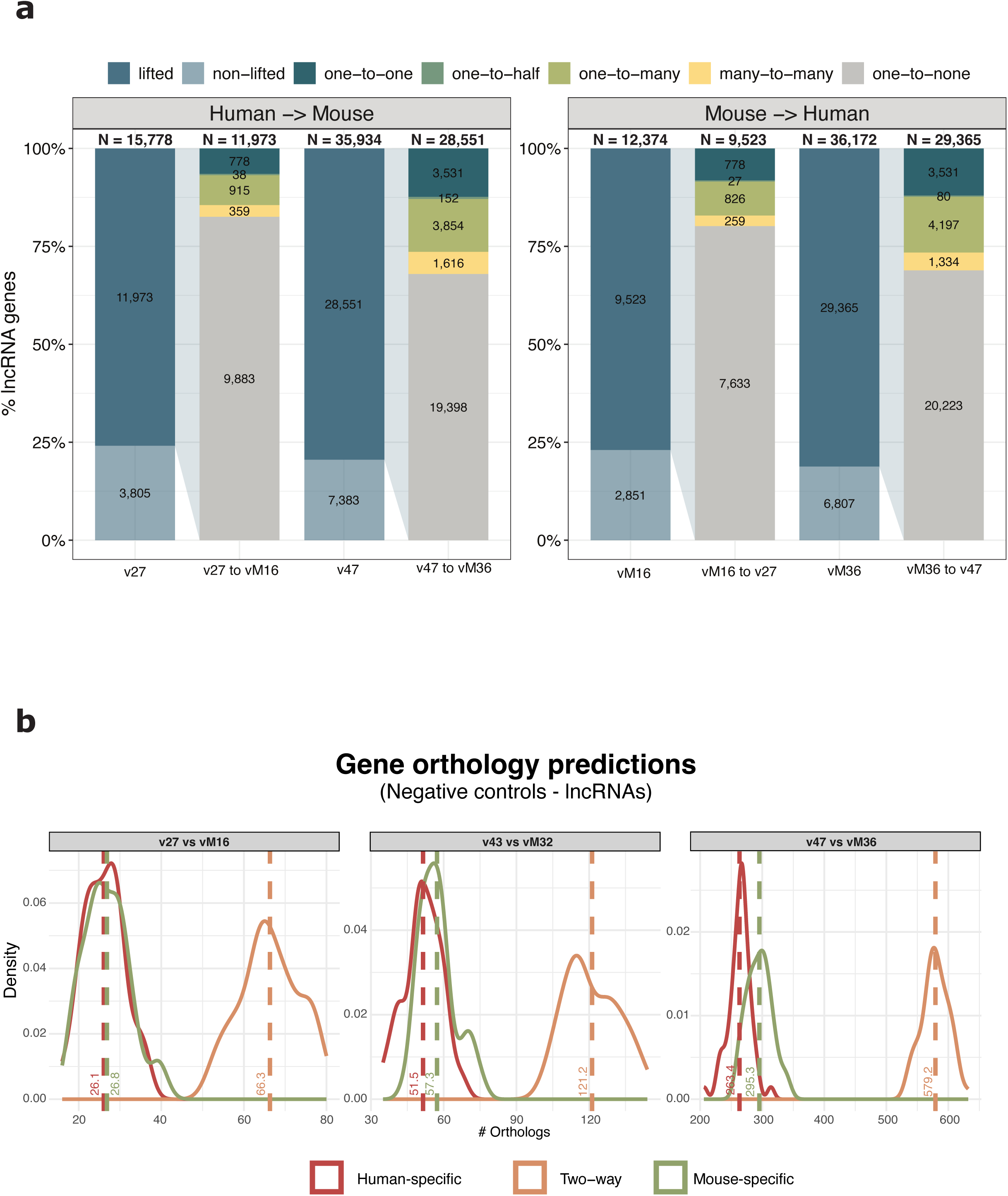
Detecting positionally conserved lncRNAs between human and mouse genomes. **A)** The number of orthologs in each orthology class. *One-to-half* are species-specific orthologs (one-way), that cannot be verified by reciprocal definition; **B)** Orthology predictions using negative controls generated through the shuffle GTF approach.

**Figure S14.**
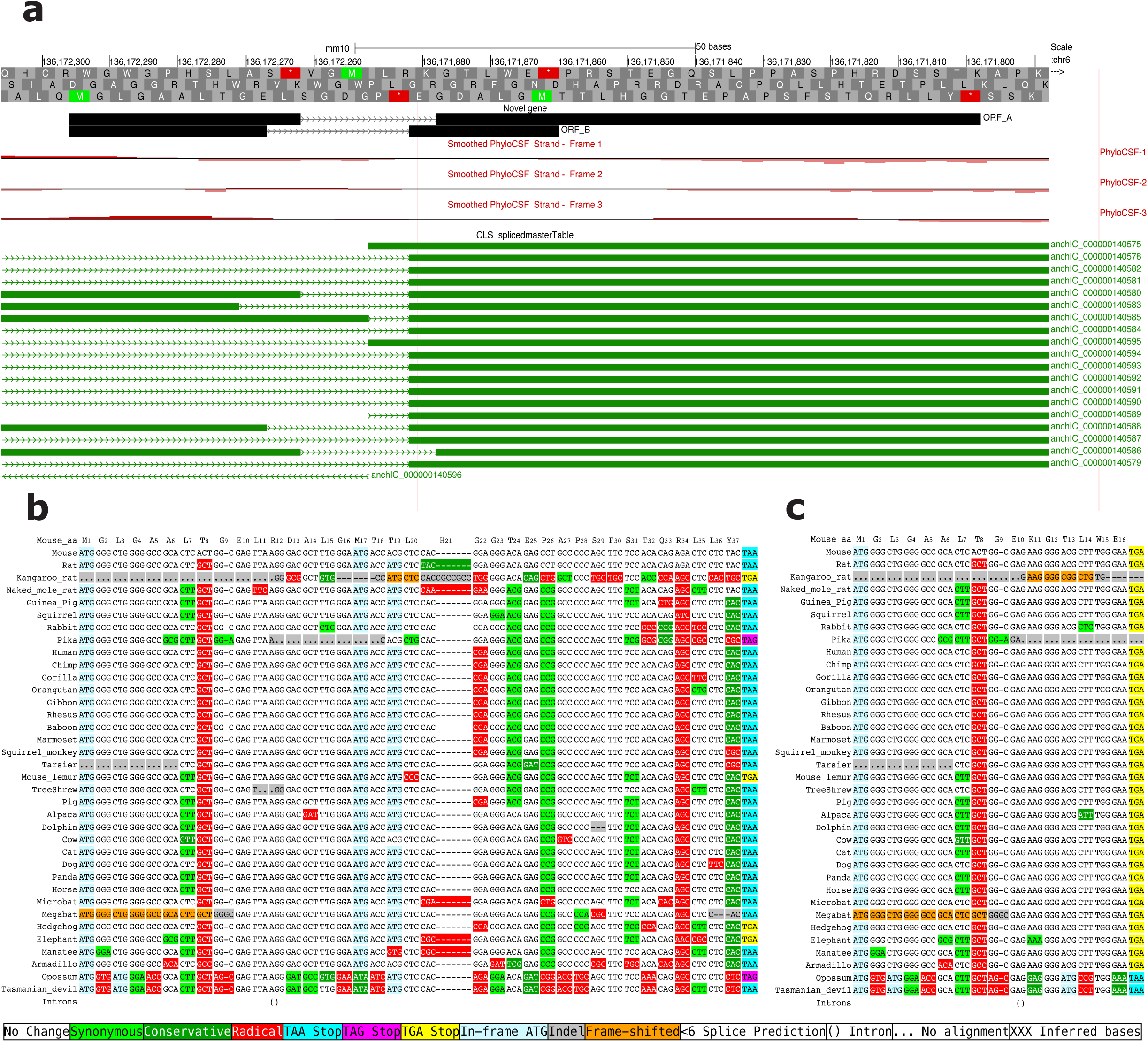
Novel 2-transcript protein-coding mouse gene found by searching CLS transcripts for regions with high PhyloCSF score not already annotated as protein-coding. The 37 and 16 amino acid transcripts share most of the first exon and overlap in different frames in the second exon. The human ortholog is also a novel protein-coding gene (not shown) but it is not contained in any CLS transcript. **A)** UCSC Genome Browser image showing the two transcripts, overlapping CLS transcripts, and PhyloCSF signal indicating evolutionary signature of conserved protein-coding DNA. **B)** and **C)** Mammal genome alignments of the two transcripts, rendered to show features indicative of protein-coding evolution, including frame conservation and predominance of synonymous substitutions (light green), by CodAlignView (https://data.broadinstitute.org/compbio1/cav.php).

**Figure S15.**
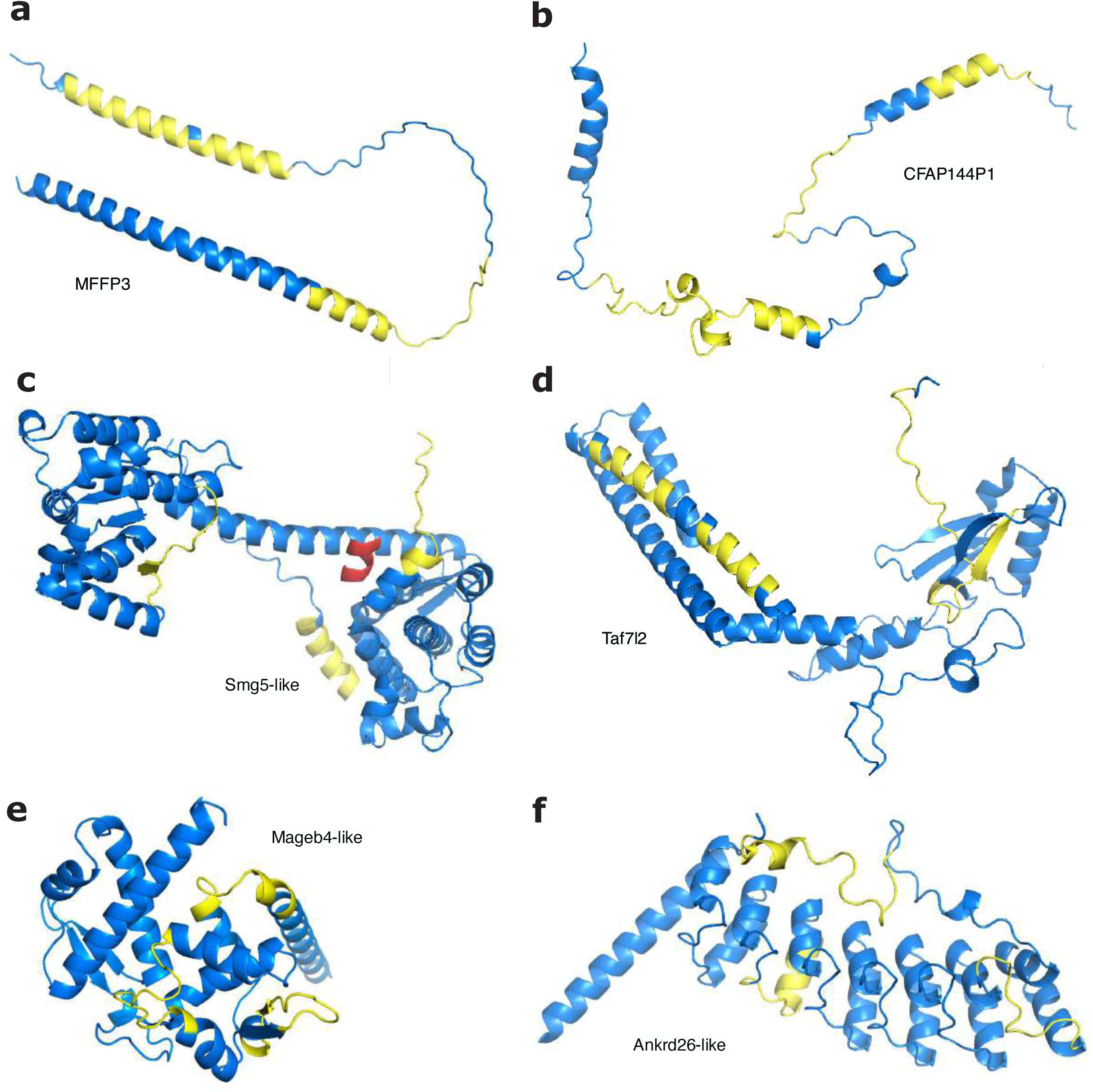
Possible coding regions for which we detected at least two non-overlapping peptides for transcripts in the human and mouse CLS analyses. Structures predicted using the HHPRED server or AlphaFold3. The detected peptides are mapped to the structures In yellow. **A)** Human predicted pseudogene MFFP3, expressed in testis and N-terminally truncated with respect to its parent. The ORF is also present in mouse, but it is even more truncated and would be a single helical protein. **B)** Human predicted pseudogene CFAP144P1, detected in testis and sperm and in higher quantities than the parent gene. **C)** A Smg5-like ORF in mouse with peptides detected in neural tissues. It is substantially different from the parent gene, but it conserves important functional residues. A Smg5 ligand (shown in red) has been mapped onto the model and the Smg5-like residues that would bind this ligand are highly conserved. **D)** Mouse Taf7l2, annotated as lncRNA in GENCODE and peptides detected in testis and epididymis. **E)** The globular domain of a mouse Mageb4-like protein, peptides detected in testis. **F)** The N-terminal domain of a mouse Ankrd26-like protein, peptides detected in testis and epididymis. We detected peptides for a novel Ankrd26-like protein in human too, but it is not clear whether the two genes are related.

**Figure S16.**
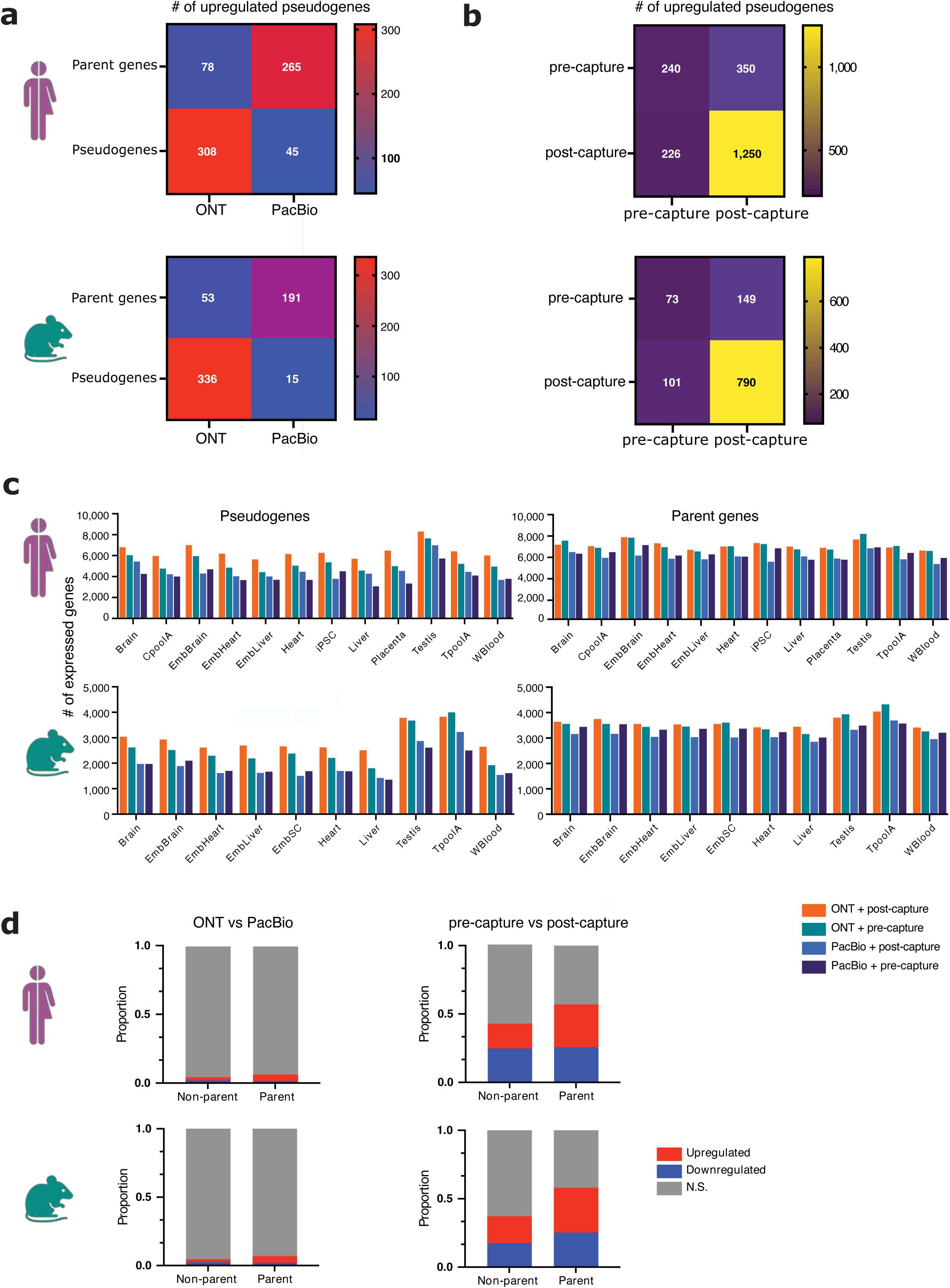
Comparison of the effect of sequencing technology and capturing on expression levels of pseudogenes and parent protein-coding genes. **A)** Heat map plot indicating the number of pseudogenes and parent genes that were upregulated when comparing the ONT and PacBio sequencing. **B)** Pairs of pseudogenes and parent genes that were upregulated in pre-capture or post-capture. **C)** Number of expressed pseudogenes and parent genes in various tissues, depending on the sequencing and capturing technology. **D)** and **E)** Bar charts indicating the proportion of significantly upregulated or downregulated parent and non-parent protein-coding genes when comparing **D)** ONT and PacBio sequencing and **E)** pre-capture and post-capture.

**Figure S17.**
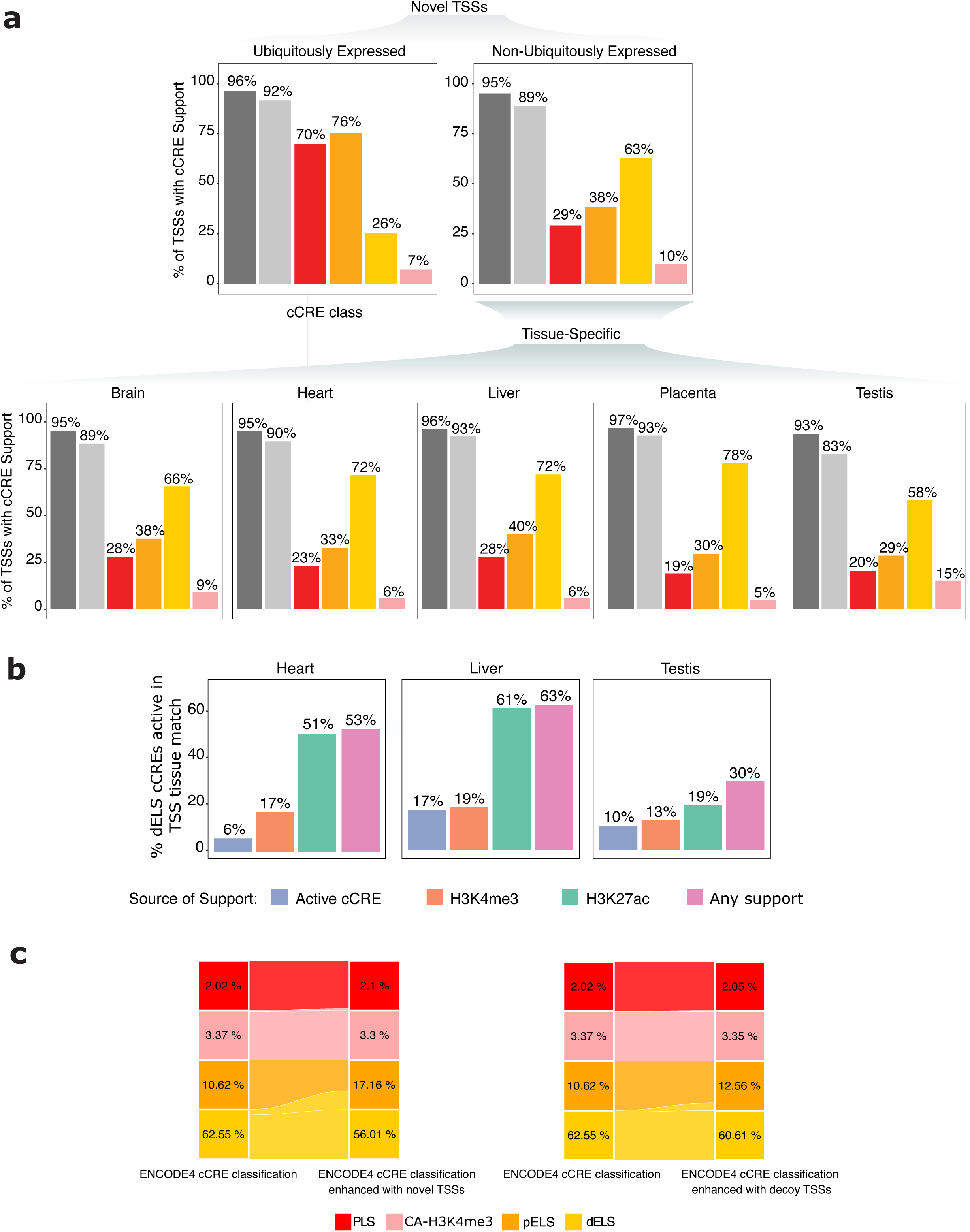
cCREs support for novel CLS TSSs. **A)** Bar plot showing the proportion of TSSs of novel CLS models supported by different types of cCREs, distinguishing between ubiquitously expressed and non-ubiquitously expressed TSSs. The type of cCRE is color-coded; “any class” includes additional types of cCREs not shown in the bar plot (CA-CTCF, CA-TF, CA, TF). In the lower panel, a similar representation focuses on tissue-specific TSSs across the five different tissues. **B)** Bar plot showing the proportion of dELS intersecting tissue-specific CLS transcripts characterized by chromatin activity in the same tissue as the corresponding TSS. “Active cCRE” means that the cCRE was attributed a category different than “low-DNase” in the corresponding tissue, “H3K4me3” and “H3K27ac” means that the TSS of the CLS transcript was found within 2kb from a peak of H3K4me3 or H3K27ac in the corresponding tissue. “Any support” is the union of active cCREs, H3K4me3, and H3K27ac-supported TSSs. **C)** Alluvial diagram showing the re-classification of TSS-proximity-dependent cCRE categories in the ENCODE registry. Two pairs of categories are shown *i)* PLS versus H3K4me3 marking in accessible regions (CA-H3K4me3), and *ii)* pELS versus dELS, which share the same histone marking signatures, but relying on different proximities to closest TSS (200bp and 2kb, respectively). The percentages indicate the proportion of cCREs from the entire registry that belong to each category in the original classification (left-side of either panel) compared upon enhancement with novel TSSs (left panel, right side) and **B)** decoy model TSSs (right panel, right side).

**Figure S18.**
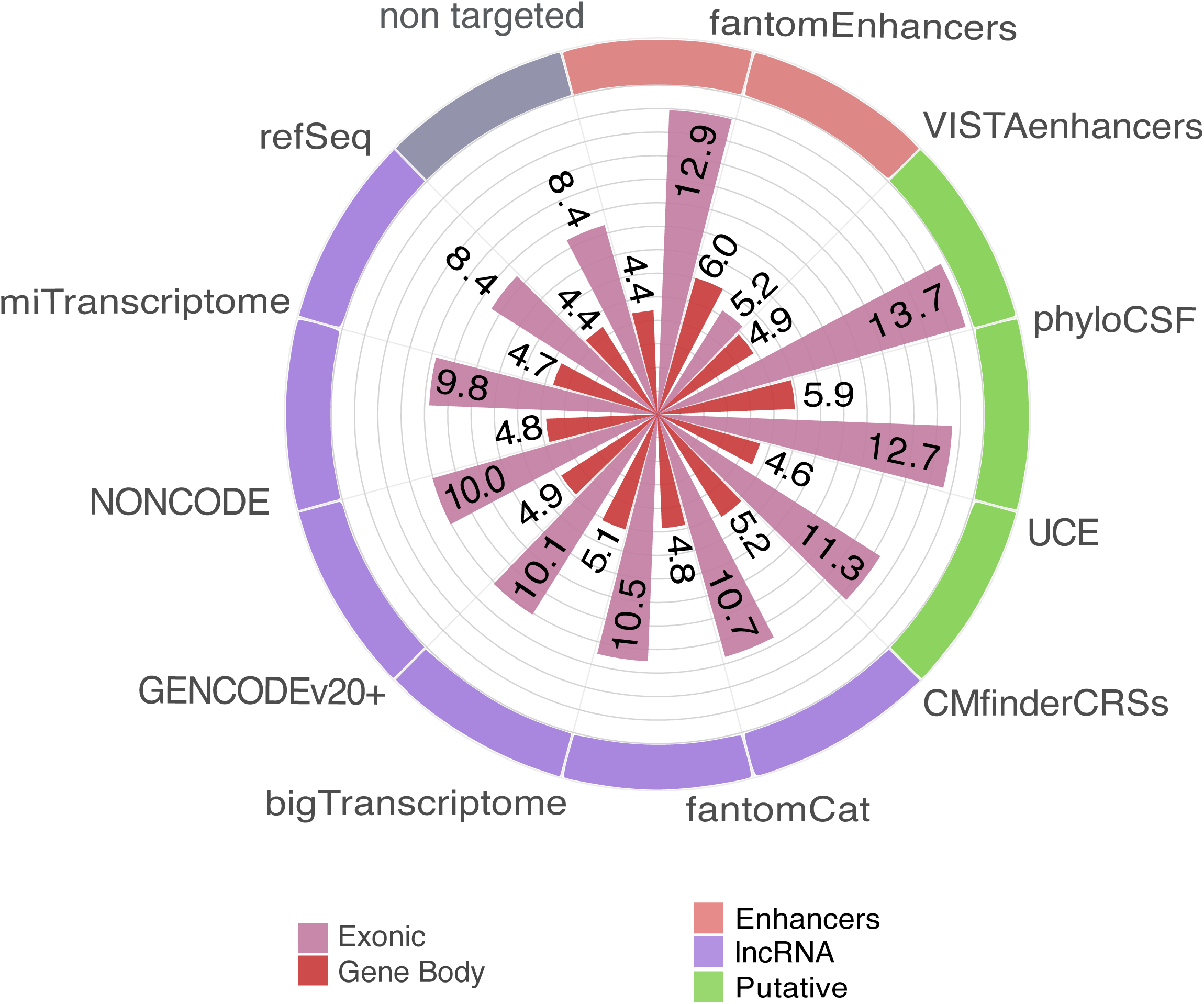
GWAS density in intergenic lncRNA transcripts. **A)** GWAS density (hits/100kb) computed along the exon projection of novel lincRNA transcripts across the different targeted catalogs, colored by targeted catalog. For visualization purposes, the frequencies for gene bodies (9.2) and exons (58.7) in transcripts captured via the GWAS catalog are not reported. When dissecting the contribution of the different targeted catalogs, novel lincRNAs targeted via phyloCSF, and regions predicted to encode structural RNAs, are among the ones showing the highest density. Remarkably, these are the type of elements in which single nucleotide disruptions are likely to have the larger functional impact.

**Figure S19.**
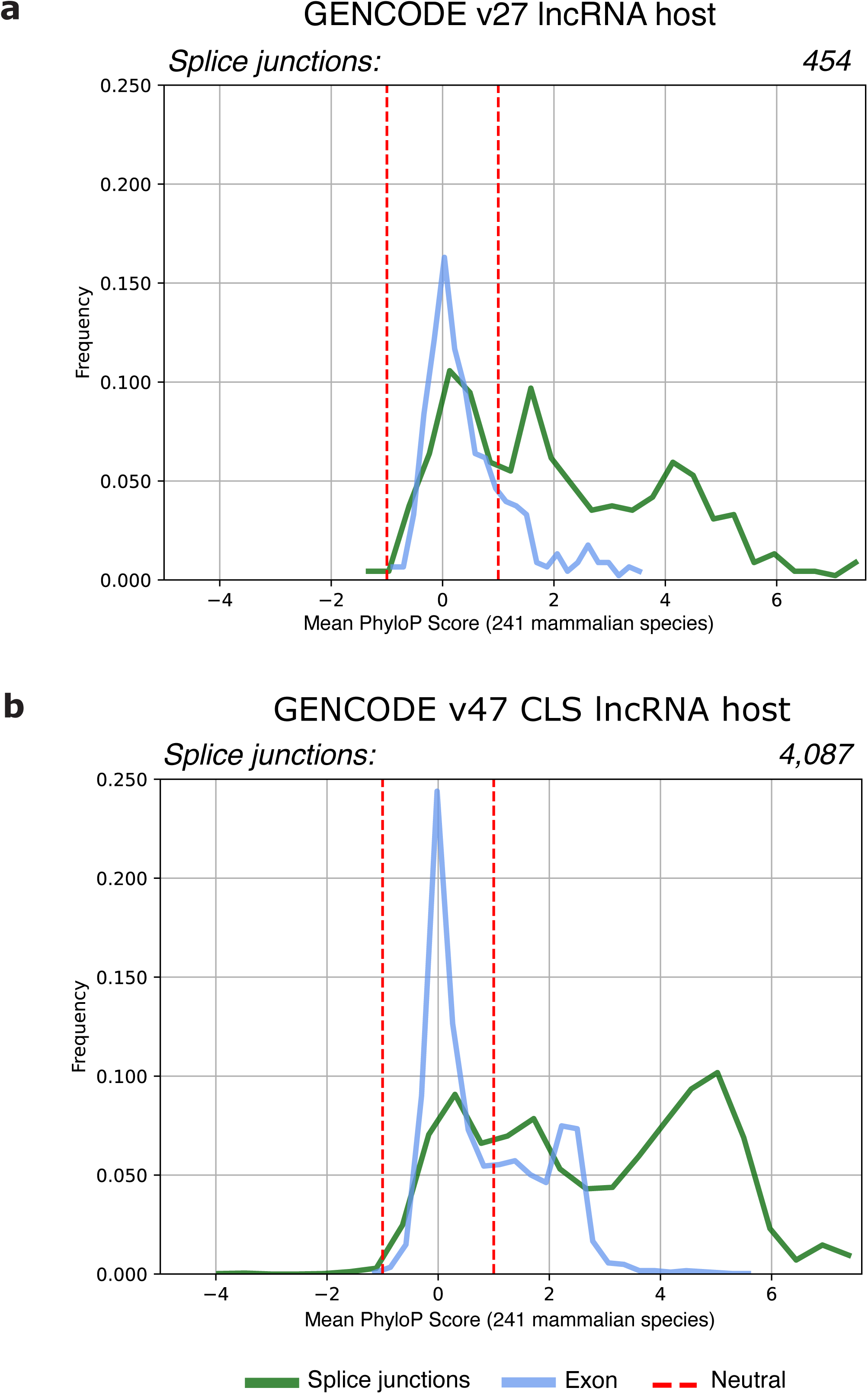
Novel host genes of small RNAs. Frequency of per-transcript exon and splice-junction mean PhyloP scores for lncRNAs hosting miRNAs or snoRNAs outside protein-coding loci. The dashed red lines indicate the range considered under neutral selection. **A)** 454 GENCODE v27 host lncRNA transcripts with 20% of exons and 64% of the splice-junctions classified as conserved, and 71% and 28% respectively neutral-evolving **B)** 4,087 GENCODE v47 host lncRNA transcripts derived from novel lincRNAs with 39% of exons and 74% of the splice-junctions classified as conserved and 74% and 23% respectively neutral-evolving.

**Figure S20.**
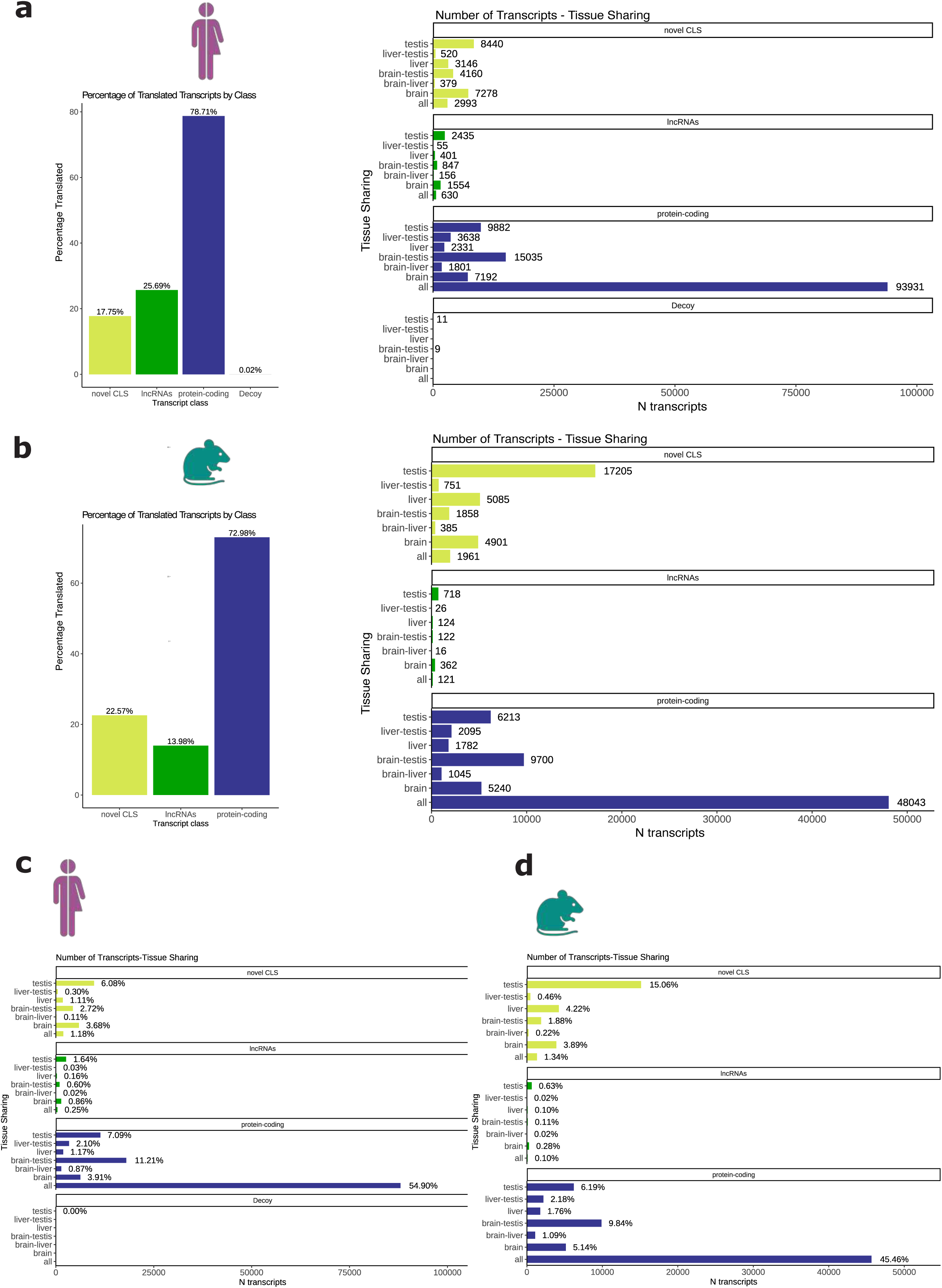
Non-canonical ORF (ncORFs) in novel lincRNAs. **A)** Human data. Left: Percentage of translated transcripts (i.e., transcripts containing ncORFs) by class: novel lincRNA transcripts (26,916, 17.8%), lncRNAs (6,078, 25.7%) and protein-coding genes (133,810, 78.7%). Right: Number of translated transcripts per class and tissue in which translation is detected. Testis means only in testis, testis-brain only in testis and brain, all in the three tissues including liver. **B)** Mouse data. Left: novel lincRNA transcripts (32,146, 22.6%), lncRNAs (1,489, 14%) and protein-coding genes, (74,118, 73%). Right: Number of translated transcripts per class and tissue in which translation is detected. Subsampling by read number for **C)** human and **D)** mouse Top: Percentage of translated transcripts per class in human after subsampling. Bottom: Percentage of translated transcripts per class in mouse after subsampling. Tissue specificity of translated ncORFs is much larger in testis and brain than in liver in both species. Moreover, the number of novel lincRNA transcripts exclusively translated in testis is much larger in mouse than in human.

**Figure S21.**
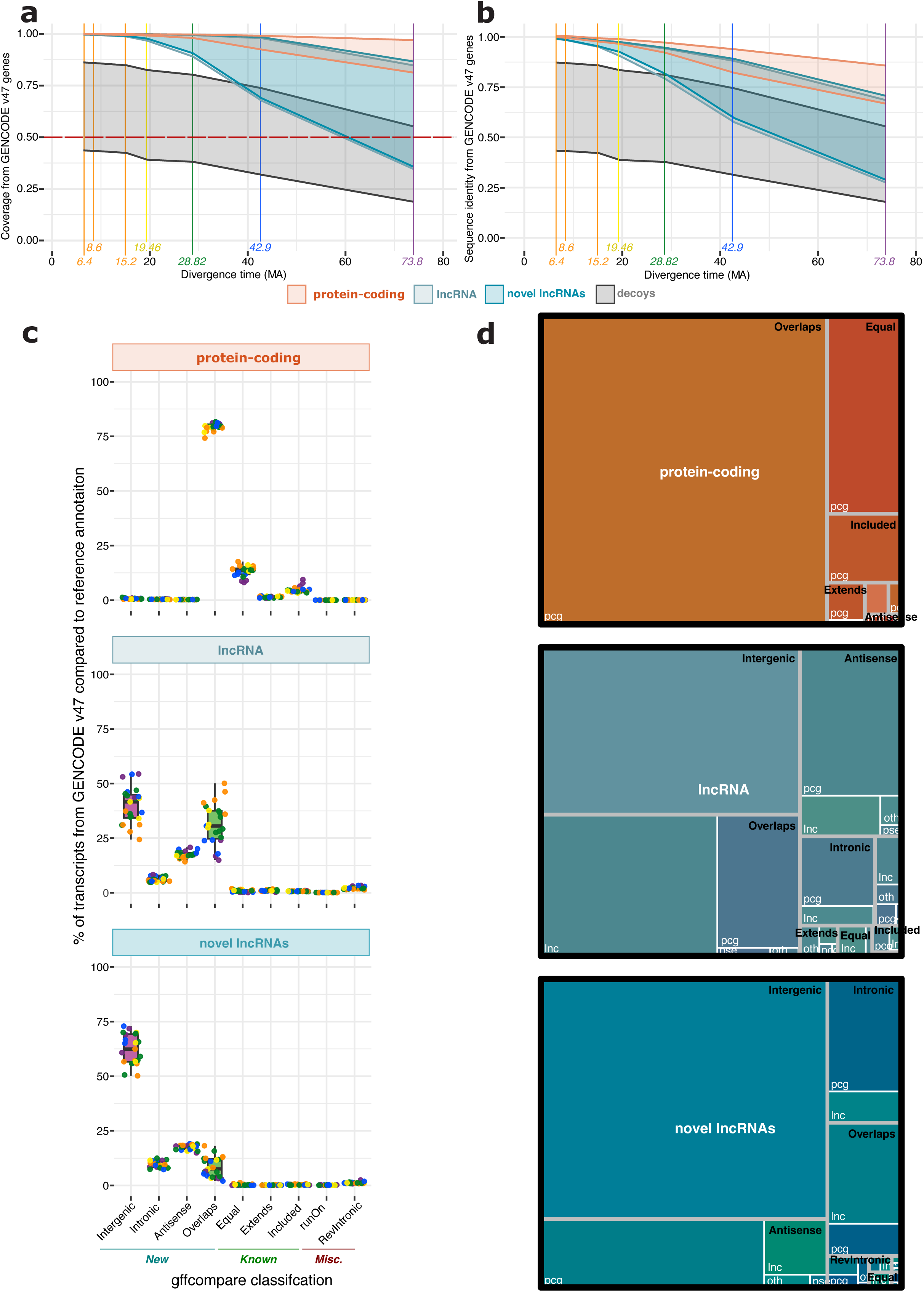
Characterization of lifted GENCODE v47 transcripts across 32 primate species. **A)** Coverage and **B)** sequence identity of protein-coding genes, annotated lncRNAs, novel lncRNAs, and decoy models lifted from GENCODE v47 to 32 primate species across 5 phylogenetic groups (colored lines). For each biotype, the upper and lower lines represent the medians of the 3rd and 1st quartiles, respectively, for all species at a given divergence time. The dashed red lines indicate the thresholds used to classify genes as successfully lifted. **(C)** Classification of lifted protein-coding and lncRNA transcripts from GENCODE v47 based on their correspondence to existing transcripts annotated in RefSeq for each primate species. Each point represents a species, colored according to its divergence time from human. **(D)** Treemap visualization of the classification categories defined in (C), showing how lifted transcripts relate to the biotypes present in the RefSeq primate annotations. Grey boxes indicate the classification types, while white boxes correspond to the assigned biotypes in the reference annotations.

**Figure S22.**
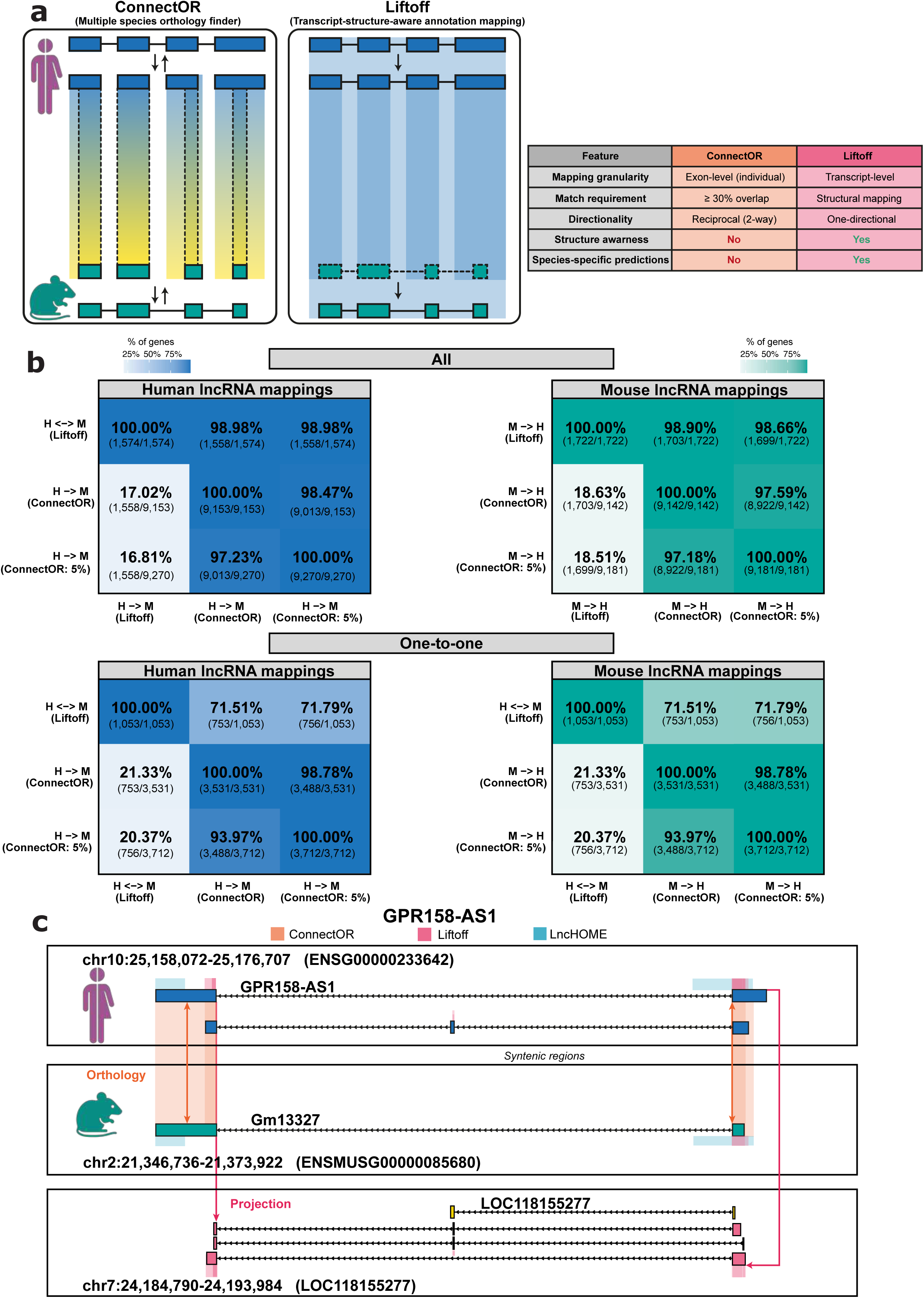
Comparative Analysis of ConnectOR and Liftoff. **A)** Diagram showing the ConnectOR and LiftOff workflows. Human genes are shown in blue and mouse genes in green, with shaded regions indicating mapped areas. **B)** Overlap between ConnectOR lncRNA orthologs and Liftoff mappings. **C)** Performance of ConnectOR and LiftOff in mapping the human *GPR158-AS1* lncRNA gene to mouse and common marmoset (*Callithrix jacchus*). Pink and orange shades represent regions identified by ConnectOR and LiftOff, respectively, while yellow highlights indicate regions detected in mouse by the alternative method for identifying syntenic lncRNAs enriched with RNA-binding protein (RBP) binding site patterns (coPARSE-lncRNAs).

**Figure S23.**
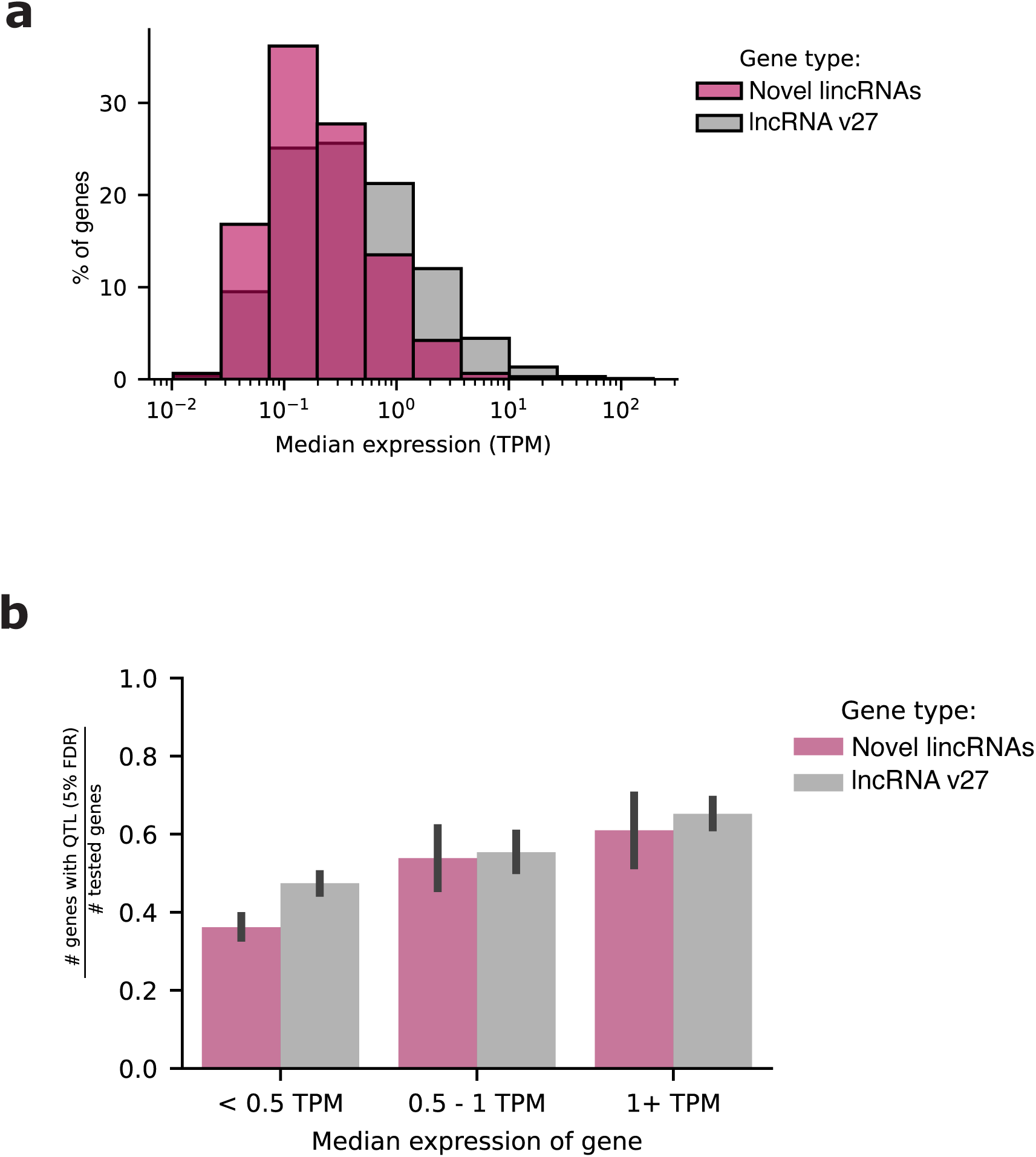
**A)** Percentage of genes in each median TPM bin across samples for novel lincRNAs and annotated lncRNAs (v27). **B)** The proportion of novel lincRNAs and annotated lncRNAs found as eQTLs for those expressed at a median TPM of < 0.5 TPM, 0.5 - 1 TPM, and > 1 TPM. Tested genes for eQTLs are those sufficiently expressed (see Methods). Error bars represent 95% confidence interval.

**Figure S24.**
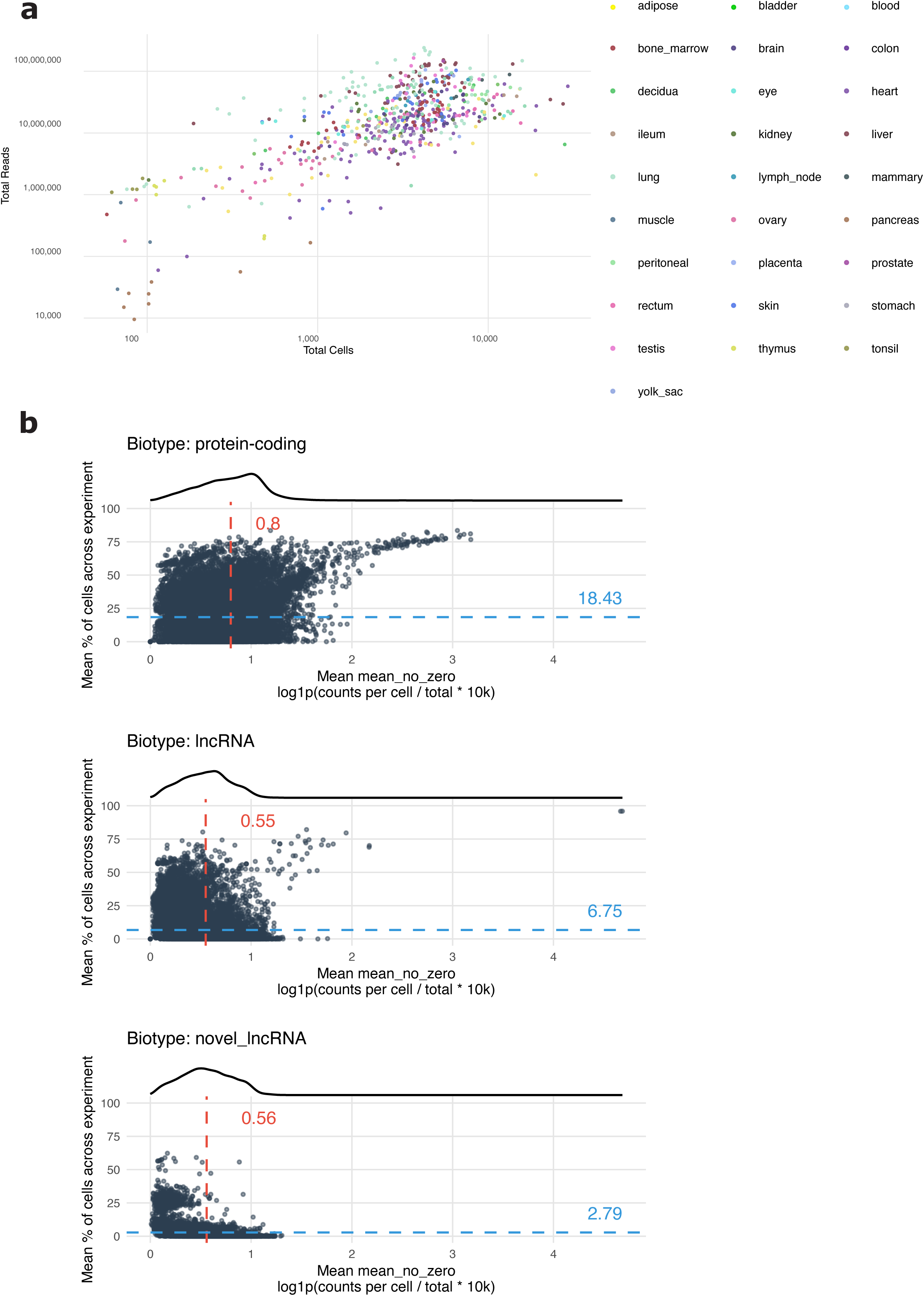
**A)** Summary of the single-cell and single-nuclei experiments requantified in this study, colored according to the tissue of origin as reported in the Human Commons Cell Atlas. Total number of cells is intended after stringent filtering applied (see Methods). Total number of reads is the total sum of raw counts, in the filtered cells. **B)** Summary of expression of genes, split per biotype, across the single-cell and single-nuclei experiments requantified in this study. For each experiment, the mean of the scaled counts across cells is obtained for each gene, excluding zeros. On the x axis is displayed the average of the mean obtained across experiments for each gene. The y axis shows the average proportion of cells where the gene has been detected (scaled counts > 0).

**Figure S25.**
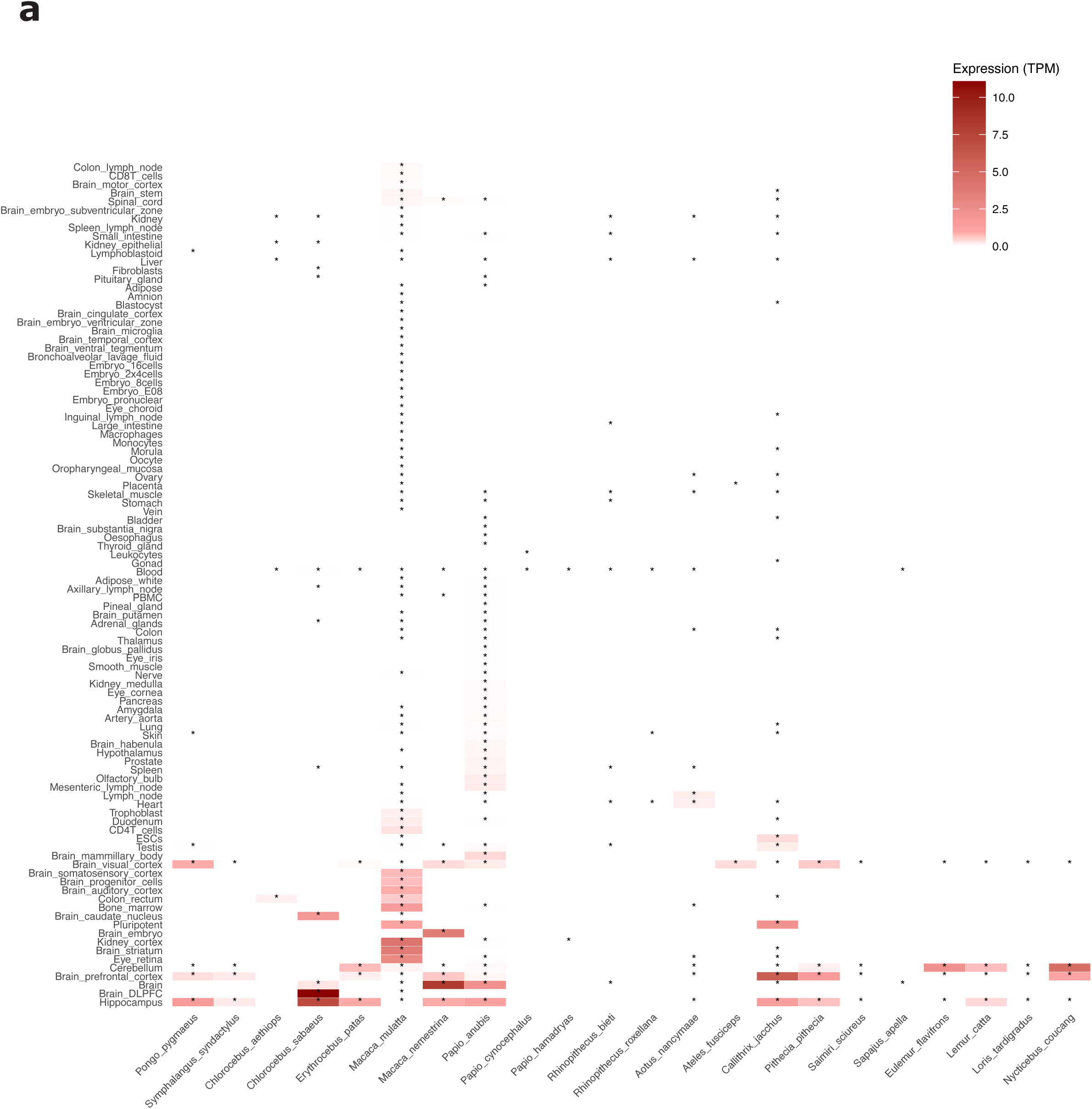
Expression (in TPM) across different primates species (phylogenetically ordered), for the tissues requantified in this study. An asterisk marks the cases where the data was collected.

**Figure S26.**
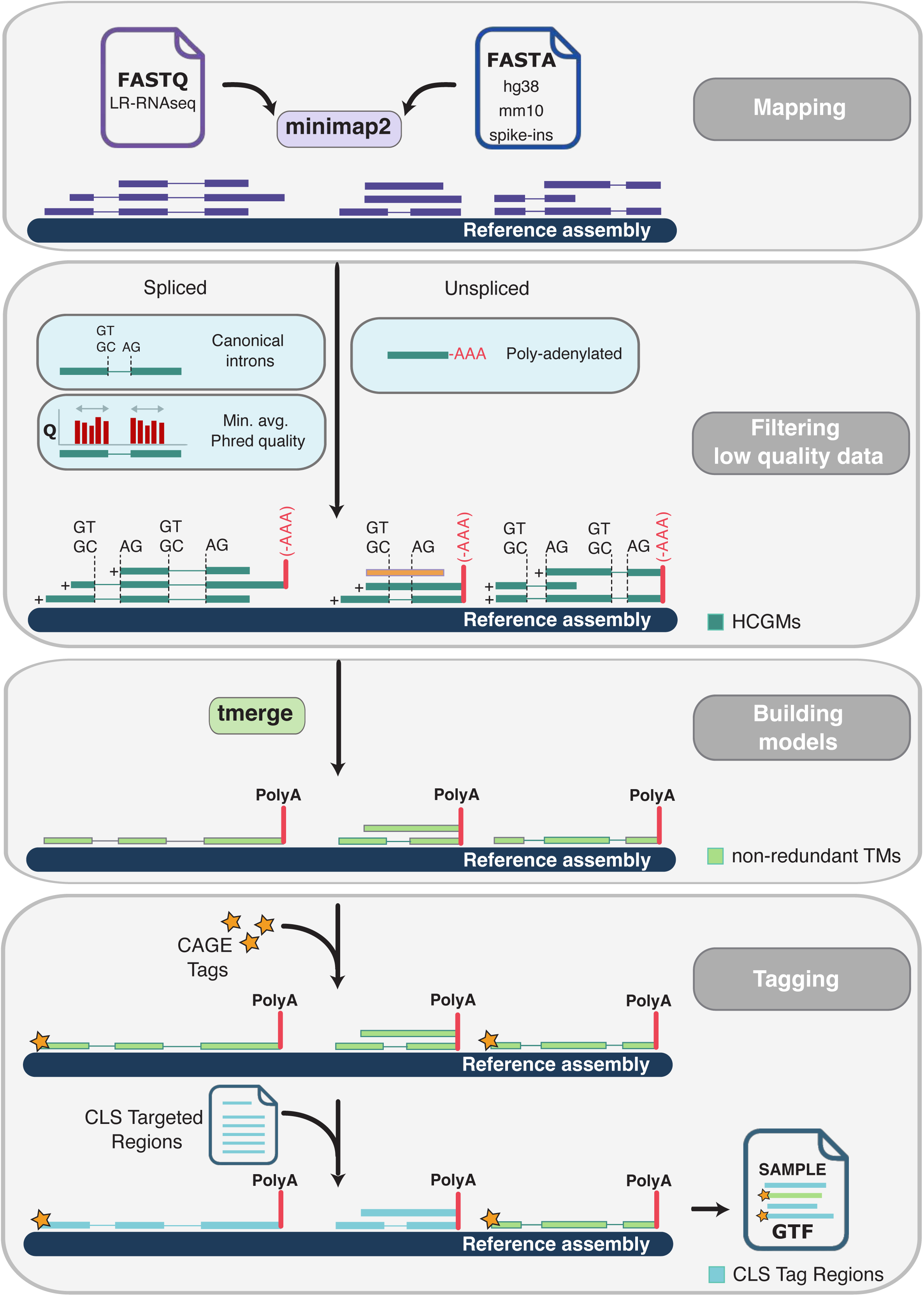
Overview of the LyRic pipeline for long-read RNA-Seq transcript model reconstruction. LyRic is a Snakemake-based pipeline designed to identify high-confidence, full-length transcript models from CapTrap-Seq long-read RNA-Seq data. The pipeline begins with the mapping of long reads to a reference genome using *minimap2*. Mapped reads undergo stringent quality control filtering to retain only High-Confidence Genome Mappings (HCGMs), defined as spliced alignments containing canonical splice-junctions or, in the case of monoexonic reads, the presence of a clipped poly(A) tail. Filtered high-confidence reads are processed separately based on splicing status. Compatible reads are merged using *tmerge* to generate non-redundant transcript models (TMs), while avoiding artifacts such as intron chain extension and correcting splice-junction overhangs. LyRic can tag the models using external and internal evidence. In this project, we have tagged CLS transcript models with CAGE tags, poly(A) tails and, in the case of post-capture samples, the target region from which the transcript likely originates. LyRic output is a GTF file containing full-length transcript models.

**Figure S27.**
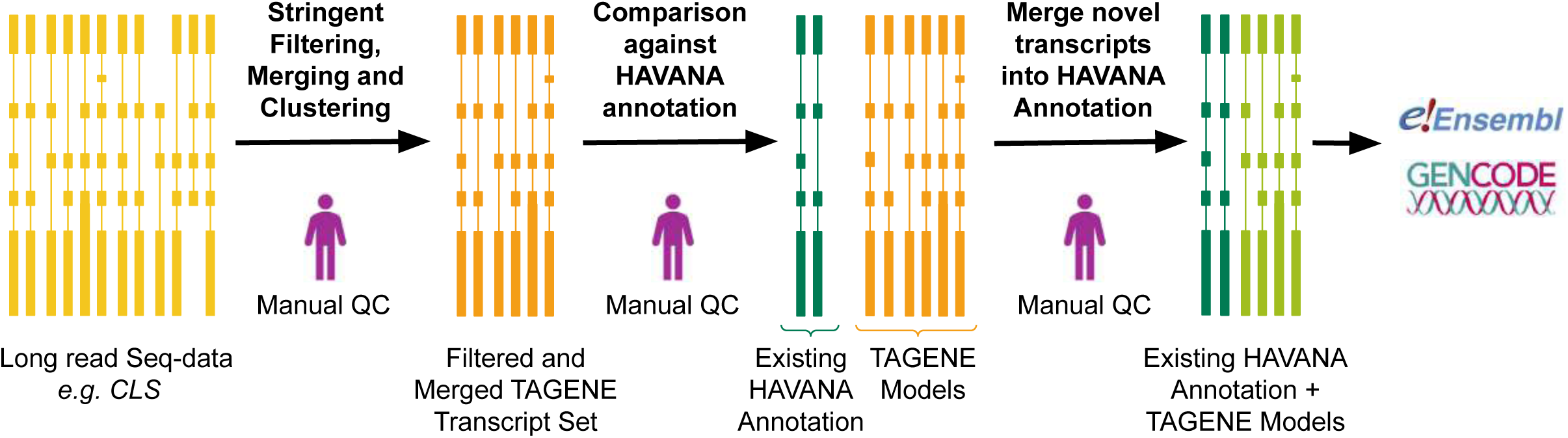
Overview of the TAGENE pipeline for high-stringency, large-scale annotation of lncRNAs using CLS long-read data. TAGENE is a semi-automated annotation pipeline developed to incorporate CLS long-read data into the GENCODE gene set under the stringent quality standards of reference annotation. It was designed to scale with large datasets and to mimic the principles of manual annotation performed by the HAVANA group, while focusing exclusively on lncRNA models. The workflow begins with aligned CLS reads from the LyRic pipeline and incorporates multiple filtering steps to ensure TAGENE could achieve near-manual stringency without requiring manual review of every model. TAGENE preserves annotation integrity by enforcing conservative rules for splice site inclusion, transcript boundary assignment, and merging behaviour, all tailored to the annotation of non-coding genes. Iterative input from GENCODE annotators informed the development, enabling the pipeline to serve as a scalable yet stringent framework for integrating long-read transcriptomics into the reference gene set.

