## Supplemental_TEXT for "The GENCODE CLS project: massively expanding the lncRNA catalog through capture long-read RNA sequencing"

### Glossary

‘**CLS**‘ Capture Long-read Sequencing, library preparation protocol in which probes against targeted regions of the genome are used to capture tailored transcript sequences.

‘**pre-capture**’ refers to the library preparation employing CapTrap protocol, and therefore extends to all the samples and models yielded from those.

‘**post-capture**’ refers to the library preparation employing CapTrap protocol in conjunction with CLS, and therefore extends to all the samples and models yielded from those.

‘**Raw reads’** refers to sequencing reads produced on the PacBio or ONT platforms.

‘**Aligned reads**’ are ‘raw reads' preprocessed and aligned to the genome using LyRic.

**‘LyRic models’** are transcript models generated by LyRic from the ‘aligned reads’ in each individual sample.

**‘CLS (anchored) models’** are models generated upon anchoring the ‘LyRic models’ to the 5’ and 3’ ends support, and then merging them across tissues, technologies and developmental stages.

‘**CLS transcripts**’ are intron chains resulting from merging ‘CLS models’, ignoring differences in transcripts termini.

**‘CLS loci**’ are ‘CLS models’ merged together to build regions of continuous transcription on the same strand, thus generating uniquely identifiable loci.

**‘Novel CLS transcripts’** are CLS transcripts not present in GENCODE v27 or vM16. They can overlap or not previously annotated transcripts.

‘**Novel lncRNA transcripts**’ are novel CLS transcripts now annotated in GENCODE v47 or vM36.

‘**Novel intergenic lncRNA (lincRNA) transcripts**’ are ‘novel lncRNA transcripts’ not overlapping any annotation as of GENCODE v27 or vM16.

### The GENCODE CLS project: massively expanding the lncRNA catalog through capture long-read RNA sequencing

#### GENCODE experimental protocols employed to survey the human and mouse transcriptomes

GENCODE has been at the forefront of using advanced methodologies to produce targeted transcriptomic data aimed specifically at gene and transcript annotation. The focus has been to survey the fraction of the transcriptome that is not usually well accessed by standard experimental protocols, complementing therefore the wealth of transcriptomic data produced by the scientific community and deposited in public data archives. The methodologies have evolved through the years towards the automatic production of high-quality full-length transcript sequences that can be incorporated in the GENCODE annotation with minimal manual curation efforts. During the pilot phase of ENCODE, extensive experimental validation of annotated transcripts was performed using 5' RACE and RT-PCR on multiple tissues, followed by Sanger sequencing[1]. With the development of tiling arrays, we used their multiplexing capacity to implement the RACEarray normalization strategy, by which 5' RACE products are hybridized into a tiling array[2]. Subsequently, RT-PCR is carried out from detected exons selected to correspond to novel, not previously annotated, transcripts[3]. Massively parallel sequencing superseded genome-wide tiling arrays, and we implemented RT-PCR-Seq, in which RT-PCR products from primers designed to validate exons are pooled and sequenced using Illumina[3]. We incorporated long-read sequencing technologies into the GENCODE experimental pipeline as soon as they became available. We developed RACE-Seq in which RACE products were pooled and sequenced using the ROCHE 454 FLX+ platform[4]. More recently, we implemented the Capture Long-read Sequencing (CLS) strategy, in which probes against targeted genome regions are used to capture transcript sequences, which are then sequenced using Pacific Biosciences long-read technology[5]. Over the years, GENCODE has produced, therefore, a unique, massive collection of tens of thousands of targeted transcriptome readouts that have contributed significantly to the quality of the annotation.

#### CapTrap-CLS capture efficiency

After mapping all reads to the 92 ERCC spike-ins, we observed a substantial fold increase for the 42 targeted, ranging consistently from 31 to 51-fold depending on the species, tissue and sequencing platform (from ~2.5% pre-capture to ~97% post-capture, on average, **Figure S1B**).

Mouse samples showed an overall higher read enrichment rate. In total, between 7% and 39% of all reads in humans and between 9% and 59% of all reads in mouse, depending on tissues and sequencing platforms, mapped to targeted regions in the post-capture samples. This leads to an on-target rate between 3.7 to 15-fold for human, and 7 to 36-fold in mouse (**Figure S1C**).

Overall, PacBio samples showed higher target enrichment rates in both species, despite a lower read throughput, from 6 to 15 in human, and from 9 to 36 in mouse. Similarly, the fold increase for ONT varied from 4 to 9, and from 7 to 30 (**Figure S1C**).

Embryonic, adult liver tissue and testis showed the highest enrichment rates for human in PacBio and ONT, while it was the case for adult liver and white blood followed by liver in mouse.

#### LyRic

To support the comprehensive and annotation-light reconstruction of full-length transcript models from the capture CapTrap-Seq long-read RNA-Seq data, we developed LyRic, a modular Snakemake-based bioinformatic pipeline[6]. LyRic is structured into four core components: **Mapping**, **Filtering**, **Building Models**, and **Tagging**, and incorporates quality control throughout the pipeline to ensure high-confidence results at every step (**Figure S26**). LyRic can process long-read RNA-Seq read data files individually, or if associated with a project, it can process them collectively, eventually producing a single set of transcript models across all input files. This is particularly relevant for the GENCODE annotation project. Here we describe the initial processing of individual files by LyRic. The collective post-processing, which has been employed in our work here, is described in the Methods section.

**Mapping:** Raw long-read RNA-Seq data (FASTQ) is aligned to the reference genome (e.g., hg38, mm10, or spike-in sequences) using *minimap2*[7]. These initial alignments form the foundation for downstream filtering and model building.

**Filtering:** LyRic identifies High-Confidence Genome Mappings (HCGMs) by selecting spliced alignments containing canonical splice-junctions (GT-AG, GC-AG) and meeting minimum average Phred quality score. For monoexonic reads, the presence of a clipped poly(A) tail is required to confirm transcript integrity. When short-read RNA-Seq data is available, LyRic optionally applies additional filtering to spliced HCGMs using this orthogonal short-read support to validate High-quality Short-read Supported (HiSS) models. In this study, short-read support was not used, and transcript models were generated exclusively from long-read HCGMs. LyRic's conservative filtering strategy balances high sensitivity with a low false positive rate, as demonstrated in the LRGASP benchmarking project[8].

**Building Models:** HCGMs (or HiSS reads, when applicable) are grouped and merged using *tmerge* to form non-redundant transcript models[9]. Spliced and monoexonic reads are processed independently to prevent inappropriate merging. *tmerge* also implements safeguards against intron chain overextension and corrects exon/intron overhangs at splice-junctions. Resulting models are further grouped into loci using *buildLoci*[10].

**Tagging:** In the final stage, LyRic can annotate transcript models using internal and external evidence, such as support by CAGE peaks and poly(A) sites. These are used, in our case, during the process of merging transcripts models across samples. In the case of post-capture samples, LyRic also annotates the probed region from which the transcript models likely originate.

#### Analysis of CLS models

The per-sample transcript models obtained using LyRic were "anchored" (see **Methods, Figure S2**) and merged together across all samples and technologies to produce a consolidated CLS annotation of 1,212,480 unique CLS models in human and 1,092,208 models in mouse. As an initial characterization of these models, we compared them to GENCODE v27 and vM16 annotations using SQANTI3[11], [https://guigolab.github.io/CLS3_GENCODE/SQANTI_reports/Human_CLStranscripts_v27.html, https://guigolab.github.io/CLS3_GENCODE/SQANTI_reports/Mouse_CLStranscripts_vM16.html ].

A larger fraction of models were built exclusively from ONT reads, 69% and 66% respectively in human and mouse, compared to 20% and 22% exclusively from PacBio reads, while about 11% of the models are detected by both technologies. 55% and 46% of the models were built exclusively from post-capture samples, compared to 37% and 44% pre-capture, with 8.5% and 9.6% of the models commonly produced between the two designs, in human and mouse respectively. The higher fraction of models detected in adult tissues with respect to embryo ones (64% and 67%, in human and mouse), is mainly attributable to the higher number of adult tissues employed.

The majority of the CLS models (about 95%) originate from one or two tissues only; models detected across sequencing technologies, developmental stages, and capture strategies are also more broadly detected across tissues (**Figure S8**). This relatively low overlap can be partially explained by the high redundancy of intron-chains retained upon anchored merging. Indeed, by anchoring 5' and 3' supported ends, we are intrinsically preserving highly similar transcript models with only slight differences at the termini.

##### Novel CLS models

To assess the novelty of the CLS models, we compared their structures to the GENCODE v27 and vM16 reference annotations using gffcompare[12], and assigned them to nine simplified categories based on the extent of their overlap, further grouped in two disjoint classes; known, and novel (**Table S3**). Of all CLS models, 262,843 were novel in human and 303,996 in mouse; the larger number of novel mouse models is likely attributable to the less advanced state of the annotation compared to human.

##### CLS transcripts

To further reduce the redundancy across the different models, we collapsed them into unique intron chains. We simply merged the CLS anchored models sharing the same internal exon-intron structure (as identified by gffcompare[12]), ignoring eventual variation at the terminal exons, and therefore neglecting end-support information.

A different strategy was employed for monoexonic models, which were extended into a single one when overlapping more than 50% of their length, again irrespective of the end support.

The separate projection of spliced and monoexonic transcripts resulted in a set of 468,598 unique intron chains and 57,709 monoexonic transcripts in human, and a total of 407,194 unique intron chains and 76,231 monoexonic transcripts in mouse. In the main manuscript and from this point onwards, we use the term "CLS transcripts" to refer to the union of unique intron chains and monoexonic transcripts (526,307 in human and 483,425 in mouse, **Figure 1CD**, **Figure S3**).

Different tissues produced different yields of CLS transcripts, with testis being among the most productive in both human and mouse, followed by brain (both in adult and embryo) in human and pool of tissues in mouse (**Figure S4A,B**).

Overall, 161,817 of all the CLS transcripts were novel in human (with respect to GENCODE v27) and 178,974 in mouse (with respect to vM16), of which 62,734 and 79,777 entirely intergenic (**Figure S3**, **Figure S4C,D**). The majority of these novel CLS transcripts were spliced (71% in human and 64% in mouse), and highly supported by independent recount3[13] reads; 80% of the CLS spliced transcripts in human and 91% in mouse have all the splice junctions supported by at least one recount3 read (**Figure S9A**). The transcripts obtained through PacBio have a higher recount3 support (90%, human; 96%, mouse) as compared to the ones obtained through ONT (73%, human; 86%, mouse), while as expected, the highest support was observed for the transcripts that are detected through both the technologies (94%, human; 98%, mouse) (**Figure S9A**). Further, the transcripts detected through both the pre- and post-capture strategies also showed a good recount3 support across all the tissues (> 93% in human; > 96.5% in mouse) (**Figure S9BC**). Overall, employing a threshold of 50 recount3 reads, 75% of all transcripts being supported in human, and 78.7% in mouse (**Figure S9DE**). We observed a high recount3 support even for the novel and completely intergenic transcripts.

For CLS transcripts, the extent of agreement between the two technologies increases, which makes the set of transcripts obtained through PacBio almost a subset of those gained through ONT (**Figure S8BC**). More precisely, in human, 66% of the CLS transcripts have been generated exclusively from ONT, while 25% are now shared between the two technologies. Similarly, 62% of the CLS transcripts have been generated exclusively from ONT in mouse, while 28% are shared.

The overlap also increases when comparing the transcripts detected pre- and post-capture; in human, 57% of the CLS transcripts are detected uniquely post-capture, while 24% are shared across designs. Similarly, 44% of the CLS transcripts are detected exclusively post-capture in mouse, and overall 24% are shared across designs (**Figure 1CD, Figure S3**).

The consensus across different tissues also increases, with about 20% of the CLS transcripts detected in three or more tissues, in accordance with the increase in consensus across sequencing technologies, developmental stages, and capture strategies (**Figure S8BC**).

##### Novel CLS Loci

Eventually, CLS transcripts sharing any overlap on the same strand were brought together to build 74,073 and 88,192 loci in human and mouse, respectively. An additional round of intersection with the reference annotation (GENCODE v27 for human, and GENCODE vM16 for mouse) was carried to track whether the newly identified loci overlaid previously annotated genes, and assign novelty at the loci level accordingly. We identified 36,815 novel CLS loci in human and 51,263 in mouse, respectively.

##### CLS models and target detection

Among all the regions targeted by this experiment, 37% in human and 36% in mouse eventually detected transcription (**Figure S5B**). We observed that the majority of the overall detected regions were detected uniquely post-capture (25.5% and 20.7% of all the regions, respectively in human and mouse).

Of all the targeted regions, 28.6% help detect 107,981 completely novel CLS transcripts (**Figure S5C, S6AC**), representing 66.7% of all novel CLS transcripts in human (**Figure S6E**). For mouse, 29.6% of the regions helped detect 105,399 completely novel CLS transcripts, representing 58.9% of all the novel ones (**Figures S5C, S6BD**), proving the effectiveness of the capture. Of these novel transcripts detected thanks to the targeted regions, 88% and 82% in human and mouse were detected only post-capture (**Figure S6EF**), highlighting the efficiency of the capture in enriching poorly annotated regions.

#### The TAGENE Pipeline

As a reference annotation resource, GENCODE has strict criteria for the inclusion of models into its gene set, and these must evolve when new technologies become available to support annotations. Over time, the sequencing quality of long-read technologies has improved, alongside methodological advances in the algorithms used to process and map the data. Nonetheless, GENCODE does not incorporate aligned transcriptomics data directly into the gene set, i.e., using an unsupervised, computational approach. This is because reference gene annotation strives towards perfection in terms of the quality of transcript models included. Thus, it is generally considered that the inclusion of incorrect models, e.g., those that do not accurately represent genuine biological transcripts, is more harmful to the gene set than the omission of correct models. In other words, in this context, reference annotation takes a conservative approach.

Historically, the accuracy of the reference annotation has been ensured by the fact that all GENCODE models are constructed manually by expert human annotators, who examine the ‘evidence’ for each prospective model (e.g., transcriptomics data) in the context of the genome sequence. For the first decade of the GENCODE project, human and mouse annotations were built almost entirely using cDNAs and EST sequences deposited in INSDC databases, the number of which proved to be tractable for a purely manual approach, which was deployed chromosome-by-chromosome, locus-by-locus.

Today, the primary challenge provided by long-read datasets to reference gene annotation is in terms of the sheer volume of reads produced. These numbers are not tractable for GENCODE manual annotation, as previously performed. However, we consider that full manual annotation is not necessary for long-read datasets. Primarily, this is because the size of next-generation datasets, including short-read RNA-Seq libraries, provides key information leveraged to support the annotation process. Fundamentally, the most important aspect is confidence in the structure of a transcript model, i.e., its splice-junctions, start, and end points. Historically, the need to individually check each of these elements of a prospective model was a major time burden on the manual annotation process. It is now well established that splice-junction accuracy can be controlled, at least to some extent, in model construction through the use of short-read data to supplement long-read alignments. However, before our work here, it was not apparent how this should be achieved to create reference gene annotations.

Thus, to maintain reference annotation standards for the incorporation of CLS data, it was first necessary to fully understand the quality of the aligned reads produced by LyRic. At the same time, given the size of these datasets, we needed to devise a pathway for the large-scale incorporation of these reads into GENCODE that was not dependent on manual annotation, and yet nonetheless could achieve high stringency. The combination of these two strands evolved into creating the TAGENE annotation workflow (**Figure S27**).

TAGENE, as far as possible, has been designed to replicate the manual annotation process in terms of its logistical stages. However, the process is greatly simplified as we are not attempting to annotate coding sequences, given that the scope of this phase of the project is entirely focused on the annotation of lncRNAs. Thus, in creating GENCODE gene models, TAGENE needs to accomplish several things: *i)* ensure that models have accurate splice-junctions, *ii)* set appropriate first and last coordinates for each model, i.e., start and end points, which would ideally correspond to transcription start sites (TSSs) and polyadenylation sites as discussed below, and *iii*) ensure that 'merging' behavior is appropriate, i.e., the process by which two or more overlapping models may or may not be ultimately joined into a single GENCODE model. To ensure that TAGENE works in line with the principles of manual annotation, our solution was to develop an iterative workflow based on substantial input from the expert annotators in the HAVANA group, which were fed back to the development team to make adjustments for the next run.

Splice-junction accuracy is crucial to the overall quality and consistency of the GENCODE gene set; we take a conservative approach to the assessment of introns for inclusion, where only canonical splice sites supported by contiguous alignment of transcriptional data are allowed. Thus, the HAVANA group at EMBL-EBI performed a manual assessment of the quality of splice-junctions contained in CLS reads aligned through LyRic. At this point, we were specifically interested in examining introns that had low expression, considering that this set was more likely to include false splicing reactions. For all aligned reads produced by LyRic, we first produced scores for all introns. Specifically, we measured the level of expression of each splicing reaction (as opposed to the expression of the overall transcript) using short-read data processed by the recount3 project. Next, 10 aligned read introns for each recount3 intron score between 0 and 199 were randomly selected to create a set of 2000 introns for manual checking. Each intron was assessed for accuracy based on standard GENCODE annotation guidelines, and independently of any other evidence. Introns identified as being incorrect, i.e., not annotatable by GENCODE, were QC'd a second time by a second annotator. Where annotators were satisfied that an intron was incorrect, we attempted to extrapolate a reason for this outcome. This allowed for us to classify false introns into several categories, outlined below, and then used to devise additional filtering stages for TAGENE. To ensure the high stringency of reference annotation reads were screened for *i)* recount3 score, *ii)* tandem repeats overlap, *iii*) pseudogenes overlap, *iv)* opposite strand mismapping to coding genes, and *v)* splice sites misalignments.

Of the 165 incorrectly spliced introns identified, 115 had a recount3 score below 50. The majority fell into one of the "incorrect alignment" categories previously defined. It was therefore decided to set a conservative recount3 intron score threshold of 50, attempting to remove the majority of the observed alignment errors. We estimate that approximately 50% of introns with a score below 50 are, in fact, likely correct, although this proportion is inappropriately low to support reference annotation. Further, we identified 14 examples in which the splice-junction of a CLS read was localized within a genomic tandem repeat, as defined by Tandem Repeats Finder. In short, tandem repeats are defined as two or more adjacent copies of a sequence of nucleotides. In these cases, we found that it was impossible to unambiguously determine the true splicing structure of the CLS read, as it aligned equally well to multiple locations. As such, we decided to filter out all models that contain a splice-junction localized within a tandem repeat. In 50 cases, CLS reads incorrectly aligned at or near a pre-existing GENCODE processed pseudogene due to the alignment of the read poly(A) tail to the genomic poly(A) sequence inserted at retrotransposition, ultimately conferring a higher mapping score to the pseudogene locus over the parent protein-coding gene. For this reason, we decided to filter out all models having same-strand exonic overlap with annotated pseudogenes. We note that this was a brute-force approach, and that substantial numbers of models overlapping pseudogenes are likely genuine. We plan to revisit this question in future iterations. Furthermore, we also found 20 examples in which LyRic misaligned a CLS read onto the opposite strand of a protein-coding gene. It was determined that these reads genuinely belonged to the protein-coding gene, but were incorrectly placed due to the pipeline's alignment software being unable to find canonical splice sites on the true strand. This was in turn caused by the poor sequence quality of the underlying reads, and it was therefore decided to filter out all models antisense to annotated genes, when the first and final coordinate of such models fell within a window defined by the gene boundaries of the existing locus. Finally, 57 cases were identified where the lack of sequence similarity between the CLS read and the genomic sequence, particularly around potential splice sites, led to ambiguous alignments deemed by a GENCODE annotator to be resolvable to an alternative, better supported, splice site. In these cases, we suspect that the underlying RNA-Seq support from the recount3 dataset likely fell victim to the same misalignment issue.

#### Merging CapTrap-Seq CLS transcripts into the existing GENCODE annotation

The GENCODE gene set (v46) already contained a rich and highly detailed catalog of lncRNA annotations (20,310 genes, 59,927 transcripts). The new CapTrap-Seq CLS transcripts therefore needed to be integrated into this existing annotation logically and consistently, and following the current GENCODE manual annotation guidelines. All novel CLS transcripts sharing exonic overlap and mapping to an intergenic region were assigned to the same new lncRNA locus and provided with a unique, stable Ensembl locus-level ID (ENSG ID) and Ensembl transcript-level IDs (ENST IDs). CLS transcripts with exonic overlap to a single pre-existing GENCODE lncRNA were automatically designated as part of the existing locus and given new Ensembl transcript-level stable IDs. Where CLS transcripts shared exonic overlap with two or more pre-existing lncRNA loci, GENCODE expert human annotators provided manual oversight to determine whether they should be merged into a single locus or maintained as separate genes. If the loci were merged, the CLS transcripts were automatically added to the new single gene; otherwise, manual supervision was required. The decision-making was carried out under the current GENCODE manual annotation guidelines, evaluating the presence of transcriptional data to support TSSs, poly(A) sites, and consistent use of the proposed full-length intron chains as determined via RNA-Seq intron counts from the recount3 dataset. The maintenance of previously well-defined lncRNA genes was prioritized by manually supervising the addition of CLS transcripts to lncRNAs with known gene names/symbols. Additionally, we consulted the HUGO Gene Nomenclature Committee (HGNC) at the University of Cambridge regarding all cases in which lncRNAs with known gene names/symbols were to be merged, to determine which gene name/symbol should be retained.

#### Protein-coding genes

Although this study focuses on lncRNAs, ~84,000 novel human and ~98,000 mouse CLS models map to ~12,000 and ~14,000 known protein-coding genes, respectively. This data has so far led to the inclusion of ~9,500 very high stringency protein-coding transcripts in GENCODE v49, with many more transcripts likely to be added in future.

At the same time, we have investigated the possibility that our CLS data could uncover previously unknown protein-coding genes. Thus, we carried out an initial study using two methods. First, we interrogated large-scale tissue-based proteomics experiments for evidence of translation. We found convincing results for seven human novel protein-coding genes with multiple peptides not annotated as coding in the Ensembl/GENCODE reference set. All seven proteins have known human paralogs, and four are substantially truncated compared to the parent gene (**Figure S15**, structures predicted using the HHPRED server[14] or AlphaFold3[15]). Six proteins were detected principally or wholly in testis. For example, *C5orf60* was annotated as non-coding in both Ensembl/GENCODE and RefSeq, but we detected ten peptides for the protein in our analysis. We detected peptides for the correct *WASHC1* gene that has recently been uncovered as part of the novel T2T-CHM13 assembly[16,17]. For mouse, analysis of proteomics data led to the discovery of 23 protein-coding genes that were novel to GENCODE, most of which were also expressed in testis (**Figure S15**). The coding status of many of these ORFs was already predicted by other projects (mostly RefSeq in the case of mouse), with GENCODE having previously considered many of them to be pseudogenes.

Second, we used PhyloCSF[18] to find open reading frames (ORFs) within CLS transcripts most likely to contain evolutionarily conserved novel protein-coding regions. Manual examination of over 800 top candidates in human and mouse identified one novel protein-coding gene in each (**Figure S14**). The human one has a 24 codon ORF having hg38 coordinates chr3:117729172-117729179+chr3:117997182-117997248(-). The mouse one has two isoforms, a 37 codon ORF having mm10 coordinates chr6:136171799-136171878+chr6:136172263-136172296(-), and a 16 codon ORF having coordinates chr6:136171861-136171882+chr6:136172268-136172296(-). Each of these novel genes has a novel ortholog in the other species, but it is not contained in any CLS transcript. Neither of these PhyloCSF-supported ORFs overlapped with any of the novel coding genes for which we detected peptide evidence.

#### Pseudogenes

Similarly to the case of protein-coding genes, although this study was specifically focused on lncRNAs, a number of probed regions overlapped pseudogenes (**Figure S5A**). In total 5,071 pseudogenes in human, and 2,280 in mouse were targeted in this study **(Table S7.1**). Upon re-quantification of CLS loci across all tissues and cell lines, 98.75% of alignments were successfully assigned to a feature. As expected, given that pseudogenes are generally transcribed at low levels, our capture strategy enhanced the sensitivity in recording expression changes in pseudogenes and their parent genes. For both human and mouse, we observed that more pseudogenes were upregulated using ONT compared to PacBio. In contrast, more parent genes showed upregulation with PacBio (**Figure S16A**).

Among upregulated pseudogene-parent gene pairs, 1,250 (61%) showed increased expression post-capture in human (**Figures S16B**) and 790 (71%) in mouse. Capturing had a larger impact on the quantification of expression of pseudogenes than of parent genes. Nearly half of the parent genes were not upregulated even when their corresponding pseudogenes were upregulated (**Table S7.2**, **Figure S16C**). Still, capturing had an impact on the quantification of parent genes. Over 30% of parent genes were upregulated in human (33% in mouse) after capturing, compared to only 18% (20% in mouse) of non-parent protein-coding genes. These, in contrast, were largely downregulated (**Figure S16DE, Table S7.3**). The contrasting behavior between parent and non-parent genes is expected, as protein-coding genes were not targeted in our design, but parent genes share strong sequence similarity with pseudogenes, and could be captured by our design.

#### Support of CLS models by Histone modifications

We utilized the ENCODE4 cCRE tissue-agnostic catalog to evaluate the extent to which the transcription start sites (TSSs) of our novel CLS models are supported by external epigenetic evidence. In this context, cCREs support is defined by a proximity of less than 2kb between a given TSS and the center of a cCRE. While known TSSs were mostly supported by proximal cCREs (PLS, and pELS), novel TSSs were more supported by distal cCREs (dELS), largely reflecting the fact that the ENCODE cCREs classification depends on the proximity to TSSs (**Figure 3B**).

To investigate how this enrichment correlates with the expression levels of these novel TSSs across the different tissues, we classified novel TSSs into two main groups: ubiquitously and non-ubiquitously expressed. Within the non-ubiquitously expressed group, we further identified tissue-specific TSSs. For each TSS, we gathered all the corresponding CLS models and recorded the tissues in which they were expressed, without distinguishing between adult and embryonic samples. A TSS is classified as ubiquitously expressed if its associated CLS models are expressed across all tissues. If the CLS models are expressed in only a subset of tissues, the TSS is classified as non-ubiquitously expressed. Within this latter case, if all CLS models associated with a given TSS are expressed in a single tissue, the TSS is defined as tissue-specific. From this latter set, TSSs uniquely expressed in tissue-pool or cell-pool samples were excluded, as it is not possible to determine which specific tissue or cell-type contributed to their detection. A similar proportion of ubiquitously expressed and non-ubiquitously expressed TSSs are supported by cCREs (96% and 95%, respectively). However, while the former are mostly associated with PLS and pELS, the latter are predominantly supported by dELS (**Figure S17A**). This enrichment in distal regulatory activity was even stronger among tissue-specific TSSs: 69% on average across tissues, show support by at least one dELS (**Figure S17A**).

We also evaluated whether the dELS cCREs intersecting tissue-specific TSSs showed activity in the same tissue as the corresponding CLS model. For this analysis, we focused on three adult tissues (heart, liver, and testis) with available tissue-specific cCRE catalogs and maps of H3K27ac and H3K4me3 histone modifications[19]. On average across the three tissues, 11% of the intersecting dELS showed activity (i.e., were not classified as “low DNase”) in the same tissue as the corresponding TSS. This proportion reached an average of 49% (53% heart, 63% liver, and 30% testis) after integrating peaks of H3K27ac and H3K4me3 histone modifications (**Figure S17B**).

#### Lifting over the GENCODE human lncRNA set to primate genomes and comparison with human-mouse orthology

The 32 assemblies used in this analysis included representatives from great apes, lesser apes, old and new world monkeys, as well as lemurs and lorises. All assemblies were at chromosome-level resolution, except for *Papio cynocephalus* and *Saimiri sciureus*, which are at scaffold-level, explaining the lower number of lifted features, especially protein-coding genes, compared to other species (**Figure 4B**). Among these 32 primates, 23 had RefSeq annotations, which were enhanced by incorporating lifted transcripts from GENCODE v47. As a control, lifted transcripts were classified into 9 groups according to existing annotations using gffcompare (**Figure S21CD**). Approximately 80% of protein-coding genes overlapped existing features, 15% were considered equal, and 4% as extensions. For lncRNA, three groups were overrepresented, with 42% classified as intergenic (62% for intergenic lncRNA transcripts), 30% as overlapping (10% for intergenic lncRNA transcripts), and 19% as antisense. Note that the number of certain gene biotypes may be lower than in the initial reference due to edge cases where merging different biotypes was required, particularly lncRNA and protein-coding genes. Finally, publicly available short-read RNA-Seq data were used to quantify gene expression for the enhanced annotation. RNA-Seq data were available for at least one tissue in 29 species and for more than three tissues in 21 species (**Table S15**). Note that species with a high number of features exhibit tissues associated with the nervous system, particularly the brain.

We compared Liftoff, a transcript-structure-aware annotation mapping tool, used to map the human lncRNA onto the primate genomes with ConnectOR, a synteny-based ortholog identification tool, used for the human mouse orthology analysis. (**Figure S22A**). ConnectOR identifies lncRNA orthologues by leveraging genome synteny and using a reciprocal, exon-based liftover strategy. Each exon is treated individually, and an orthology relationship is established if at least one exon from a gene is successfully lifted over in both directions (human-to-mouse and mouse-to-human) with a minimum of 30% overlap (minMatch = 0.3). This stringent, bidirectional approach ensures the identification of conserved loci and excludes species-specific predictions. Liftoff, in contrast, transfers transcript annotations from a well-annotated reference genome (e.g., human) to a target genome (e.g., mouse), based on sequence similarity and preservation of exon–intron structure. It operates in a one-directional manner and is designed to support annotation propagation rather than establish evolutionary relationships.

The Connector approach benefits strongly from our experimental approach, where orthologous regions in human and mouse are targeted and the RNA sequencing is performed in equivalent tissues.

We evaluated the overlap between the two methods and found that only 17% of human lncRNAs with mouse orthologs identified by ConnectOR were also recovered by Liftoff, whereas 99% of Liftoff-mapped genes were shared with ConnectOR (**Figure S22B**). This asymmetry underscores the broader sensitivity of ConnectOR compared to the more selective nature of Liftoff. Despite these methodological differences, a subset of genes was consistently identified by both tools. For example, both methods mapped the human lncRNA gene *GPR158-AS1* to *Gm13327* in the mouse genome (**Figure S22C**), a relationship also supported by independent studies[20]. Furthermore, Liftoff predicted multiple *GPR158-AS1* transcript mappings in the *Callithrix* genome, substantially extending the annotation of *LOC118155277*. It also projected corresponding transcripts in the remaining 31 primate species lacking initial annotations, potentially expanding their annotations.

Among the human–mouse lncRNA gene mappings, 89% (n = 1,431) of the Liftoff-mapped genes were also present in the projections across seven clades representing 32 primate species. In contrast, ConnectOR identified a smaller overlap (62%), which increased to 94% when the most evolutionarily divergent primate group (lemurs and lorises) was excluded.

In conclusion, this comparative analysis illustrates the methodological differences between the tools: ConnectOR is designed to infer conserved orthologs based on individual exon-level synteny and strict bidirectionality, whereas Liftoff focuses on preserving transcript structure to extend gene annotations. While Liftoff predictions may, in some cases, indicate potential orthology, such cases should be interpreted cautiously and ideally validated, alongside assessing the functional potential of ConnectOR predicted lncRNA orthologs.
