## Supplementary material for "The GENCODE CLS project: massively expanding the lncRNA catalog through capture long-read RNA sequencing": TABLES_captions

### Supplementary Tables Captions:

**Table S1.** Number and total coverage of the targeted regions in the capture panel.

**Table S2.** Metadata of RNA samples used in the study, including sample identifiers and associated information. Each Sample ID follows the format <Origin>_<Reference>_<Sample Description>_<Lot Number>. The table includes the origin (supplier or institution), species (HS: Homo sapiens; MM: Mus musculus), and tissue or cell-type. Several samples represent pooled tissues or cell lines as specified.

**Table S3.** Novelty tags assigned to CLS transcripts based on the different gffcompare classes.

**Table S4.** Overlap of TSS from protein-coding genes, lncRNAs and novel CLS models with repetitive regions within the human genome.

**Table S5.** Counts of current GENCODE genes and transcripts created using CLS data **1)** since human releases v27 and v46 and **2)** mouse releases vM16 and vM35.

**Table S6.** Fraction of the genome covered by annotated exons in different GENCODE versions, for human and mouse, reporting the fraction of total genomic area covered.

**Table S7. 1)** Number of upregulated pseudogenes and parent genes in human and mouse, based on whether they are targeted or untargeted. **2)** Number of pseudogene-parent gene pairs in human and mouse, grouped by upregulation status in pre-capture, post-capture, or non-significant categories. **3)** Number of differentially expressed pseudogenes and parent genes in human and mouse, based on different sequencing technologies and capturing approaches. **4)** Pseudogene-parent pairs in human and **5)** mouse.

**Table S8.** Overlap (number and fraction) of ChIP-Atlas TF peaks with TSSs of protein-coding transcripts, lncRNAs and novel CLS models.

**Table S9.** PhyloP score distributions from 241 species Zoonomia Cactus alignment for categories of GENCODE transcripts. The columns show the per-transcript counts with the frequencies of scores categorized as accelerated, neutral, or conserved. The datasets are the decoy models, and GENCODE v27 or v47 transcripts. The GENCODE transcripts are divided into subsets based on various attributes. These protein-coding mRNAs, existing lncRNAs in v27, and novel lincRNAs in v47. 'PC loci" indicate lncRNAs that overlap with protein-coding genes, and 'non-PC loci' are outside protein-coding genes. The "host" category are transcripts or unique introns that overlap miRNAs or snoRNAs on the same strand, while "non-host" have no or opposite strand overlap of these small RNAs.

**Table S10.** miRNAs count and nucleotide span in **1)** human and **2)** mouse protein-coding genes, lncRNAs and novel CLS loci.

**Table S11.** Expression patterns of the MIR513 cluster novel host gene upon remapping of the RNA Tissue Atlas with GENCODE v47. A threshold of 0.25 TPM was considered significant and is shown in green.

**Table S12.** Small RNAs in annotated and novel genic regions in **1)** human and **2)** mouse protein-coding genes, lncRNAs and novel intergenic CLS loci.

**Table S13.** Summary table of reference genome assemblies and taxonomic information for 32 selected primates. The table lists taxonomic, genomic, and assembly metadata for each species included in the study. Columns include: common and scientific names, NCBI taxonomy ID, and lineage classifications (genus, family, order). Divergence time from *Homo sapiens* is given in millions of years where applicable. Assembly metadata includes RefSeq accession numbers, assembly names, release dates, FTP links to genome FASTA files, and annotation report URLs. For short-read RNA-Seq, information on the number of available tissues and sequencing depth (selected and total reads) are also reported when available.

**Table S14**. Number of protein-coding and lncRNAs annotated across primate genomes.

**Table S15.** Short-read RNA-Seq samples used in the lncRNA validation in primate genomes.

**Table S16.** Differentially expressed genes across different brain areas and disorders. The largest absolute value of logFC and the most significant q-value are reported for each entry. The column *integration_disease* reports whether the gene was significant upon inverse normal integration across different studies for the same disease. Similarly, *meta_analysis* is TRUE if the gene was significant upon inverse normal integration across all the studies under investigation.

**Table S17. 1)** Summary of the clusters generated upon requantification of the single-cell and single-nuclei experiments collected in the Human Commons Cell Atlas. The status reports whether they are identical (PAIRED), NOVEL or a FUSION, with respect to the clustering solution upon removal of novel lincRNAs. The column *lincRNA_marker* displays whether a novel lincRNA marker was significant for the cluster; the same upon stringent filtering (see Methods) is reported in column *filter_lincRNA_marker*. Summary statistics are reported on the right. **2)** List of novel lincRNAs evaluated in this study; the columns report, from left to right, *i)* the gene ID, whether it has been detected as a marker in *ii)* any cluster, or *iii)* in a novel cluster. Summary statistics are reported on the right.
